# txci-ATAC-seq, a massive-scale single-cell technique to profile chromatin accessibility

**DOI:** 10.1101/2023.05.11.540245

**Authors:** Hao Zhang, Ryan M. Mulqueen, Natalie Iannuzo, Dominique O. Farrera, Francesca Polverino, James J. Galligan, Julie G. Ledford, Andrew C. Adey, Darren A. Cusanovich

**Affiliations:** University of Arizona, Department of Cellular and Molecular Medicine, Tucson, AZ; Oregon Health & Science University, Department of Molecular and Medical Genetics, Portland, OR; University of Arizona, Department of Pharmacology and Toxicology, Tucson, AZ; University of Arizona, Asthma & Airway Disease Research Center, Tucson, AZ; University of Arizona, Division of Pulmonary, Allergy, Critical Care & Sleep Medicine, Tucson, AZ; Banner - University Medicine North, Pulmonary - Clinic F, Tucson, AZ; Oregon Health & Science University, Cancer Early Detection Advanced Research Center, Portland, OR; Oregon Health & Science University, Knight Cancer Institute, Portland, OR; Oregon Health & Science University, Knight Cardiovascular Institute, Portland, OR

**Keywords:** Single-cell, ATAC-seq, Chromatin accessibility, Combinatorial indexing, Molecular hashing, CC16

## Abstract

Measuring chromatin accessibility is a powerful method to identify cell types and states. Performed at single-cell resolution, this assay has generated catalogs of genome-wide DNA regulatory sites, whole-organism cell atlases, and dynamic chromatin reorganization through development. However, the limited throughput of current single-cell approaches poses a challenge for implementing proper study designs, population-scale profiling, and/or very deep profiling of complex samples. To this end, we developed a 10X-compatible combinatorial indexing ATAC sequencing (“txci-ATAC-seq”), which is a combinatorial indexing framework that initially indexes (“pre-indexes”) chromatin within nuclei with barcoded transposases followed by encapsulation and further barcoding using a commercialized droplet-based microfluidics platform (10X Genomics). Leveraging this molecular hashing strategy, we demonstrate that txci-ATAC-seq enables the indexing of up to 200,000 nuclei across multiple samples in a single emulsion reaction, representing a ∼22-fold increase in throughput compared to the standard workflow at the same collision rate. To improve the efficiency of this new technique, we further developed a faster version of the protocol (“Fast-txci-ATAC-seq”) that separates sample pre-processing from library generation and has the potential to profile up to 96 samples simultaneously. We initially benchmarked our assay by generating chromatin accessibility profiles for 230,018 cells from five native tissues across three experiments, including human cortex (28,513 cells), mouse brain (48,997 cells), human lung (15,799 cells), mouse lung (73,280 cells), and mouse liver (63,429 cells). We also applied our method to a club cell secretory protein knockout (CC16^-/-^) mouse model to examine the biological and technical limitations of the mouse line. By characterizing DNA regulatory landscapes in 76,498 wild-type and 77,638 CC16*^-/-^* murine lung nuclei, our investigations uncovered previously unappreciated residual genetic deviations from the reference strain that resulted from the method of gene targeting, which employed embryonic stem cells from the 129 strain. We found that these genetic remnants from the 129 strain led to profound cell-type-specific changes in chromatin accessibility in regulatory elements near a host of genes. Collectively, we defined single-cell chromatin signatures in 384,154 nuclei from 13 primary samples across different species, organs, biological replicates, and genetic backgrounds, establishing txci-ATAC-seq as a robust, high-quality, and highly multiplexable single-cell assay for large-scale chromatin studies.

## Background

Chromatin accessibility measurement has become a widely used method to understand gene regulation and identify cell types and states. A common technique is the “assay for transposase-accessible chromatin using sequencing” (ATAC-seq) [1], in which a hyperactive mutant of the Tn5 transposase inserts sequencing adapters into sterically open (‘accessible’) regions of chromatin. After mapping the locations of these insertions, the resulting pile-up of genome-aligned reads identifies loci that are putatively active in gene regulation [1]. Performed at single-cell resolution (scATAC-seq), this assay has generated catalogs of genome-wide DNA regulatory sites, dynamic chromatin reorganization through development [2], and whole organism cell atlases [2,3].

Most modern single-cell methods generate data on hundreds to thousands of cells in parallel to enable proper characterization of heterogeneous or dynamic cellular systems. Two general strategies have been developed to generate data at this scale. First, cells can be isolated into individual reaction vessels - plate wells, microwells, or droplets. This has most commonly been implemented with microfluidics platforms, such as the commercialized products of 10X Genomics [4]. Second, iterative split-pool barcoding, as is seen in “single-cell combinatorial indexing” (sci) strategies, can index single cells while never isolating individual cells during the molecular reactions [5–7]. However, choosing one of these two approaches requires researchers to accept tradeoffs in terms of throughput and data quality. Microfluidic approaches generally have superior data quality, while combinatorial indexing benefits from flexibility, increased scalability, and cost efficiencies.

One strategy to boost the scalability of microfluidic approaches has been to “pre-index” cells or nuclei before loading them on a microfluidic device. In this way, aliquots of cells/nuclei are provided with a specific cellular/nuclear barcode via one of a variety of strategies and then aliquots are pooled before loading on a microfluidic device. The pre-index can be used along with the droplet barcode to deconvolute individual cells at the data analysis stage. This allows multiple samples to be processed in parallel and can enable some “overloading” of the droplets. For example, a single nucleus barcoding approach (SnuBar) [8] was previously demonstrated to allow for pre-indexing of nuclei in a scATAC-seq approach. However, individual molecules are not labeled in this strategy and thus droplets with multiple nuclei could not be discriminated, somewhat limiting the overall throughput. In another approach, a chimeric single-cell method combining a droplet-microfluidic system with molecular-level pre-indexing (called “dsciATAC-seq”) was previously developed, which improved the throughput of the microfluidic platform without sacrificing the data quality [9]. In this case, because the pre-indexing occurs at the molecular level (rather than the nuclear level), droplets containing multiple nuclei can still be computationally deconvoluted. However, this large-scale, single-cell approach was not developed for the 10X platform, which is more widely used for single-cell data generation. Here we demonstrate a method that takes advantage of the benefits of both combinatorial indexing and microfluidic assays by combining 96-well plate-indexed tagmentation with 10X Gel Bead-In EMulsions (GEM) encapsulation to substantially improve the throughput of the 10X platform by overloading nuclei and enabling the multiplexing of up to 96 samples in a single reaction (Fig. 1a). We call this method 10X-compatible (or <u>T</u>en<u>X</u>-compatible) <u>C</u>ombinatorial Indexing ATAC-seq (txci-ATAC-seq). We use this strategy to generate up to 200,000 cells in a single 10X reaction (∼22-fold increase in cell throughput as compared to the standard 10X Chromium scATAC-seq at a constant collision rate) and apply it to study the heterogeneity of chromatin accessibility in five primary samples, including human and mouse brain, human and mouse lung, and mouse liver, demonstrating the robustness of this approach. The scalability and flexibility of txci-ATAC-seq make it suitable for single-cell atlas efforts, population-scale studies, and experiments implementing replicates and proper study design.

**Figure 1.**
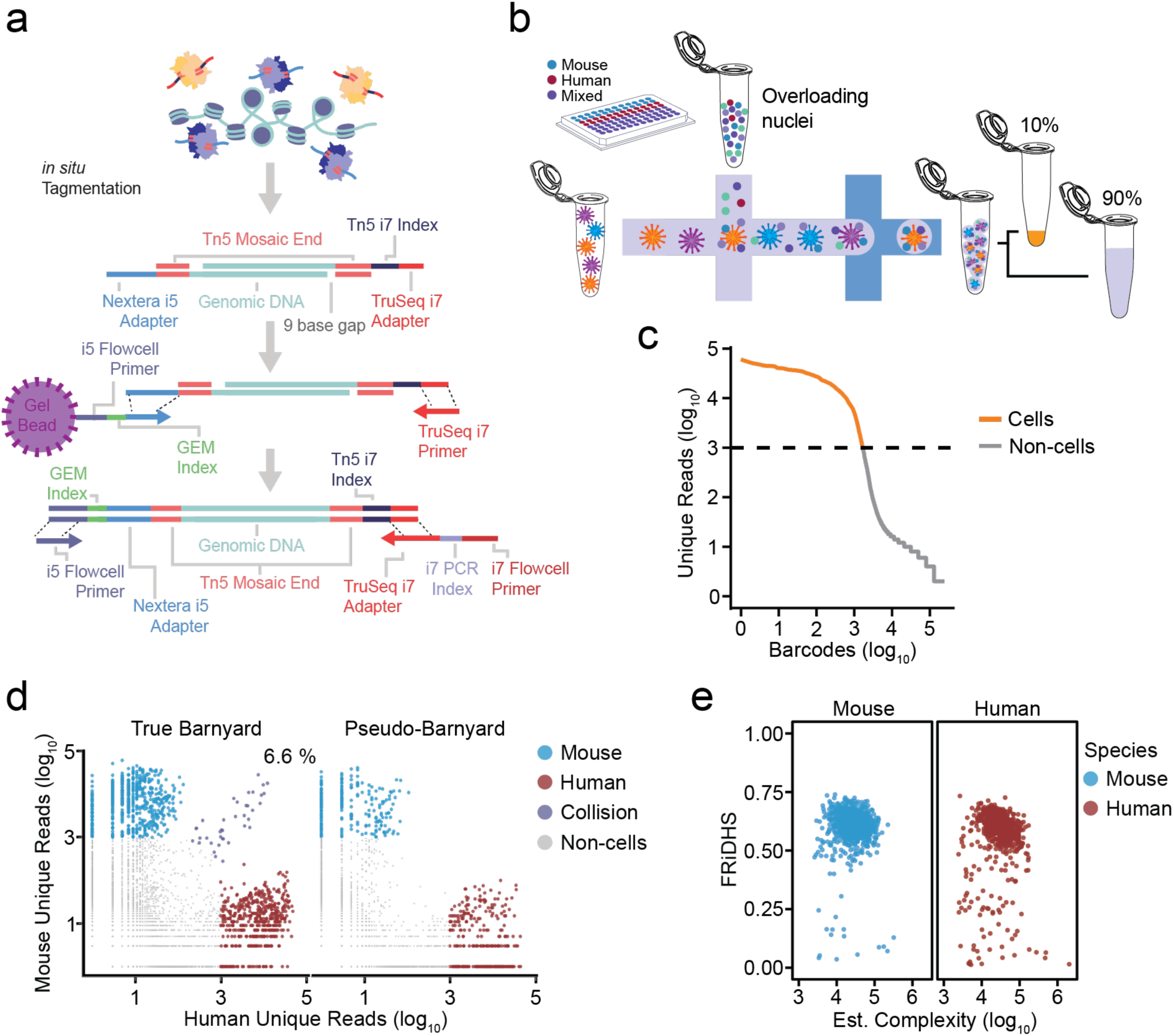
txci-ATAC-seq generates high-quality single-cell ATAC libraries at high throughput. a) Schematic of molecular details of txci-ATAC-seq library generation. b) Experimental workflow for txci-ATAC-seq barnyard library generation. After 96-plex tagmentation, nuclei are overloaded on a 10X Chromium microfluidics device. Following nucleus encapsulation in the formed droplets, 10% of the GEMs can be used for quality control and the remaining 90% for data analysis. c-e) txci-ATAC-seq QC metrics for human (GM12878) and mouse (CH12) cell lines supplemented with SBS primer during in-droplet PCR. c) “Knee” plot showing the unique reads (log_10_ scale) against the rank of each barcode (log_10_ scale) ordered from most unique reads (left) to least (right). The dashed line indicates the threshold (1000 reads) used to identify cell barcodes (orange points). d) Scatter plots showing the number of unique reads mapped to either the human or mouse genome for both true and pseudo-barnyard experiments. Values were log_10_-transformed after adding a pseudo-count of 1 to all values. The percentage shown in the true barnyard panel (6.6%) represents the estimated collision rate. e) Scatter plots showing the FRiDHS against the estimated complexity for each cell barcode detected as either mouse (blue) or human (red) cell. The estimated complexity is shown on a log_10_ scale.

## Results

### Coupling droplet-based microfluidics with indexed transposition enables the overloading of nuclei

In order to implement a strategy analogous to dsciATAC-seq [9] on the 10X platform, we first conducted a pilot experiment, tagmenting nuclei using 96 barcoded Tn5 reactions (similar to our previous sci-ATAC-seq workflows [5]) followed by pooling all nuclei and processing samples through a largely unmodified standard 10X workflow (except that we overloaded the sample with 75,000 nuclei in a lane instead of the recommended 15,300 maximum capacity). The single-cell resolution and the degree of barcode collisions (i.e. instances where one barcode represents the contents of two or more cells) were evaluated using a “barnyard” experiment in which we mixed human and mouse nuclei - using either cell lines (human GM12878 nuclei mixed with mouse CH12.LX nuclei) or tissues (human lung nuclei mixed with mouse lung nuclei). Two mixing strategies were designed on the same 96-well plate: the nuclei from the two species were either pooled during tagmentation (“true barnyard”, which was used to reflect the rate of detected collisions caused by both pre- and post-pooling events) or after the tagmentation reaction (“pseudo-barnyard”, which was used to reflect the rate of detected collisions caused by post-pooling events only) (Fig. S1a). A mixed species experiment such as this (Fig. 1b) allows for an accurate estimation of collision rate since each index is expected to align uniquely to either the human or mouse reference genome. Indexes with cross-alignment indicate collisions and allow us to empirically scale cells loaded during droplet formation. The tagmentation reactions were performed with either a modified version of the “Omni” ATAC-seq protocol [10] or the 10X protocol (See Methods). After performing indexed tagmentation on a 96-well plate and pooling all nuclei, 75,000 nuclei from the pool were loaded onto a single 10X lane. The sample and cell-specific information of the resulting libraries was deconvoluted using the combination of three barcodes introduced during the workflow: a PCR barcode (i7) used to distinguish different lanes of the 10X, a GEM barcode introduced in the droplet, and a Tn5 barcode introduced during tagmentation (Fig. 1a). Unexpectedly, regardless of barnyard type (true vs pseudo), the initial experiment exhibited an extremely high collision rate (including estimated homotypic doublets [11]), i.e. 46.0% in a true-barnyard experiment mixing two cell lines, 44.4% in a pseudo-barnyard of cell lines, and 40.1% in a true barnyard mixing lung tissues (Fig. S1b). We also tested a second tagmentation buffer (provided in the 10X kit), but obtained similar results (47.4% estimated collision rate with a true barnyard of cell lines). However, limiting our measurement to GEMs with a single-Tn5 barcode demonstrated a remarkably reduced collision rate across all tested samples and buffers (4.7%, 3.3%, 4.4%, and 8.6% for the true barnyard of cell lines, pseudo-barnyard of cell lines, true barnyard with 10X buffer, and true barnyard with lung tissue, respectively). These results suggested that most multiplets were not arising from pre-pooling events, but instead were a consequence of cross-contamination due to Tn5 barcode swapping within droplets (Fig. S1c).

We tested three different strategies to eliminate this apparent in-droplet barcode-swapping (Fig. S1d; see Methods for details): (1) adding a second round of tagmentation with an additional (unamplifiable) duplex DNA prior to pooling to exhaust excess Tn5 (“Decoy DNA”); (2) supplementing the GEM reaction with a blocking oligo containing a reverse complement sequence of the Tn5 adaptor and an inverted dideoxythymidine (“dT”) at the 3’ end to inhibit the use of free Tn5 adaptors as amplification primers (“Blocking oligo”); or (3) supplementing the GEM reaction with another primer to enable exponential amplification instead of linear amplification in the droplet PCR (“SBS primer”) with the goal of outcompeting barcode-swapping. To facilitate better optimization of experiments in overloaded droplets without imposing a significant burden of sequencing for each condition tested, we also developed a method to sample a subset of droplets after in-droplet amplification. To do so, we took 10% of the volume of droplets immediately after amplification (but before breaking the droplets) and processed both the 10% sample and 90% in parallel (Fig. 1b). In this way, we could first sequence 10% of the loaded cells to evaluate data quality and subsequently sequence the remaining 90% if warranted. We then tested all three strategies head-to-head (Fig. S2a) and used a conservative cutoff of 1000 reads to identify cells for all conditions (Fig. 1c, Fig. S2b). While all three tested strategies mitigated some of the barcode swapping, we found that the SBS primer was most efficient - reducing the estimated collision rate of cell lines from 46.0% to 6.6% in the true barnyard and resulting in no collision cells observed in the pseudo-barnyard wells (Fig. 1d, Fig. S2c). Similar results were also seen in the lung barnyard (Fig. S2c), with a collision rate of 11.1% for the true barnyard and only a single collision observed in the pseudo-barnyard when spiking in the SBS primer (data not shown). We also used the fraction of reads mapping to the ENCODE-defined DNase I hypersensitive sites (FRiDHS) and the estimated library complexity (see Methods for calculations) to evaluate the performance across all three blocking conditions. Considering the data generated for cell lines, we found that the SBS primer provided the highest FRiDHS scores (a median of 61.5% for mouse cells and 60.3% for human cells, Fig. 1e and Fig. S2d) and a comparable complexity (a median of 25,504.1 for mouse and 27,298.6 for human, Fig. 1e) with Decoy DNA but a higher complexity than Blocking oligo (Fig. S2e). Coherent trends were also observed in the lung tissues (Fig. S2d,e). Interestingly, the SBS primer strategy also caused a shift in the fragment size distribution relative to the other conditions, indicating the exponential amplification of GEM reactions is biased toward small fragments given the same number of amplification cycles (Fig. S2f). A reduced number of cycles in droplet PCR, however, can partially recover the large fragment sizes (data not shown). Nonetheless, by optimizing this hybrid protocol of barcoded transposition followed by GEM amplification, we successfully developed a novel protocol that enables multiplexing of multiple samples and unbiased profiling of chromatin accessibility at extremely high throughput on the 10X Genomics platform. Having established a working protocol, we next sought to apply it to complex tissues to evaluate the assay’s performance. Below, we described the results from five primary samples.

### Profiling chromatin accessibility of human and mouse brain tissue

To evaluate the performance of txci-ATAC-seq in complex tissues, we initially generated chromatin accessibility profiles for human cortex and mouse whole brain samples using a true-barnyard scheme with two separate experiments to test nuclei inputs of 25,000 (∼1.5X the maximum recommended input) and 75,000 (∼4.5X the maximum recommended input) on the microfluidic device (see Methods). Libraries were sequenced to an average depth of 45,622 unique reads per cell, with an estimated saturation rate of 60.3% unique reads (Fig. S3a). We observed an estimated collision rate of 0.6% and 1.3% in the 25,000 and 75,000 inputs, respectively (Fig. S3b), which resulted in a ∼24-fold increase in the throughput of a standard 10X workflow at a comparable collision rate. A majority of droplet barcodes were assigned to a single 10X nucleus barcode, with 78.94% and 60.38% of droplets containing a single nucleus for 25,000 and 75,000 nuclei loadings, respectively (Fig. S3c). Overall, we captured 17,257 and 61,171 cells for the 25,000 and 75,000 nuclei loadings, respectively (Fig. S3d). To understand sample complexity, dimensionality reduction [12] and clustering [13] were performed on the human (Fig. 2a) and mouse (Fig. 3a) cells separately. We also identified and removed cryptic doublets within species to filter out the barcode collisions passing our initial species alignment filter (Methods) [14]. We generated gene activity scores (akin to a surrogate for gene expression) using *cis*-co-accessibility networks (CCANs) anchored on promoter regions [15]. A label-transfer algorithm then assigned cell types in comparison to published RNA datasets [16–18]. The high percentage of cells assigned to the same RNA-defined cell type per cluster supported the specificity of the label-transfer approach (Fig. S3e,f). We corroborated the assigned labels by examining the cluster-wise mean gene activity scores for canonical RNA markers of cell types (Fig. 2b and Fig. 3b) [19,20]. We next sought to define marker transcription factors (TFs) per cluster *de novo* by implementing an average “area under the curve” (AUC) value [21] across both gene activity and motif accessibility [22] scores in the human cortex (Fig. 2c). This approach allows for either gene activity or motif accessibility to be informative. For example, we found that the two human inhibitory neuron clusters could be distinguished by gene activity of LIM Homeobox 6 (LHX6), while motif usage differences between them were not significant and the motif is most accessible in astrocytes. In this case, the lack of distinction in motif usage is likely driven by other TFs of the LIM family that share a very similar motif, such as LIM Homeobox 2 (LHX2).

**Figure 2.**
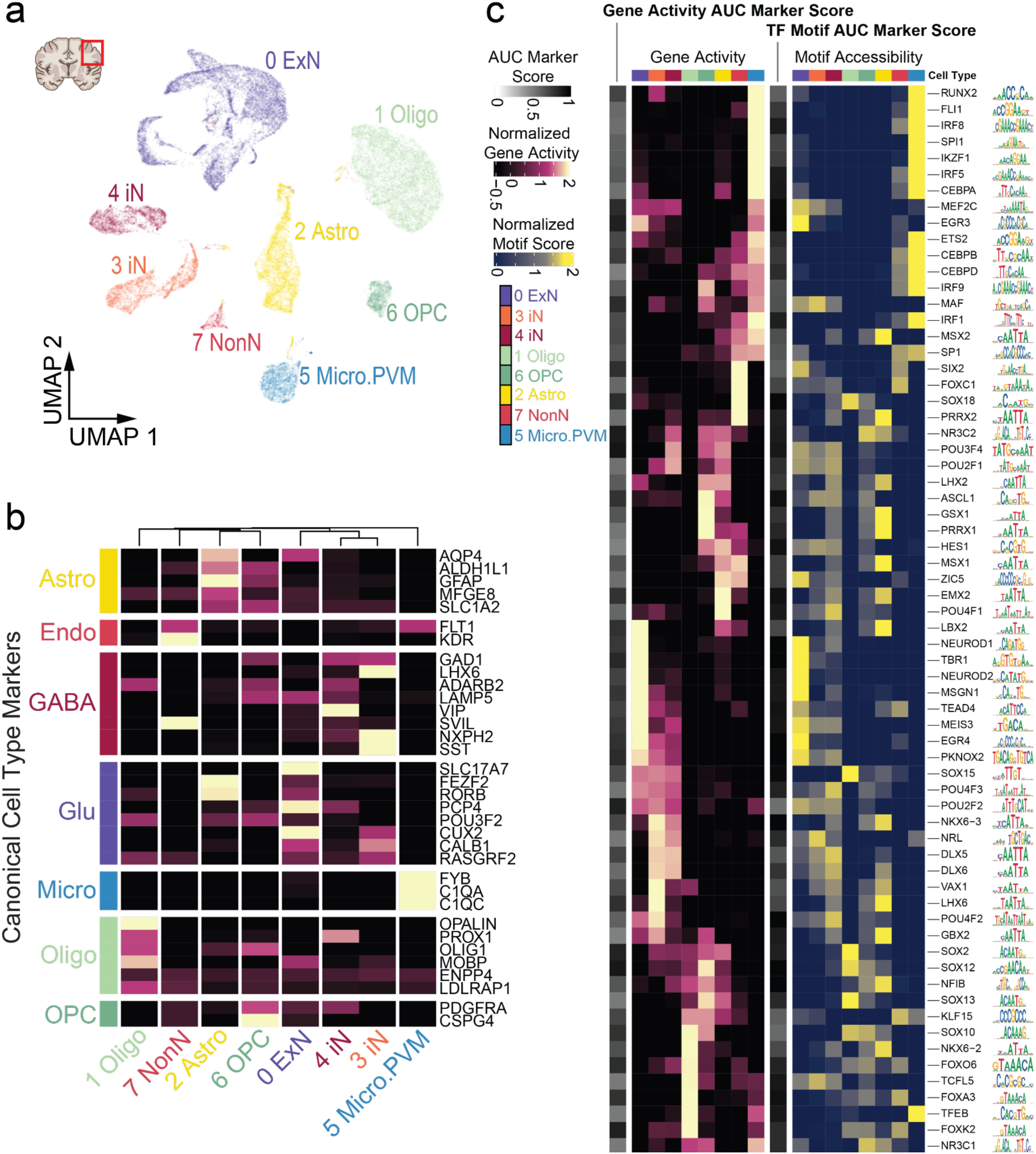
Cell type identification and marker assessment in human cortex sample. a) UMAP projection of human cortex nuclei (n = 28,663). Nuclei are colored by their predicted cell type. b) Heatmap of z-scored average gene activity score per cluster for canonical markers from Brain Map datasets. Astro: astrocytes; Endo: endothelial cells; ExN: excitatory neurons; GABA: GABAergic; Glu: Glutamatergic; iN: inhibitory neurons; Micro: microglia; Micro.PVM: microglia and perivascular macrophages; NonN: Non-neuronal; Oligo: oligodendrocytes; OPC: oligodendrocyte progenitor cells. c) *De novo* determination of TF marker genes through chromatin accessibility-derived gene activity (left) and TF motif usage (right). Z-scored average gene activity score and TF motif usage per cluster are plotted for the top 10 markers within each cluster. TF markers are ranked by AUC reported from one vs. rest Wilcoxon rank sum test. TF motifs are shown on the right as SeqLogos alongside heatmap rows.

**Figure 3.**
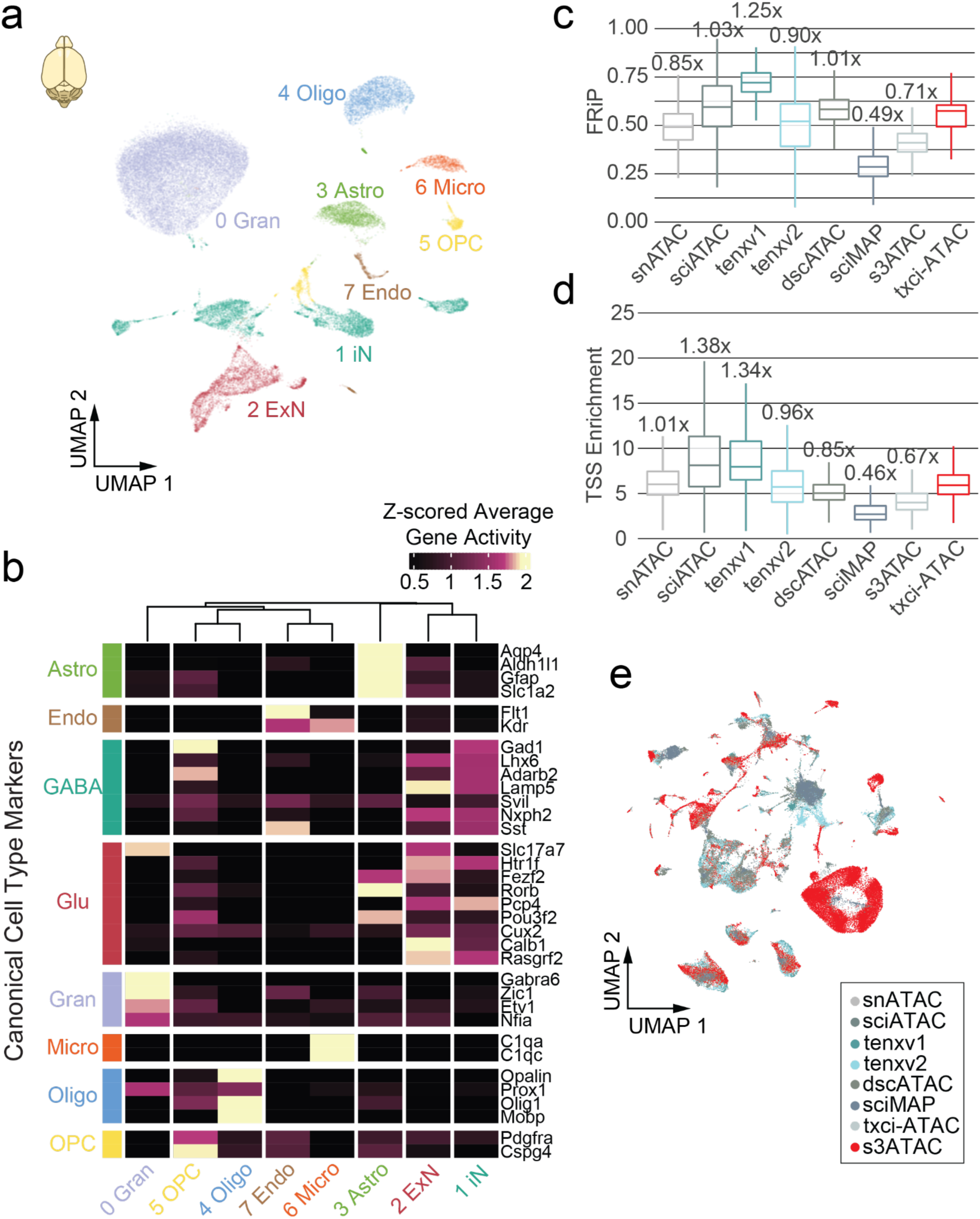
Cell type identification and marker assessment in mouse whole brain sample. a) UMAP projection of mouse brain nuclei (n = 49,765). Nuclei are colored by their predicted cell type. b) Z-scored average gene activity score per cluster plotted as a heatmap. Clusters are arranged by hierarchical clustering. Marker sets are from Brain Map marker genes. c) Boxplots of FRiP per technology using a unified peak set. Numbers over the boxplot reflect the fold-change of medians in comparison to txci-ATAC-seq. d) Boxplots of transcription start site (TSS) enrichment across technologies. Numbers over the boxplot reflect the fold-change of medians in comparison to txci-ATAC-seq. e) Harmony integrated UMAP projection of technologies (n = 75,845).

Since the mouse brain is a commonly profiled benchmark tissue of scATAC-seq methods, we compared our data to publicly available datasets for combinatorial indexing (snATAC-seq [23], sci-ATAC-seq [24], sci-MAP-seq [24], and s3-ATAC-seq [25]) and droplet-based (dscATAC-seq [9], 10X scATAC-seq v1 [26] and v2 [27]) chemistries. With all datasets merged we uncovered a unified peak set of 344,258 features of open chromatin in the mouse brain. txci-ATAC-seq performed comparably to the other technologies in terms of the fraction of reads in peaks (FRiP) (Fig. 3c) and transcription start site (TSS) enrichment at the level of individual cells (Fig. 3d), demonstrating the fourth-best FRiP (out of 8) and the fourth-best TSS enrichment. Notably, we observed that reducing the number of cycles used for in-GEM amplification (to 6 cycles) recovered the full spectrum of insert sizes in brain samples compared to the other techniques (Fig. S3g). We suspect that the different cycle numbers may explain the fragment size distribution previously observed with cell lines and lung samples as well (Fig. S2f). Estimating unique reads given a constant sequencing depth per cell (Fig. S3h), we noted that txci-ATAC-seq fell between the high-content ATAC-seq preparations (such as 10X scATAC-seq v2 chemistry or s3-ATAC-seq) and combinatorial methods (like snATAC-seq and sci-ATAC-seq). Also, txci-ATAC-seq integrated readily with other technologies on the unified peak set, with the exception of a notable increase in granule cells (Fig. 3e), potentially reflecting a higher concentration of cerebellum tissue during initial brain dissociation. Overall, txci-ATAC-seq enabled detailed epigenomic characterization of cell types in brain tissues, including the *de novo* definition of marker TFs by leveraging a combination of gene activity and TF motif usage. In tissue-matched comparisons across technologies, we found that txci-ATAC-seq performed equivalently in quality control metrics of library complexity and ATAC signals.

### Profiling chromatin accessibility of liver and lung tissue

To test the robustness of this strategy in different biological contexts, we multiplexed mouse liver and lung samples on a single 96-well plate with two replicates for each tissue (Fig. 4a). The last two rows of the plate were set up as a true-barnyard design by mixing mouse nuclei with human lung nuclei to estimate the internal collision rate for each sample. Two loading inputs (100,000 and 200,000 nuclei per lane) were tested and sequenced separately. Using a conservative cutoff of 1,000 reads to define a bona fide cell barcode (Fig. S4a,b), we recovered 67,251 (67.3%) and 104,987 (52.5%) nuclei from the 100,000 and 200,000 inputs, respectively (Fig. 4b). Since these libraries were sequenced to an average depth of 6,418.9 and 4,014.8 unique reads per cell for the 100,000 and 200,000 input libraries respectively (21.4% and 14.7% saturated, Fig. S4c,d), the slightly lower recovery rate observed for the 200,000 nuclei input may be due to the lower per-cell sequencing depth resulting in some likely cells failing to pass the read depth threshold (Fig. S4b,d). Collision rate estimates showed that pushing the loading throughput from 100,000 to 200,000 nuclei only raised the average rate from 3.6% to 4.4% (Fig. 4c). Overall, these libraries increased the yield of usable nuclei by nearly 22-fold in comparison to the standard 10X Chromium scATAC-seq at the same collision rate. While the collision rate appeared to be tissue-dependent within this experiment (with an average of 3.4% for liver and 4.6% for lung), the fold increase in the number of cells that could be processed at a 10X-equivalent collision rate aligned well with what we observed in brain tissues. In addition, we again compared a series of quality metrics between our txci-ATAC-seq data and previously obtained sci-ATAC-seq data on the same tissues [3] and demonstrated that the data generated with txci-ATAC-seq had a substantially higher quality than the original combinatorial indexing assay (Fig. 4d-f): the median FRiDHS increased from 25.5% to 56.5% for liver and from 22.8% to 53.0% for lung; the median TSS enrichment score increased from 2.5 to 4.5 for liver and from 3.2 to 5.1 for lung; the median complexity increased from 16,472.2 to 25,338.4 for lung while it decreased from 33,123.4 to 21,362.2 for liver. After filtering out low-quality nuclei and putative doublets (see Methods), we generated chromatin accessibility profiles for 152,508 primary cells, including 73,280 mouse lung nuclei, 63,429 mouse liver nuclei, and 15,799 human lung nuclei (59,348 of the nuclei recovered from the 100,000 input library and 93,160 of the nuclei recovered from the 200,000 input library).

**Figure 4.**
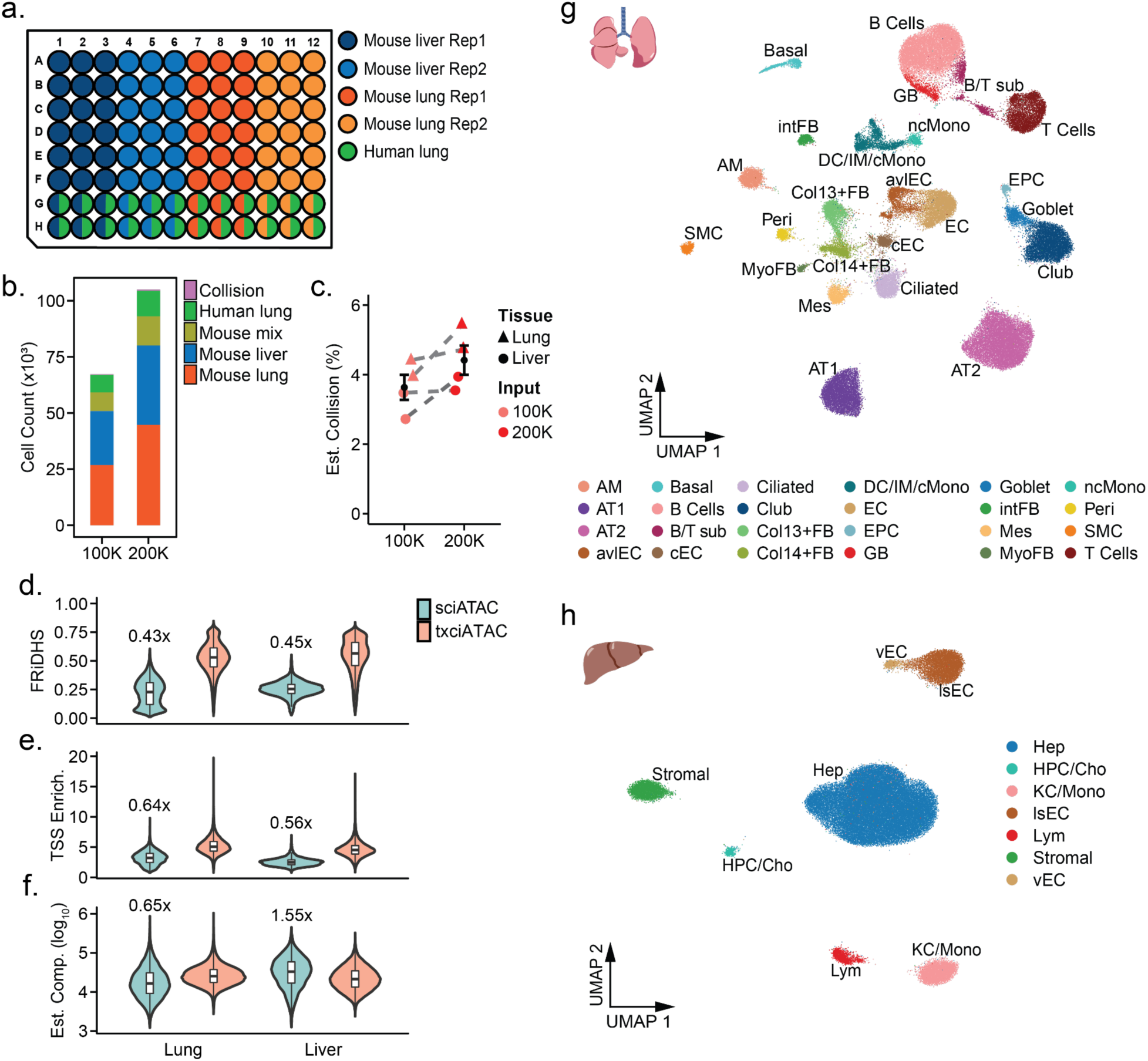
txci-ATAC-seq generates high-quality single-cell ATAC-seq data on multiple tissues in parallel at scale. a) Well assignment showing the multiplexing of primary samples. Rows 7 and 8 provide an estimate of the empirical collision rate for each sample by mixing human lung nuclei with mouse nuclei isolated from each corresponding tissue. b) The number of nuclei (with a cutoff of 1000 reads) recovered at each loading input. The colors denote the samples multiplexed for each 10X reaction. c) The estimated collision rate of each mouse sample when loading either 100,000 or 200,000 nuclei. The filled circle and triangle indicate the mouse liver and lung tissues, respectively. The error bar shows the standard error and the black point represents the sample mean at each input. The same samples between the two loading inputs are connected by a gray dashed line. d-f) The comparison of quality metrics between sciATAC-seq and txci-ATAC-seq for each cell in mouse lung and liver tissue. The (d) FRiDHS, (e) TSS enrichment score, and (f) estimated complexity (on a log_10_ scale) indicate the performance of single-cell ATAC-seq methods. The numbers over the violin plots reflect the fold-change in median compared to txci-ATAC-seq. g) UMAP visualization of mouse lung nuclei (n = 73,280) integrating two replicates across two loading inputs. Nuclei are colored by their predicted cell type. h) UMAP visualization of mouse liver nuclei (n = 63,429) integrating two replicates across two loading inputs. Abbreviations: AM, alveolar macrophages; AT1, alveolar type 1 epithelial cells; AT2, alveolar type 2 epithelial cells; avlEC, arterial/venous/lymphatic endothelial cells; B/T sub, B and T cell subpopulation; cEC, capillary endothelial cells; Col13+FB, collagen type XIII α 1 chain positive fibroblasts; Col14+FB, collagen type XIV α 1 chain positive fibroblasts; DC/IM/cMono, dendritic cells/interstitial macrophages/classical monocytes; EC, endothelial cells; EPC, epithelial progenitor cell; GB, germinal B cells; Hep, hepatocytes; HPC/Cho, hepatic progenitor cells/cholangiocytes; KC/Mono, Kupffer cells/monocytes; intFB, interstitial fibroblasts; lsEC, liver sinusoidal endothelial cells; Lym, lymphocytes; Mes, mesothelial cells; MyoFB, myofibroblasts; ncMono, nonclassical monocytes; Peri, pericytes; SMC, smooth muscle cells, vEC, venous endothelial cells.

To dissect the diverse chromatin landscapes present in these heterogeneous tissues, we performed an iterative peak calling and clustering method to parse out the distinct cell populations. In brief, we called peaks on aggregated reads for all cells, scored individual cells for insertion events in these reference peaks, and then carried out dimensionality reduction and cluster identification using Seurat [28]. A second round of peak calling was performed on cells from each cluster separately, and the peaks identified for all clusters were then merged and used as a reference set to perform dimensionality reduction again and re-cluster the cells. The associated cell type for each cluster was predicted by label transfer using previously published single-cell/single-nucleus RNA-seq (scRNA-seq/snRNA-seq) and sci-ATAC-seq datasets from mouse lung tissue (Fig. S5; [3,29]), mouse liver tissue (Fig. S6; [3,30]) and human lung tissue (Fig. S7; [31–33]). The predicted labels were further manually curated according to the top gene activity scores (by summing the read counts in gene bodies and promoters [16]) measured in each cluster. As a result, we identified 24 clusters representing distinct cell types in mouse lung tissue (Fig. 4g) and 7 clusters in mouse liver tissue (Fig. 4h). Even relatively rare cell types such as goblet cells (1335 cells,1.8% of total), pericytes (833 cells, 1.1% of total), and myofibroblasts (366 cells, 0.5% of total) in mouse lung tissue were identified, in contrast to the previous sci-ATAC-seq atlas. To evaluate the performance of txci-ATAC-seq in cell type prediction, we randomly subsampled (without replacement) our mouse lung data to have the same number of cells as that in sci-ATAC-seq 1,000 times and ran cell type prediction using label transfer. As compared to the combinatorial indexing assay, txci-ATAC-seq exhibited improved prediction accuracy (Fig. S8). Further validating our approach in human lung tissue, we identified 9 distinct clusters (Fig. S9a) and found that the human lung nuclei exhibited consistent clustering (Fig. S9b) and data quality (Fig. S9c-e), regardless of which mouse sample they were mixed with in the barnyard experiment. In addition, while we did observe some stratification of mouse hepatocytes according to the individual mouse replicate, no other cell type showed evidence of batch effects (Fig. S10).

### Development of Fast-txci-ATAC-seq to improve multiplexing capability

While txci-ATAC-seq enables multiplexing of multiple samples, processing dozens or hundreds of samples in a single experiment is still laborious. To further increase the multiplexing capability of our method, we developed a “faster” protocol for txci-ATAC-seq (Fast-txci-ATAC-seq) by performing the transposition reaction directly on frozen nuclei, which enables freezing nuclei on a 96-well plate or in 8-tube strips sequentially over time and then performing barcoded transposition immediately upon thawing the nuclei. To evaluate the performance of the faster version of our protocol, we applied it to mouse lung nuclei that were isolated from either wild-type (WT) or club cell secretory protein deficient (CC16^-/-^) mice with three replicate lungs for each genotype. The standard txci-ATAC-seq was also performed on the same samples separately. CC16 is a secreted protein encoded by the *Scgb1a1* gene that is produced predominantly by club cells, an epithelial cell type of the airways. This “pneumoprotein” plays an important role locally in protecting the lung against oxidant injury [34] and inflammatory diseases, such as asthma [35] and chronic obstructive pulmonary disease (COPD) [36]. It has also been linked with more systemic effects on human health as evidenced by its association with overall cancer risk [37]. After processing and pooling all samples for each protocol (Fig. S11a), we loaded 50,000 and 100,000 nuclei on the 10X Genomics platform for Fast-txci-ATAC-seq and used 100,000 and 200,000 nuclei as inputs for the standard assay. The removal of low-quality nuclei and predicted doublets resulted in similar recovery rates between the two protocols with 44.3% nuclei (10,937 WT nuclei and 11,213 CC16^-/-^ nuclei) at the 50,000 input and 43.6% nuclei (21,688 WT nuclei and 21,961 CC16^-/-^ nuclei) at the 100,000 input for the faster protocol, compared to 50.0% nuclei (24,962 WT nuclei and 25,011 CC16^-/-^ nuclei) at the 100,000 input and 52.1% nuclei (51,536 WT nuclei and 52,627 CC16^-/-^ nuclei) at the 200,000 input for the standard txci-ATAC-seq (Fig. S11b). An examination of QC metrics demonstrated that both assays can provide high-quality single-cell data despite a slightly lower complexity observed in the faster version (Fig. S11c-e). Using the iterative clustering strategy and label transfer with a scRNA-seq reference, we identified 23 distinct cell clusters in mouse lungs profiled by the standard txci-ATAC-seq (Fig. 5a) and then used them to further annotate the Fast-txci-ATAC-seq lungs. The joint embedding of both assays revealed that the faster protocol recapitulated the mouse lung heterogeneity in chromatin accessibility characterized by the standard protocol (Fig. S11f) with minimal batch effects (Fig. S11g).

**Figure 5.**
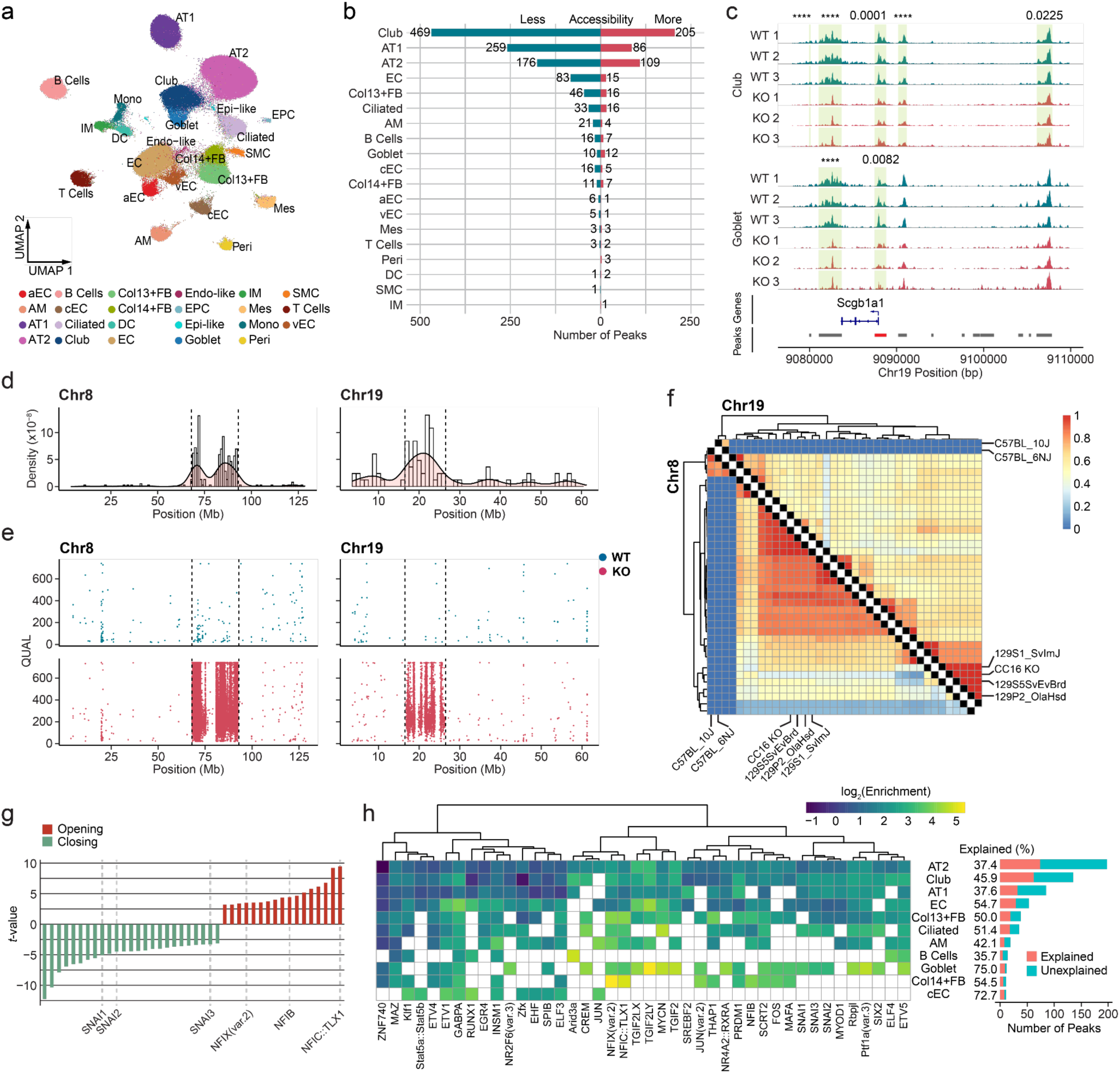
Chromatin accessibility dynamics induced by CC16 deficiency and genetic variants. a) UMAP visualization of WT and CC16^-/-^ mouse lung nuclei (n = 154,136) across two loading inputs by integrating 6 animals with 3 replicates from each group. Nuclei are colored by their predicted cell type. The abbreviation of cell labels was described in Fig. 4 except for aEC (arterial endothelial cells), Endo-like (endothelial-like cells), and Epi-like (epithelial-like cells). b) The number of differential peaks identified between CC16^-/-^ and WT samples for each cell type. The blue bars indicate the peaks less accessible in the knockout samples and the red bars represent the more accessible peaks. c) Aggregated chromatin accessibility surrounding the *Scgb1a1* (CC16 gene) locus in club and goblet cells per sample. The aggregated accessibility signal for each sample was normalized by the scaling factor that was computed as the number of cells in the sample multiplied by the mean sequencing depth for the cells in that sample. The WT tracks are labeled in blue and the knockout ones are in red. The genomic regions for the significantly less accessible peaks identified in CC16^-/-^ samples per cell type are highlighted by green shade (5 peaks in club cells and 2 peaks in goblet cells). The associated adjusted p-value is shown above the tracks at their corresponding peak region. Adjusted p-values less than 0.0001 are given four asterisks. The peak annotation for the promoter region of *Scgb1a1* is colored red. d) Chromosomal distribution of the midpoint of differential peaks identified on chromosomes 8 and 19 with the genomic location and density estimate plotted on the x- and y-axis, respectively. e) Chromosomal distribution of SNVs identified on chromosomes 8 and 19 for both WT (blue) and CC16^-/-^ (red) samples. The regions between the dashed lines indicate the SNV hotspots where the knockout samples exhibited a substantially higher number of SNVs than WT samples. The y-axis shows the Phred-scaled quality score generated by BCFtools. f) Heatmap showing the Jaccard similarity between the hotspot SNVs identified in CC16^-/-^ lungs and the SNPs derived from 36 different strains on chromosome 8 (lower triangle) and 19 (upper triangle). g) “Functional” motifs for which gains or losses of the motif instances are associated with significant changes in chromatin accessibility. The motifs associated with increased chromatin accessibility (“opening”) are shown in red and those associated with decreased chromatin accessibility (“closing”) are colored in green. The y-axis represents the Student’s *t*-test statistic value. Two motif families (one for transcriptional activators and one for transcriptional repressors) are highlighted on the x-axis. h) Cell-type-specific enrichment for the motifs that explain chromatin accessibility changes in SNV hotspots. The bar plot next to the enrichment heatmap shows the total number of differential peaks located in the SNV hotspots for each cell type, which is stratified by the peaks that can be explained by the SNV-driven difference in motif presence (red) and unexplained peaks (blue). The values next to the bars denote the percentage of peaks explained. Only the cell types with more than 10 differential peaks identified in the SNV hotspots are shown.

### Cell-type-specific regulation of chromatin accessibility in CC16^-/-^ mouse lungs

We next used the cells profiled by the standard assay to explore meaningful differences in chromatin accessibility between WT and CC16^-/-^ mice. We were initially interested to identify differences of biological import, including 1) chromatin accessibility of the *Scgb1a1* locus being restricted to club and goblet cells (Fig. S12a), 2) significantly differentially accessible peaks across a variety of cell types (Fig. 5b; Table S1) - many of which were only identifiable at such high throughput (Fig. S12b,c) - 3) some evidence for potential autoregulation of CC16 (Fig. 5c and Fig. S12d), and 4) differential peaks enriched for TF motifs (Fig. S13a,b; Table S2) and molecular pathways (Fig. S13c; Table S3). However, we also noted an unexpected number of differential peaks in two genomic loci on chromosomes 8 and 19 (Fig. 5d and Fig. S13d). While the *Scgb1a1* gene is located on chromosome 19 (mm10 chr19:9,083,636-9,087,958), this chromosome-specific enrichment of differential peaks was unexpected. To better understand the concentration of signal in these two loci, we carried out variant calling on the ATAC-seq data and identified 5,909 single nucleotide variants (SNVs) differing from the reference genome in WT samples and 50,054 SNVs in CC16^-/-^ samples (Fig. S13e). The vast majority of SNVs in the CC16^-/-^ samples were located in two hotspots (Fig. 5e) on chromosome 8 (n = 37,822; 75.6%) and chromosome 19 (n = 5,610; 11.2%), essentially perfectly matching the locations where the differential peaks were identified. To trace the origin of the CC16^-/-^ SNVs, we further mapped the hotspot SNVs to the single nucleotide polymorphism (SNP) profiles that were previously defined in 36 different mouse strains relative to the C57BL/6J mouse reference genome [38]. Almost all of our identified SNVs matched to the SNPs identified in the three 129 strain references (Fig. 5f) on chromosome 8 (an average of 96.3% of SNVs matched the SNPs defined in each of the 129 strains) and chromosome 19 (an average of 96.6% SNVs matched the SNPs defined in each of the 129 strains). Given that the CC16^-/-^ mice were generated using 129-derived embryonic stem (ES) cells, we conclude that the hotspot SNVs are remnants of the 129 genome, a common problem with knockout models [39]. Notably, we found ∼90% of SNVs residing in intronic and intergenic regions for both WT and CC16^-/-^ samples (Fig. S13f), suggesting ATAC-seq may have been a particularly powerful choice of assay for capturing such genetic variation and thus may serve as a cost-effective alternative to whole genome sequencing in genotyping knockout models.

Although the SNV-driven phenotype confounded the analysis of *Scgb1a1* effects, it provided an opportunity to explore the extent and mechanism by which genetic variants can modulate chromatin accessibility, even in a cell-type-specific manner. To this end, we took all the peaks that were differentially accessible in the hotspot regions (413 peaks) and looked for TF motifs that were gained or lost due to SNVs. The functional motifs were defined as those whose chromatin accessibility exhibited a significant positive or negative correlation with the gain or loss of the motif in the knockout mice relative to the WT mice. We identified 42 functional motifs and found that gaining the motifs for the transcriptional activators, e.g. certain members of the nuclear factor I (NFI) family and the ETS-domain family, tended to increase chromatin accessibility (Fig. 5g and Fig. S14a). On the other hand, gaining a repressor motif, such as motifs for the Snail and Scratch families, was likely to reduce chromatin accessibility (Fig. 5g and Fig. S14b). Finally, we investigated whether there were cell-type-specific enrichments for specific functional motifs being gained or lost (Fig. 5h). We observed that gains and losses of NFI TFs, including NFIB, NFIC::TLX1, and NFIX (var.2), were highly enriched in both Col13^+^ and Col14^+^ fibroblasts and similarly, gains and losses of the ARID3A motif, which is required for B cell lineage development [40], was highly enriched in differential peaks in B cells. In sum, functional motifs being gained or lost were able to account for a substantial number of differentially accessible peaks observed in the SNV hotspot regions in different cell types - ranging from 35.7% of differentially accessible peaks from the regions for B cells to 75% of differentially accessible peaks from the regions for goblet cells (Fig. 5h).

## Discussion

Limited throughput, prohibitive cost, and variance between batches have put some limitations on the implementation of single-cell techniques, which nonetheless are proving invaluable resources for studying health and disease. To reduce those limitations, we paired combinatorial indexing with a droplet-based microfluidic system to substantially increase the scalability of the commercial single-cell device by loading up to 200,000 nuclei in a single emulsion reaction. In addition, a “faster” version protocol was developed, which greatly expedited sample processing and improved the multiplexing capability. The scalability and flexibility allow txci-ATAC-seq to establish unbiased regulatory definitions across various disease, genetic, and/or environmental states. Other strategies do exist for multiplexing samples on microfluidic single-cell platforms, such as membrane barcoding-based approaches that tag cellular or nuclear membrane components [41–43] and genetic deconvolution of samples [44]. However, those methods index at the cellular/nuclear level and so the scalability is restricted by the maximum number of singlets that can be generated because multiplets cannot be deconvoluted. Conversely, the molecular indexing strategy used in our design along with that previously implemented on a different commercial instrument (dsciATAC-seq) [9] and on RNA profiling (scifi-RNA-seq) [45] allows for multiplets to be deconvoluted, resulting in the ability to load substantially more nuclei per lane and therefore provide larger-scale sample multiplexing and increased cost savings. Because a range of nuclei inputs was tested in this study, we also computed the number of droplets containing a single Tn5 barcode to estimate the number of singlets recovered at each input. We found that the number of singlets peaked around the 100,000 nuclei input level with an average of 26,402 single-nucleus droplets generated (Fig. S15). This represents the maximum empirical yield for single-cell multiplexing techniques based on cellular indexing, such as “cell hashing” [41,42], MULTI-seq [43], and SnuBar [8]. In contrast, our molecular indexing approach yielded 17,257 deconvoluted nuclei at the 25,000 input, 34,568 deconvoluted nuclei at the 50,000 input, 61,171 deconvoluted nuclei at the 75,000 input, an average of 68,216 deconvoluted nuclei at the 100,000 input and an average of 128,334 deconvoluted nuclei at the 200,000 input. Given our results and the reported metrics for scifi-RNA-seq [45], we are confident that an even higher throughput is achievable with txci-ATAC-seq (even more so if one were to leverage 384 barcoded transposition reactions). Furthermore, the recently reported multi-omic approach, ISSAAC-seq [46], could also potentially benefit from this framework, by integrating the same tagmentation strategy applied here with an initial step of *in situ* reverse transcription, to generate a novel high throughput, cost-efficient multi-omics assay. We also note that the design of txci-ATAC-seq is directly applicable to other existing single-cell methods employing a combinatorial indexing framework, such as sci-MET [47], CRISPR-sciATAC [48], and sci-CAR [49].

The improved study design and statistical rigor made possible by more cost-effective inclusion of replicates and larger sample sizes with techniques such as txci-ATAC-seq will be essential for realizing the full potential of single-cell approaches. In addition, the scalability of scATAC-seq techniques also plays an important role in identification of peaks for rare cell populations. In the absence of a comprehensive catalog of regulatory elements, peak calling is an essential step to define features in both bulk and single-cell ATAC-seq data analysis. The power to call peaks, however, heavily depends on the number of reads used [50]. For single-cell data, that means profiling the accessible regions from a rare cell population is not only limited by the sequencing depth per cell but also by the number of cells captured from that population. Therefore, an ultra-high throughput scATAC-seq method, like txci-ATAC-seq, will enable finer definitions of peaks and should better characterize particularly dynamic or heterogeneous systems.

The CC16^-/-^ mice characterized here have been used by several groups to investigate the role of CC16 in COPD and infectious diseases [36,51,52]. We found that the remnant 129 genetic material elicited profound changes in chromatin accessibility (in a cell-type-specific manner in many instances), requiring caution when evaluating the existing congenic knockout models. In addition, we identified 42 different motifs gained or lost in at least one differentially accessible peak from the 129 strain regions, which were capable of explaining the observed accessibility changes for 37.5% of those peaks. The remainders may have been caused by more subtle changes in motif affinity, trans effects, or may not have been tested in our analysis (as we required both gained and lost events for a given motif to be considered).

There are several caveats worth keeping in mind when interpreting our results. First, to marry microfluidics and combinatorial indexing on the 10X system, we converted the in-droplet linear amplification into an exponential amplification. This could result in major differences in amplification behavior. However, we have not systematically tested the optimal number of cycles in this regime. In addition, our analysis approach is based on the assumption that each droplet contains at most one barcode bead. It is worth noting, however, that “barcode multiplets” (i.e. the droplets containing multiple beads or the beads containing multiple oligonucleotide barcodes) have been observed in 10X Chromium scATAC data [53]. The resulting artifact “cells” may confound the interpretation of txci-ATAC-seq, and so implementation of methods to detect and remove the effects of “barcode multiplets” may be warranted.

## Conclusions

Taken together, txci-ATAC-seq provides unprecedented opportunities to generate unbiased single-cell atlases of chromatin accessibility for large cohorts with various genetic backgrounds or case-control studies, thus establishing reliable references of single-cell chromatin landscapes in a variety of experimental settings. We hope that this method will encourage more widespread adoption of scATAC-seq, a powerful technique for understanding organismal development and disease processes.

## Methods

### Cell lines

The GM12878 (Coriell Cell Repository) and CH12.LX (kind gift from the Sherman Weissman lab) cells were cultured at 37 °C with 5% CO2 in RPMI 1640 medium (GIBCO, cat. no. 11875-093) containing 15% FBS (GIBCO, cat. no. 10437-028), 100 U/ml Penicillin Streptomycin (GIBCO, cat. no. 15140-122). Cells were counted and split into either 300,000 (GM12878) or 100,000 (CH12.LX) cells/ml three times a week.

### Human and mouse brain tissue samples

Human cortex samples from the middle frontal gyrus were sourced from the Oregon Brain Bank from a 50-year-old female of normal health status. Samples were collected by an OHSU neuropathologist, placed into a labeled cassette, and cryopreserved in an airtight container in a - 80 °C freezer. The duration of time between the time of death and brain biopsy sample freezing, or post-mortem interim (PMI), was <24 hours.

Mouse brain tissue was collected as discarded tissue from mice used for unrelated studies approved by the OHSU IACUC. Whole mouse brains were dissected from sacrificed C57BL/6J mice and flash-frozen in an isopentane-LN2 double-bath and stored at -80 °C.

### Mouse lung and liver tissue samples

All animal activity was approved by the University of Arizona IACUC. Mice were euthanized via exsanguination followed by cervical dislocation to ensure death. For the samples used to evaluate the performance of txci-ATAC-seq in Fig. 4, whole mouse lungs and liver were dissected from 2 male C57BL/6J mice that were 24 weeks old.

For the samples used to study the CC16-mediated chromatin dynamics in Fig. 5, age-matched (∼8 weeks) WT and CC16^-/-^ male mice on a C57BL/6J background (as described in [54,55]) were used to dissect whole lungs. Three replicates from each genotype were profiled. All six animals were born and raised in the same room and were tested to be specific-pathogen free according to standard protocols using sentinel mice from the same room.

The dissected samples were flash-frozen in liquid nitrogen and then transferred to -80 °C for long-term storage.

### Human lung tissue samples

Lung pieces were obtained from two deceased male donors (a 36-year-old American Indian and a 62-year-old Hispanic Latino) as soon as possible after the time of death through the Arizona Donor Network. All human lung samples were quickly frozen in the -80 °C freezer and stored there prior to nuclear extraction.

### Nuclei isolation

#### Nuclei isolation of cell lines

The nuclei isolation followed the procedures described in [10]. The cells were collected and washed with 1x PBS (pH 7.4, Gibco, cat. no. 10-010-023) supplemented with 0.1% BSA (New England Biolabs, cat. no. B9000S), and then resuspended in 200 μl of ATAC-seq lysis buffer, which was made by supplementing ATAC resuspension buffer (RSB) with detergents (see below). RSB buffer is 10 mM Tris-HCl (pH 7.5, Invitrogen, cat. no. 15567027), 10 mM NaCl (Invitrogen, cat. no. AM9759), and 3 mM MgCl2 (Invitrogen, cat. no. AM9530G) in nuclease-free water. RSB was made in bulk and stored at 4 °C long-term. On the day of the experiment, the ATAC lysis buffer was made by adding 0.1% IGEPAL (Sigma, cat. no. I3021), 0.01% digitonin (Invitrogen, cat. no. BN2006), and 0.1% Tween-20 (Bio-Rad, cat. no. 1610781) to RSB. The detergent percentages reported are final concentrations. After resuspending cell pellets in the lysis buffer, they were incubated on ice for 3 min, and then the lysis was stopped by adding 1 ml RSB containing 0.1% Tween-20. The nuclei were counted with a hemocytometer by diluting 10 μl nuclei in 40 μl of 2x Omni TD Buffer (20 mM Tris HCl pH 7.5, 10 mM MgCl2, and 20% Dimethyl Formamide) followed by adding 50 μl Trypan blue solution. In our previous report [50], we found that adding nuclei straight to Trypan blue solution will cause inflation of nuclei, and diluting nuclei in TD buffer before exposure to Trypan blue improves the nuclei integrity. Following counting, we centrifuged nuclei at 500 r.c.f for 10 min at 4 °C and removed the supernatant. Then, the nuclei were either used to perform downstream experiments directly or resuspended in a nuclei-freezing buffer (NFB) containing 50 mM Tris-HCI (pH 8.0, Invitrogen, cat. no. 15568025), 5 mM Magnesium Acetate (Sigma, cat. no. 63052), 25% glycerol (VWR, cat. no. RC3290-32), 0.1 mM EDTA (Fisher, cat .no. AM9260G), 5 mM DTT (Fisher, cat. no. P2325), and 2% (v/v) protease inhibitor (Sigma, cat. no. P8340) for storage. The NFB was adopted from [56] and we previously used this buffer for preservation of nuclei for sci-ATAC-seq [2,3,57]. After diluting in NFB, 1 ml aliquots of the nuclei were flash-frozen in liquid nitrogen and then transferred to a liquid nitrogen dewar for long-term storage.

#### Nuclei isolation from brain tissue

At the time of nuclei dissociation, 50 ml of nuclei isolation buffer (NIB-HEPES) was freshly prepared with final concentrations of 10 mM HEPES-KOH (Fisher Scientific, BP310-500 and Sigma Aldrich 1050121000, respectively), pH 7.2, 10 mM NaCl (Fisher Scientific S271-3), 3mM MgCl2 (Fisher Scientific AC223210010), 0.1 % (v/v) IGEPAL CA-630 (Sigma Aldrich I3021), 0.1 % (v/v) Tween-20 (Sigma-Aldrich P-7949) and diluted in PCR-grade Ultrapure distilled water (Thermo Fisher Scientific 10977015). After dilution, two tablets of Pierce™ Protease Inhibitor Mini Tablets, EDTA-free (Thermo Fisher A32955) were dissolved and suspended to prevent protease degradation during nuclei isolation.

An at-bench dissection stage was set up prior to nuclei extraction. A petri dish was placed over dry ice, with fresh sterile razors pre-chilled by dry-ice embedding. 7 ml capacity Dounce homogenizers were filled with 2 ml of NIB-HEPES buffer and held on wet ice. Dounce homogenizer pestles were held in ice-cold 70% (v/v) ethanol (Decon Laboratories Inc 2701) in 15 ml tubes on ice to chill. Immediately prior to use, pestles were rinsed with chilled distilled water. For tissue dissociation, mouse and human brain samples were treated similarly. The still-frozen block of tissue was placed on the clean pre-chilled petri dish and roughly minced with the razors. Razors were then used to transport roughly 1 mg of the minced tissue into the chilled NIB-HEPES buffer within a Dounce homogenizer. Suspended samples were given 5 minutes to equilibrate to the change in salt concentration prior to douncing. Tissues were then homogenized with 5 strokes of a loose (A) pestle, another 5-minute incubation, and 5-10 strokes of a tight (B) pestle. Nuclei were transferred to a 15 ml conical tube and pelleted with a 400 r.c.f centrifugation at 4 °C in a centrifuge for 10 minutes. The supernatant was removed and pellets were resuspended in 5 ml of ATAC-PBS buffer (APB) consisting of 1X PBS (Thermo Fisher 10010) and 0.04mg/ml (f.c.) of bovine serum albumin (BSA, Sigma Aldric A2058). Samples were then filtered through a 35 µm cell strainer (Corning 352235). A 10 μl aliquot of suspended nuclei was diluted in 90 μl APB (1:10 dilution) and manually counted on a hemocytometer with Trypan Blue staining (Thermo Scientific T8154). The stock nuclei suspension was then diluted to a concentration of 2,857 nuclei/μl in APB. Dependent on experimental schema, pools of tagmented nuclei were combined to allow for the assessment of pure samples and to test index collision rates.

#### Nuclei isolation of human lung, mouse lung, and mouse liver tissue

The human and mouse samples were dissected and stored at -80 °C. The nuclei isolation procedure of lung and liver tissues was performed following the single-nucleus isolation protocol described in [58]. To do so, we cut a ∼0.1-0.2 g piece from either human or mouse samples removed from -80 °C and kept it on dry ice until use. The tissue block was thawed almost completely on ice for 1 min and then injected with 1 ml of cell lysis buffer, which was made of 1x cOmplete protease inhibitor cocktail (1 tablet per 10 ml solution, Sigma-Aldrich, Cat. 11836153001) in Nuclei EZ prep buffer (Sigma-Aldrich, Cat. NUC101), into the center of the tissue with a 30G needle and syringe. Following lysis buffer injection, the tissue was chopped into small pieces with scissors and then transferred along with the lysing buffer into a gentleMACS C tube (Miltenyi Biotec, Cat. 130-096-334). An additional 1 ml of lysing buffer was added into the C tube to make a final volume of 2 ml. The minced tissue was then homogenized using a gentleMACS tissue dissociator by running the ‘m_lung_01’ program followed by the first 20 sec of the ‘m_lung_02’ program. After homogenization, tissue lysate was briefly centrifuged to reduce foam and then passed through a 40 μm cell strainer in a 50ml tube. After passing the sample through, the strainer was rinsed with 4 ml of washing buffer (PBS with 1% BSA). The nuclei were counted with a hemocytometer (see “nuclei isolation of cell lines” for details), and centrifuged at 500 r.c.f for 5 min at 4 °C. Then, we removed the supernatant and resuspended the nuclei in the NFB to make a concentration of 4-5 million nuclei/ml. 1 ml aliquots of the nuclei were flash-frozen in liquid nitrogen and then transferred to a liquid nitrogen dewar for long-term storage.

### Sample multiplexing

A 96-well plate pre-loaded with 5 μl of 500 nM pre-indexed Tn5 transposase per well (iTSM plate, kind gift of Illumina Inc.) was used to multiplex samples and perform barcoded transposition. Before using, the iTSM plate was thawed on ice and briefly mixed at 1400 rpm for 30 seconds on a pre-chilled thermomixer, and then quickly spun to collect the enzyme at the bottom of the wells. To avoid sequencing with a custom recipe, the Tn5 enzyme was loaded with a common Tn5ME-A and a custom Tn5ME-B containing a partial sequence of i7 TruSeq primer (see Table S4 for oligo sequence) and an 8 bp unique barcode (Table S5). Both Tn5ME-A and Tn5ME-B were annealed to the Tn5MErev (Table S4) before loading to Tn5.

### Barnyard experiments

Two different barnyard settings were designed to estimate the total collisions arising from pre- and/or post-pooling events. To test the total collision rate, the human and mouse cells were mixed in the same well at a 1:1 ratio to perform barcoded transposition (“true barnyard”). The collision rate driven by events downstream of pooling was evaluated by performing barcoded transposition on wells containing pure species (“pseudo-barnyard”) and pooling the human and mouse nuclei afterward. Detailed information about the cell sources used in each barnyard assay and each figure is shown in Table S6.

### Optimization of *txci-ATAC-seq* protocol

#### Coupling barcoded transposition with standard 10X protocol

The nuclei isolated from human and mouse lungs were removed from the liquid nitrogen dewar (See “Nuclei isolation of primary samples” for details) and then thawed in the water bath at 37 °C for 1 to 2 min until a tiny ice crystal remained. After thawing, the nuclei stored in 1 ml freezing buffer were diluted with 3 ml RSB supplemented with 0.1% Tween-20 and 0.1% BSA (RSB washing buffer) and then centrifuged at 500 r.c.f for 10 mins in a pre-chilled (4 °C) swinging-bucket centrifuge. The nuclei pellet was resuspended with another 1 ml of RSB washing buffer and then transferred to a 1.5 ml LoBind tube through a 40 μm Flowmi Cell strainer (Bel-Art SP Scienceware, Cat. 14-100-150). The filtered nuclei were pelleted at 500 r.c.f for 5 min in a pre-chilled fixed-angle centrifuge and then resuspended in 25 μl of 1.25x Tagment DNA Buffer (Nextera XT Kit, Illumina Inc. FC-131-1024). For cell cultures, the human and mouse nuclei were freshly isolated as described in “Nuclei isolation of cell lines” and resuspended in 50 μl of 1x Nuclei Buffer (10X Genomics, PN-2000207). Then, we counted nuclei for each sample and added 5000 nuclei diluted in 20 μl of 1.25x Tagment DNA buffer to each well of the iTSM plate (see “Sample multiplexing” for details), except for the wells used to test the 10X reagents in which 5000 nuclei diluted in 5 μl of 1x Nuclei Buffer were added to a mixture of 7 μl of ATAC Buffer B (10X Genomics, PN-2000193) and 3 μl of barcoded Tn5. The plate layout and well IDs for each barnyard condition was shown in Fig. S1 and Table S6. The tagmentation was performed at 55 °C for 30 min on a thermocycler with a heated lid. To quench the Tn5 activity, we added a 2x Tagmentation Stop Buffer containing 40 mM EDTA (Invitrogen™, Cat. AM9260G) and 1mM Spermidine (Sigma-Aldrich, Cat. S0266-1G) to the transposition reactions at a 1:1 ratio and incubated the plate on ice for 15 min. We found that stopping the transposition reaction was unnecessary and thereby removed this step from our final txci-ATAC-seq protocol. All nuclei were pooled and centrifuged at 500 r.c.f for 10 min. After aspirating the supernatant, nuclei were resuspended in 400 μl 1x Nuclei Buffer and pelleted again. Then, we carefully removed the supernatant and resuspended nuclei in 30 μl 1x Nuclei Buffer. After quantification of nuclei with a hemocytometer, 75,000 nuclei were taken and diluted in 1x Nuclei Buffer to make a total volume of 15 μl, which underwent the standard 10X Chromium Next GEM protocol (v1.1, Document No. CG000209 Rev D from Steps 2 to 4) except following steps. For Sample Index PCR (step 4.1), we substituted the Single Index N Set A with a 25 μM i7 TruSeq primer and added 2.5 μl of customized i7 primer (Table S7) to each 10X library followed by performing 8 cycles of PCR amplification. The resulting library was sequenced on a NextSeq 550 Platform (Illumina Inc.) using a Mid Output Kit with the following cycles: Read 1, 50 cycles; i7 index, 8 cycles; i5 index, 16 cycles; Read 2, 77 cycles.

#### Blocking barcode-swapping

Flash-frozen (human and mouse lung samples and human cell line, see “Nuclei isolation” for details) and fresh nuclei (mouse cell line, see “Nuclei isolation” for details) were used to test the efficiency of strategies to block barcode-swapping. The flash-frozen nuclei were thawed, washed, and filtered following the procedures described in the “Coupling barcoded transposition with standard 10X protocol” section. Both flash-frozen and freshly isolated nuclei were resuspended in 100 μl of PBS containing 0.04% BSA (PBSB) and quantified using a hemocytometer (See “Nuclei isolation of cell lines” for details). After counting, the nuclei were diluted in PBSB to a concentration of 2,857 per μl (20,000 nuclei per well in 7 μl) and then mixed with a Tagmentation buffer solution (TBS, which was modified from the Omni protocol [10]) followed by transferring to the iTSM plate (see “Sample multiplexing” for details). Each 13 μl of TBS contains 12.5 μl of Illumina Tagment DNA Buffer, 0.25 μl of 1% Digitonin in DMSO (Promega (2%), Cat. PRG9441), and 0.25 μl of 10% Tween-20 (Bio-Rad, Cat. 1610781) in nuclease-free water. The barcoded transposition reaction was performed at 37 °C for 30 min on a thermocycler with a heated lid at 47 °C. Each blocking condition was assigned to 8 columns leading to a total of two 96-well plates for all three conditions. The plate layout and well IDs for each barnyard design in each blocking condition were shown in Fig. S2a and Table S6. After tagmentation, the nuclei used to test the Decoy DNA were transferred to a new 96-well plate with a multi-channel pipette, and 2.5 μl of 50 μM duplex DNA (see Table S8 for the oligo sequence) was added to each well followed by incubating at 55 °C for 10 min. Then, we added the 2x Tagmentation Stop Buffer (see “Coupling barcoded transposition with standard 10X protocol” for details) to the transposition reactions at a 1:1 ratio for all three blocking conditions and incubated the plates on ice for 15 min. Subsequently, the nuclei from the same blocking condition were pooled together and pelleted at 500 r.c.f for 10 min at 4 °C. After removal of supernatant from each tube, the nuclei were washed with 500 μl of 1x Nuclei Buffer (10X Genomics, PN-2000207) with centrifugation of 500 r.c.f for 5 min at 4 °C and resuspended in 25 μl of 1x Nuclei Buffer. Then, we counted nuclei with Trypan blue on a hemocytometer and diluted 100,000 nuclei in 1x Nuclei Buffer to make a total of 15 μl for each blocking condition. The resulting 3 aliquots of nuclei were run on separate lanes of the 10X as per the manufacturer’s instructions (10X Chromium Next GEM Single Cell ATAC protocol v1.1, Document No. CG000209 Rev D) with the following modifications. During GEM Generation and Barcoding (Step 2.1a), the nuclei dedicated to evaluating the Blocking oligo were mixed with the Master Mix supplemented with 2.5 μl of 100 μM DNA oligo incorporating an inverted dT at the 3’- end (see Table S8 for the oligo sequence); and the nuclei dedicated to testing the SBS primer were mixed with the Master Mix supplemented with 2.5 μl of 25 μM full SBS primer (Table S8) for in-droplet exponential amplification. After GEM PCR (Step 2.5a), a 10 μl PCR product (10% GEM) was slowly aspirated and transferred to a new PCR tube and subjected to Post GEM Incubation Cleanup in parallel with the 90% sample. Following cleanup, we performed the Sample Index PCR on the 10% sample (step 4.1) by supplementing the PCR mixes of SBS primer, Decoy DNA, and Blocking oligo with 2.5 μl of 25 μM barcoded i7 TruSeq primer (Table S7), which was used to replace the Single Index N Set A. The PCR mixes were amplified and monitored on a Bio-Rad CFX Connect Real-time cycler. The amplification was stopped when it appeared to be leveling off (i.e., the SBS primer was stopped at 4 cycles; the Decoy DNA and Blocking oligo were stopped at 15 cycles). To monitor the relative efficiencies of amplification in our initial test, we ended up introducing 2 different barcoded SBS primers in the SBS condition: one barcode was used for in-droplet amplification and another barcode was used for final library sample indexing. Both barcodes were assigned to thousands of reads per cell, indicating that both reactions were working. However, the theoretical expectation for the ratio between the two barcodes was 1/16 (because the second primer was used for 4 cycles of PCR). When we examined the ratio in our actual data, it was consistently ∼⅓, indicating that the sample index amplification is not perfectly efficient (Fig. S16). Therefore, in subsequent experiments using lung and liver tissues, we reduced the in-droplet PCR to 8 cycles and added an additional cycle of PCR for sample indexing. The resulting libraries with 10% GEM were pooled together with a library from an unrelated experiment to balance nucleotide diversity through the fixed sequence at the Tn5MErev region in Read 2, and then sequenced on a NextSeq 550 Platform (Illumina Inc.) using a Mid Output Kit with the following cycles: Read 1, 50 cycles; i7 index, 10 cycles; i5 index, 16 cycles; Read 2, 92 cycles. While 8 cycles in i7 index and 77 cycles in Read 2 were sufficient for the libraries generated in this study, we ran 10 and 92 cycles for those two steps, respectively, to accommodate the other library.

### txci-ATAC-seq using brain tissue samples

Tagmentation plates were prepared by the combination of 1430 μl of TBS with 770 μl nuclei solution. The TBS recipe was described in “Blocking barcode-swapping”, but a different version of Digitonin (Bivision 2082-1) was used here. This solution was mixed briefly on ice. 20 μl of the mixture was placed into the 96-well iTSM plate (see “Sample multiplexing” for details). Tagmentation was performed at 37 °C for 60 minutes on a 300 r.c.f Eppendorf ThermoMixer with a lid heated to 65 °C. Following this incubation, plate temperature was brought down with a 5-minute incubation on ice to stop the reaction. Tagmented nuclei were then pooled into a single 15 ml conical tube. 5 ml of tagmentation wash buffer (TMG) was prepared consisting of a final concentration of 10 mM Tris Acetate pH 7.5 (Sigma 93352 and Sigma A6283, respectively), 5 mM MgAcetate (Sigma M5661), and 10% (v/v) glycerol (Sigma G5516), diluted in PCR grade water. 1 ml of TMG was added on top of the chilled tagmented nuclei. Nuclei were pelleted at 500 r.c.f for 10 minutes at 4 °C. Most of the supernatant was removed with care not to disturb the pellet. Then 500 μl of TMG was added to the pellet and the tube was once again spun at 500 r.c.f. for 5 minutes at 4 °C. 490 μl was removed leading to a low volume of concentrated nuclei. Loading buffer was prepared consisting of 10% (v/v) glycerol, 20 mM NaCl, 10 mM Tris-Cl pH 7.5 (Life technologies AM9855), 0.02 mM EDTA (Fisher Scientific AM9260G), 0.2 mM DTT (VWR 97061-340), and 0.2x (v/v) TB1 (Illumina Inc.). The nuclear pellet was resuspended with an additional 30 μl of loading buffer. An aliquot of 2 μl of sample was diluted 20-50X and quantified with Trypan Blue on a hemocytometer. Depending on the experiment, a 14 μl nuclei solution containing the desired amount of nuclei in the loading buffer was then combined with 1 μl of 75 μM short SBS oligo (Table S8).

The 10X Chromium was then run with the custom nuclei solution as per the manufacturer’s instructions (10x Document CG000209 Rev D) with the following adaptations. In step 2.4e during GEM aspiration and transfer, 100 μl GEM volume was split into two tubes, with one receiving 10 μl and the other 90 μl (henceforth referred to as 10% and 90% samples). In step 2.5.a, GEM incubation cycles were limited to 6. For Pre-PCR wash elution (Step 3.2.j) the 10% sample was eluted in 8.5 μl whereas the 90% sample was eluted in 32.5 μl. For step 3.2.n, the 10% sample had 8 μl transferred to a new strip, while the 90% sample had 32 μl transferred to a new strip. At step 4.1.b, the sample Index PCR mix was split with 11.5 μl and 46 μl being combined with the 10% and 90% samples, respectively. For step 4.1.c, 1 μl and 2 μl of a 10 μM i7 TruSeq primer was used, respectively. For step 4.1.d, 8 and 7 PCR cycles were used, respectively. Libraries were then checked for quality and quantified by Qubit DNA HS assay (Agilent Q32851) and Tapestation D5000 (Agilent 5067-5589) following the manufacturer’s instructions. Libraries were then diluted and sequenced on a NextSeq 500 Mid flow cell or a NovaSeq 6000 S4 flow cell (Illumina Inc.).

### txci-ATAC-seq using human lung, mouse lung, and mouse liver tissue samples

Flash-frozen nuclei isolated from human lung, mouse lung, and mouse liver tissues were thawed, washed, and filtered following the procedures described in “Coupling barcoded transposition with standard 10X protocol”, and then resuspended in 150 μl PBSB (PBS containing 0.04% BSA). To count nuclei, we added 1.5 μl of 300 μM DAPI to 150 μl of PBSB containing nuclei for a final concentration of 3 μM DAPI, and incubated the nuclei on ice for 5 min. Then, we mixed nuclei with 2x Omni TD buffer in a 1:1 ratio and loaded 10 μl on a Countess Cell Counting Chamber Slide to count the nuclei with Countess II Automated Cell Counter.

After counting nuclei, we diluted the samples with PBSB to a concentration of 2,857 per μl and mixed 7 μl of nuclei solution (20,000 nuclei) with 13 μl of TBS (see “Blocking barcode-swapping” for details) for each well. This 20 μl nuclei/transposition mixture was then added to each well of the iTSM plate pre-loaded with 5 μl of barcoded Tn5 per well (see “Sample multiplexing” for details) to make a total volume of 25 μl reaction per well. As shown in Fig. 4a, 20,000 mouse nuclei were added to each well from rows A to F. But for rows G and H, 10,000 mouse nuclei were mixed with 10,000 human nuclei and then transferred to each well to estimate the empirical collision rate for each sample. The well IDs for different tissue mixtures were specified in Table S6. After loading nuclei, the iTSM plate was sealed and briefly shaken at 1000 rpm for 1 min on a pre-chilled thermomixer. The barcoded transposition was performed at 37 °C for 1 hour on a thermocycler with a heated lid at 47 °C. At the end of incubation, the plate was briefly centrifuged at 500 r.c.f for 10 seconds and then chilled on ice for 5 min to stop the transposition reaction. After quenching enzyme activity, the nuclei were pooled into a 12-tube strip and then transferred to a 15 ml conical tube preloaded with 400 μl tagmentation washing buffer (TMG, which contains 10mM Tris Acetate pH 7.8 (Boston BioProducts, Cat. BB-2412), 5mM Magnesium acetate (Sigma, Cat. 63052-100ML) and 10% (v/v) glycerol (VWR, Cat. RC3290-32) diluted in nuclease-free water). Subsequently, we added 50 μl/well of TMG to the first row of the plate and pipetted them throughout the whole plate to wash out the residual nuclei remaining in the plate. After washing the last row of the plate, the TMG was transferred to the same conical tube that was used to collect the barcoded nuclei. The pooled nuclei were then centrifuged at 500 r.c.f for 10 min in a pre-chilled swinging-bucket centrifuge at 4 °C. After aspirating the supernatant, the nuclei were resuspended in 500 μl TMG and then transferred to a 1.5 ml LoBind tube through a 40 μm Flowmi Cell strainer. The nuclei suspension was then centrifuged at 500 r.c.f for 5 min in a pre-chilled fixed-angle centrifuge at 4 °C. After centrifugation, 400 μl of supernatant was removed. The 100 μl of supernatant left from the first aspiration was then carefully removed by pipetting with a P200 pipette tip to avoid disturbing the nuclei pellet. The nuclei were resuspended with 30 μl of loading buffer containing 1x TB1 (Illumina Inc.), 1x standard storage buffer (Illumina Inc.), and 5 μM short SBS oligo (a final concentration of 1 μM SBS after resuspending nuclei in 10X Master Mix; see Table S8 for the oligo sequence), and then counted with a hemocytometer (see “nuclei isolation of cell lines” for details). After counting, the volume of solution containing the appropriate number of nuclei was taken and diluted with the loading buffer to make a total volume of 15 μl, which was used as an input into the 10x Chromium Controller. The GEM generation, Barcoding, and Post GEM Incubation Cleanup were performed following Steps 2 and 3 described in the 10X Chromium Next GEM Single Cell ATAC protocol (v1.1, Document No. CG000209 Rev D) except for Step 2.5, in which 8 cycles were used for GEM incubation. For Sample Index PCR (step 4.1), we substituted the Single Index N Set A (10X Genomics) with 25 μM i7 TruSeq primer containing an 8 bp custom barcode (Table S7) and added 2.5 μl of customized i7 primer to each 10X library. The PCR was performed following the 10X protocol shown in Step 4.1 but with 5 total cycles. The Double Sided Size Selection was then conducted as described in Step 4.2 shown in the 10X protocol. Following the size selection, the txci-ATAC-seq libraries were quantified by Qubit 1X dsDNA HS Assay Kit (Invitrogen, Cat. Q33231) and run on a 6% PAGE gel to check the library quality. To balance nucleotide diversity of the fixed sequence at the Tn5MErev region in Read 2, we pooled these libraries with 5% of bulk ATAC libraries (from an unrelated experiment) and sequenced them on a NextSeq 550 Sequencer (Illumina Inc.) using a High Output Kit with following cycles: Read 1, 51 cycles; i7 index, 10 cycles; i5 index, 16 cycles; Read 2, 78 cycles. The txci-ATAC-seq library only has 8 bp of i7 barcode, but we ran 10 cycles in i7 index to accommodate the barcode length of the bulk ATAC libraries. In cases where txci-ATAC-seq libraries are sequenced alone, we recommend either spiking in an appropriate amount of PhiX as per the manufacturer’s instruction or performing dark cycles for the cycles from 9 to 27 in Read 2.

### Fast-txci-ATAC-seq

To perform txci-ATAC-seq directly on frozen nuclei, the nuclei isolated from WT and CC16^-/-^ mouse lungs (see “Nuclei isolation of human lung, mouse lung, and mouse liver tissue” for details) were diluted in NFB (see Nuclei isolation of cell lines) at 3,175 nuclei/μl. For each sample, 6.3 μl of diluted nuclei (20,000 nuclei) were added to each well of an 8-tube strip for a total of 8 wells. Then, the nuclei were flash-frozen in liquid nitrogen and transferred to -80 °C for storage. The paired WT and CC16^-/-^ samples were processed together but each pair was processed on a separate day. When performing Fast-txci-ATAC-seq, the nuclei flash-frozen in the tube strips were thawed on ice and 13.7 μl of transposition buffer (which contains 12.5 μl of 2X Illumina Tagment DNA Buffer, 0.7 μl of 10X PBS, 0.25 μl of 1% Digitonin, 0.25 μl of 10% Tween-20) was added to each well containing nuclei followed by adding 5 μl of 500 nM pre-indexed Tn5 transposase per well. Then, the barcoded transposition reaction was performed on all 6 samples simultaneously by incubating at 37 °C for 60 min. Since each sample was distributed into 8 wells, a total of 48 Tn5 barcodes were used. As described above in the txci-ATAC-seq protocol, the barcoded nuclei were then cooled down on ice, pooled, washed, and loaded on the 10x Chromium Controller with either 50,000 or 100,000 nuclei in a lane.

### Data processing and analysis

Raw code for the brain analysis is available at https://mulqueenr.github.io/scidrop/. Raw code for the cell line and lung/liver datasets is available at https://github.com/cusanovichlab/scidropatac. The specific programs (and their version) used in data analyses were as follows: bcl2fastq (v2.19.0 for brain analysis and v2.20.0.422 for the other samples, Illumina Inc.), Trimmomatic (v0.36) [59], SAMtools and tabix (v1.7 for brain analysis and v1.10 for the other samples) [60,61], BWA-MEM (v0.7.15-r1140) [62], Bowtie2 (v2.4.1) [63], Perl (v5.16.3) [64], MACS2 (v.2.2.7.1 for brain analysis and v2.1.2 for the other samples) [65], bedtools (v2.28.0) [66], Python (2.7.13 [67] and 3.6.7 [68]), PyPy (5.10.0), pybedtools (0.7.10) [69], R (v4.1.1) [70], cisTopic (v0.3.0) [12], Cicero (v1.3.4.10) [15], Signac (v1.0.0 for brain analysis and v1.5.0 for the other samples) [16], Presto (v1.0.0) [21], chromVAR (v1.16.0) [22], Seurat (v4.1.0) [28], uwot (v0.1.8) [71], Harmony (v1.0) [72], irlba (v2.3.5) [73], mclust (v5.4.9) [74], edgeR (v3.40.0) [75], rGREAT (v2.0.2) [76], KEGGREST (v1.38.0) [77], BCFtools (v1.15.1) [78], GATK (4.3.0.0) [79], MOODS (1.9.4) [80], ggplot2 (v3.3.5) [81], and ComplexHeatmap (v2.5.5) [82].

### Computational analysis of brain samples

There were some deviations in the analysis of the brain samples, which are detailed below.

### Preprocessing for brain tissues

After sequencing, data was converted from bcl format to FastQ format using bcl2fastq with the following options “--with-failed-reads”, “--no-lane-splitting”, “--fastq-compression-level=9”, “-- create-fastq-for-index-reads”. Data were then demultiplexed, aligned, and de-duplicated using the in-house scitools pipeline [83]. Briefly, FastQ reads were assigned to their expected primer index sequence allowing for sequencing error (Hamming distance ≤2) and indexes were concatenated to form a “cellID”. Reads that could be assigned unambiguously to a cellID were then aligned to reference genomes. Paired reads were first aligned to a concatenated hybrid genome of hg38 and GRCm38 (“mm10”, Genome Reference Consortium Mouse Build 38 (GCA_000001635.2)) with BWA-MEM. Reads were then de-duplicated to remove PCR and optical duplicates by a Perl script aware of cellID, chromosome number, read start coordinate, read end coordinate, and strand. From there, the putative single-cells were distinguished from debris and error-generated cellIDs by both unique reads and percentage of unique reads.

### Barnyard analysis for brain tissues

With single-cell libraries distinguished, we next quantified contamination between nuclei during library generation. We calculated the read count of unique reads per cellID aligning to either human reference or mouse reference chromosomes (Fig. S3b). CellIDs with ≥90% of reads aligning to a single reference genome were considered bona fide single cells. Those not passing this filter were considered collisions. The collision rate was estimated using the equation in [11] to account for cryptic collisions (two cells from the same species). Bona fide single-cell cell IDs were then split from the original FastQ files to be aligned to the proper hg38 or mm10 genomes with BWA-MEM as described above. Human and mouse assigned cellIDs were then processed in parallel for the rest of the analysis. After alignment, reads were again de-duplicated to obtain proper estimates of library complexity.

### Dimensionality reduction for brain tissues

Pseudo-bulked data (agnostic of cellID) was then used to call read pile-ups or “peaks” via MACS2 with the option “--keep-dup all”. Narrowpeak bed files were then merged by overlap and extended to a minimum of 500 bp for a total of 350,261 peaks for human and 292,304 peaks for mouse. A scitools Perl script was then used to generate a sparse matrix of peaks × cellID to count the occurrence of reads within peak regions per cell. FRiP was calculated as the number of unique, usable reads per cell that are present within the peaks out of the total number of unique, usable reads for that cell for each peak bed file. Tabix-formatted files were generated using samtools and tabix. The count matrix and tabix files were then input into a SeuratObject for Signac processing. We performed LDA-based dimensionality reduction via cisTopic with 28 and 30 topics for human and mouse cells, respectively. The number of topics was selected after generating 25 separate models per species with topic counts of 5,10,20-30,40,50,55,60-70 and selecting the topic count using selectModel based on the second derivative of model perplexity. Cell clustering was performed with Signac “FindNeighbors” and “FindClusters” functions on the topic weight × cellID data frame. For the “FindClusters” function call, resolution was set to 0.01 and 0.02 for human and mouse samples, respectively. The respective topic weight × cellID was then projected into two-dimensional space via UMAP by the function “umap” in the uwot package. To check for putative doublets within species, we then ran scrublet analysis and removed the scrublet-identified doubles from further analysis [14]. A second iteration of sub-clustering was performed on each cluster to better ascertain cell type diversity. This was done as described above with the data subset to just the cells within the respective cluster for both cisTopic model building and UMAP projection. Resolution per subcluster was set *post hoc* based on cell separation in UMAP projection. CCANs and the resulting gene activities were generated through the Signac wrapper of Cicero. Genome-wide accessibility of known TF motifs was calculated per cell using the JASPAR database (release 8) [84] via chromVAR.

### Cell Type Identification for brain tissues

For cell type identification, we used previously existing single-cell RNA datasets of the human M1 cortex [85], and mouse whole cortex and hippocampus [86,87]. We applied the Signac label transfer strategy between the annotated single-cell RNA with our gene activity scores at the level of our sub-clustered cell groups. For cell type refinement, we plotted the average gene activity score per subcluster for a set of RNA-defined marker genes, as well as markers defined within our datasets on the gene activity scores using the Signac “FindMarkers” function as described above. Subcluster dendrograms were generated by using base R functions dist and hclust through running Z-scored average gene activity on internally-defined markers and based on “ward.D2” clustering of Euclidean distance. The resultant dendrogram was used for both pre-defined and internally defined marker sets. Results were plotted via ComplexHeatmap.

### TF marker ranking

TFs were ranked for specificity across sub-clusters, based on combined motif accessibility (generated through chromVAR) and gene activity (generated through cicero). AUC values were determined per cluster via the Wilcoxon test as reported by the “wilcoxauc” function in Presto. An average AUC of motif accessibility and gene activity was used for ranking TFs. A set of top 5 markers per sub-cluster was filtered for duplicates and then plotted via ComplexHeatmap.

### Comparison across scATAC-seq mouse brain datasets

FastQ files for sciATAC-seq, sciMAP, snATAC-seq, dscATAC-seq, and s3-ATAC-seq were downloaded via the SRA toolkit. 10X scATAC-seq v1 and v2 chemistries FastQ files were obtained through the 10X Genomics website. Files were then demultiplexed following the original author’s instructions to generate a scitools analogous cellID and were processed through the scitools pipeline as described above. Briefly, after alignment to a consistent mouse reference genome (GRCm38), files were treated to de-duplication in parallel before merging. For each dataset, cellIDs were filtered to those with at least 1,000 unique reads, and then merged into a single bam file. Peaks were called as previously described, resulting in 344,258 regions of accessibility. Per cell FRiP was calculated using this peak set. TSS enrichment values were calculated for all cells using the method established by the ENCODE project (https://www.encodeproject.org/data-standards/terms/enrichment), whereby the aggregate distribution of reads ±1,000 bp centered on the set of TSSs generates 100 bp windows at the flanks of the distribution as the background and then the maximum window centered on the TSS is used to calculate the fold-enrichment over the outer flanking windows. Signac was then used to generate a SeuratObject as described above, and data underwent dimensionality reduction after integration using Harmony, with argument “nclust=14”. A UMAP was plotted as described above with cellIDs colored in relation to the technology used, and the original author-assigned cell type.

### Computational analysis of human lung, mouse lung, and mouse liver tissue samples

#### Preprocessing

Fastq files were generated using bcl2fastq with the following options: “--ignore-missing-bcls”, “-- no-lane-splitting”, and “--create-fastq-for-index-reads”. Then, we modified the fastq files by attaching the first 8 bp (Tn5 barcodes) of Read 2 to the header and removing the first 27 bp (8 bp of Tn5 barcodes + 19 bp of Tn5 mosaic end) from Read 2 with a custom python script. Barcodes that did not perfectly match any of the expected barcodes were converted to the closest matching barcode if the edit distance was no greater than 2. Barcodes matching more than 1 expected barcode after correction were removed. After barcode correction, we demultiplexed samples based on a combination of Tn5 barcodes and i7 sample indices and generated a combined barcode for each read by concatenating the i7 sample index, 10X bead barcode, and Tn5 barcode. Next, we removed the sequence adaptors and low-quality reads using trimmomatic with following parameters: “LEADING:3; TRAILING:3; SLIDINGWINDOW:4:10; MINLEN:20” and then mapped the trimmed reads to a hybrid hg38/mm10 reference genome using Bowtie2 with a maximum fragment length of 2000 pb (-X 2000) and 1 base trimmed from the 3’ end of each read (−3 1). Following mapping, only the reads confidently (MAPQ ≥ 10) aligned to the assembled nuclear chromosomes and in proper pairs (determined by “-f3” and “-F12” options in SAMtools) were preserved for downstream analysis. To eliminate PCR duplicates, we removed all fragments that possessed the same combined barcode and identical start and end coordinates, keeping a random representative read for each end of the molecule using a custom script.

#### Peak calling

The deduplicated bed files were used to call peaks with MACS2, considering a 200 bp window centered on the read start using the parameters ‘--nomodel --keep-dup all --extsize 200 --shift - 100’. Because each peak may have multiple summits and will therefore be listed multiple times in the resulting peak bed file, the peaks output from MACS2 were then merged into a single peak set for each sample using bedtools “merge”. The consolidated peaks were then intersected with the ENCODE blacklist (mm10 [88] or hg38 ENCFF356LFX) to remove signal-artifact regions using bedtools “intersect” with “-v” option.

#### Calculation of ATAC-seq QC metrics

*FRiDHS.* The FRiDHS score was determined using orthogonal peak references identified in DNase-seq data. The GM12878 DHS peaks combined the 2 replicates of narrowPeak-formatted files obtained from the ENCODE consortium (ENCSR000EMT). The CH12.LX DHS peaks combined the 2 replicates of narrowPeak-formatted files obtained from the ENCODE consortium (ENCSR000CMQ). The mouse lung DHS peaks combined the 3 replicates of narrowPeak-formatted files obtained from the ENCODE consortium (ENCSR000CNM). The mouse liver DHS peaks combined the 14 replicates of narrowPeak-formatted files obtained from the ENCODE consortium (ENCSR000CNI). The human lung DHS peaks combined the narrowPeak-formatted files obtained from 2 separate DNase-seq data but from the same individual (ENCODE Donor Accession: ENCDO845WKR; ENCODE Experiment Accession: ENCSR164WOF and ENCSR058VBM). The overlapping peaks between replicate bed files were consolidated using bedtools “merge”, and the peaks overlapped with ENCODE blacklist (mm10 [88] or hg38 ENCFF356LFX) were removed using bedtools “intersect” with “-v” option. We removed the reads mapping to the non-nuclear genome and performed deduplication before calculating FRiDHS. The reads overlapping the DHS peak reference were counted using the “BedTool.intersect” function from pybedtools with “u=True”.

*TSS enrichment*. The human and mouse TSS coordinates were obtained from the Gencode human reference v39 [89] and Gencode mouse reference vM23 [90], respectively. To build TSS references, we first collected the most upstream base (accounting for strand) of each transcript using a custom R script, and then only the TSSs of gene types and transcript types listing the following terms were included: “protein_coding”, “lncRNA”, “IG_C_gene”, “IG_D_gene”, “IG_J_gene”, “IG_LV_gene”, “IG_V_gene”, “IG_V_pseudogene”, “IG_J_pseudogene”, “IG_C_pseudogene”, “TR_C_gene”, “TR_D_gene”, “TR_J_gene”, “TR_V_gene”, “TR_V_pseudogene”, “TR_J_pseudogene”. We also excluded transcripts with a tag of “readthrough_transcript” or “PAR”. These filters were similar to the filtering strategy used by the 10X single-cell ATAC-seq pipeline [91]. The TSS enrichment score for each cell was calculated using the TSSEnrichment function in the Signac package.

*Estimated complexity*. The nuclear genome mapped reads and deduplicated reads were used to estimate the complexity for each cell using the same calculation as Picard [92] implemented in R.

### Collision rate estimation

For each combined barcode, we quantified the number of deduplicated reads mapping to the human and mouse genome and filtered out the combined barcodes with fewer than 1,000 total reads. The collision barcodes were determined as the cell barcodes that had more than 10% of reads aligned to the minor genome. Since the cell doublets can be generated by either two cells from the same species or cells from distinct species, the observed collisions only reflect approximately half of the collision events that in fact occur in the experiment. To this end, we estimated the actual collision rate using the equation in [11].

### Dimensionality reduction and clustering

An iterative peak-calling strategy was used to perform dimensionality reduction and cluster cells. The first round of clustering was performed with a pseudo-bulk peak reference, which was identified by calling peaks on deduplicate reads from identified cells (≥ 1,000 reads). Then, a binarized peak (column) by cell (row) matrix was generated by scoring the peaks defined in the previous step for overlap with reads from each cell. The low complexity cells and features were removed using Signac “CreateChromatinAssay” function by setting “min.cells = 50 and min.features = 200” for mouse samples and setting “min.cells = 15 and min.features = 200” for human samples followed by filtering out the cells considered as outliers for QC metrics (DHS region reads > 20,000, FRiDHS < 0.2 and TSS enrichment score < 2). Potential cell doublets were identified by performing a modified version of the Scrublet workflow [14] on each txci-ATAC-seq library separately. In brief, we transformed the filtered cell/peak matrix with the term-frequency inverse-document-frequency (TF-IDF) algorithm by computing log(TF×IDF) as described in [93] and then calculated the first 30 components for PCA using the irlba R package. Simulated cell doublets were created by randomly sampling 50% of observed cells from the original matrix and summing them with another 50% of randomly sampled cells. The matrix of simulated doublets was then binarized and transformed with the same TF-IDF implementation. Subsequently, we projected the transformed doublets into the PCA space generated by the observed data and performed L2-normalization on the resulting matrix including both observed and simulated cells with Seurat “L2Dim” function. The L2-normalized reduction was then used to compute the fraction of simulated doublet neighbors for each cell using Seurat “FindNeighbors” function with dimensions 2 to 30 and setting “k.param” as 129 (mouse lung nuclei from the 100,000 lane), 166 (mouse lung nuclei from the 200,000 lane), 120 (mouse liver nuclei from the 100,000 lane), 147 (mouse liver nuclei from the 200,000 lane), 62 (human lung nuclei from the 100,000 lane), and 74 (human lung nuclei from the 200,000 lane). We derived the “k.param” values using the k_adj_ equation in Scrublet. Finally, a doublet score was calculated for each cell using the appropriate equations described in Scrublet.

Assuming a bimodal distribution, a threshold for doublet scores was calculated with the simulated cells by identifying the boundary between the doublets incorporating highly similar cells (“embedded”) and the doublets of dissimilar cells (“neotypic”) using the mclust R package [74] with less than 5% uncertainty that the doublets were classified into the “neotypic” category. After removing the doublets detected, we computed the latent semantic indexing (LSI) matrix by running singular value decomposition (SVD) on the TF-IDF normalized matrix using Signac and then clustered cells using Seurat “FindNeighbors” function with the dimensions of reduction from 2 to 30 followed by Seurat “FindClusters” implementing the SLM algorithm with default resolution. For tissues with replicates, the LSI matrix was integrated by individual with Harmony [72] prior to cell clustering.

Each cell cluster from this first round of clustering was then used to identify peaks independently, and all cluster peaks were merged into a single reference set. Subsequently, a second round of clustering was performed using this updated peak set. With the same workflow, we used in the first round of clustering, a binarized count matrix generated with cluster-identified peaks was created and used to perform normalization, dimension reduction, integration (for tissues with replicates), and clustering, except that the resolution parameter used to determine the community size was set differently for each tissue (i.e., 0.8, 0.2 and 0.3 were used for mouse lung tissue, mouse liver tissue, and human lung tissue, respectively). Regarding liver samples, we decided to consolidate clusters 0, 7, 8, and 9 because of no visible separation between them in 2D UMAP space. For visualization purposes, the data was projected into a two-dimensional space via Seurat “RunUMAP” function with 30 dimensions (excluding the first component, which represented the sequencing depth).

### Cell type annotation

The cell types associated with each cluster were predicted by label transfer using publicly available sc/snRNA-seq and sci-ATAC-seq data. Only the cell types including at least 50 cells in the reference dataset were used to infer the cell types in the query dataset. To annotate cell types with transcriptome data, we used previously published data from steady state mouse liver, mouse lung, and healthy human lung samples to construct an “integrated” reference for each tissue in each species using the Seurat scRNA-seq integration pipeline. In all cases, the 5000 most variable genes across reference samples were selected to find integration “anchors”. The mouse lung reference was built by integrating three samples (a scRNA-seq sample and two replicate samples of snRNA-seq) from a single study [29]. The mouse liver reference was created by integrating samples generated with three different protocols (snRNA-seq and scRNA-seq using cells isolated via either *ex vivo* or *in vivo* enzymatic digestion method) from a single study as well (https://www.livercellatlas.org/download.php; [30]). The human lung reference was established by integrating two scRNA-seq datasets obtained from two independent studies [31,32]. After creating RNA-seq references, we estimated transcriptional activity across the genes selected for integration by quantifying the txci-ATAC-seq counts in both the 2 kb region upstream and the gene body of each gene using the Signac “GeneActivity” function. The prediction of cell type was then achieved by performing canonical correlation analysis on the gene activity scores calculated from ATAC-seq data along with the integrated scRNA-seq reference using Seurat “FindTransferAnchors” function followed by transferring annotations from reference to query cells using “TransferData” function in which the 2nd to 30th components of the LSI matrix calculated on ATAC-seq data was used to compute the weights of the local neighborhood of anchors.

For annotating cells with a chromatin reference, we downloaded the fastq files of mouse sci-ATAC-seq data from [3] (GEO accession number: Lung RepA, GSM3034631; Lung Rep B, GSM3034632; Liver, GSM3034630) and mapped them to the mm10 reference genome using Bowtie2. In terms of the human reference, the cell by bin (5kb) matrices were downloaded from 4 lung samples [33] (GEO accession number: GSE165659) and binarized prior to cell type prediction. To ensure that the same features were measured in the reference and query datasets, we summarized the reads from txci-ATAC-seq to either the peaks identified from the pseudo-bulk sci-ATAC-seq data (Mouse) or 5 kb genomic windows (human) for each query cell and only retained the features that were detected in at least 50 (mouse) or 15 (human) cells in both datasets. The label transfer was performed using Seurat “FindTransferAnchors” function with reference.reduction = “lsi” and reduction = “lsiproject” followed by “MapQuery” function with reference.reduction = “lsi”.

The final cell types were determined by applying a majority vote strategy to each cluster. For undetermined clusters or clusters with inconsistent labels between the RNA-seq and ATAC-seq reference-based predictions, the cell types were determined by screening the high activity genes for each cluster (identified using the Seurat “FindMarkers” function) for activity patterns consistent with gene expression either characterized in the scRNA-seq reference or (for mouse) collected in UCSC Tabula Muris [94,95]. The color palette used for cell types in the UMAPs was selected from colors available in the ArchR package [96].

### Identification of differential peaks

An edgeR-based pseudo-bulk method was used to identify differential peaks between WT and CC16^-/-^ mouse lungs for each cell type. To do so, we aggregated the reads for all cells from the same replicate in a cluster-wise manner, which resulted in 3 biological replicates for each genotype per cell type. The lowly accessible peaks in each differential test were filtered out using the “filterByExpr” function with default parameters followed by calculating normalization factors with “calcNormFactors()”. Then, we estimated dispersions using “estimateDisp()” with a design matrix and “robust = TRUE” and performed hypothesis testing using the quasi-likelihood F-test. The Benjamini and Hochberg (BH) method [97] was used to control the false discovery rate (FDR).

### Variant calling and filtering

The variant calling was performed jointly across all replicates from the same genotype using BCFtools with “bcftools mpileup” followed by “bcftools call” command. Only the SNVs that met the following criteria were used for downstream analyses: a Phred-scaled quality score (QUAL) of at least 20, a sum of read depth (DP) across all three replicates of at least 10, and the same genotype in at least two of three replicates.

### Motif analysis

The motif position frequency matrices obtained from the JASPAR database (version 2020) [98] were used for all motif analyses. For motif enrichment analysis, we applied the Signac “FindMotifs” function to all differentially accessible peaks per cell type to identify the enriched motifs using a GC-content-matched set of peaks created from the accessible peaks as a background. The multiple testing correction was performed with the BH procedure [97].

To identify the SNV-driven gains and losses in motif matching, we first generated alternative DNA sequences over the differentially accessible peaks from SNV hotspots (mm10 chr8:68,000,000-93,000,000 and mm10 chr19:16,500,000-26,500,000) for WT and CC16^-/-^ samples by replacing the reference bases at the variation sites with the hotspot SNVs identified in each genotype using GATK “FastaAlternateReferenceMaker” tool. Then, we matched the motifs against the alternative DNA sequences using the MOODS package with a p-value cutoff of 0.0001. To identify functional motifs capable of accounting for the chromatin accessibility changes, we tested for associations between the log_2_(fold-change) of differentially accessible peaks and the gain or loss of motifs in CC16^-/-^ background using the student’s *t-*test and controlling for multiple testing with the BH method [97]. When counting differences in accessibility that might be explained by specific motif-disrupting SNVs, we only considered the instances that exhibit a coherent change in chromatin accessibility with the overall motif effect to be explanatory (i.e., depending on whether gained/lost motifs are positively or negatively associated with peak accessibility, each of the potential explanatory instances for that motif also needs to display concordant increases or decreases in accessibility to be counted as explanatory).

### Functional analysis

The KEGG pathways enrichment analysis was performed by applying rGREAT on all differential peaks identified for each cell type using both binomial and hypergeometric tests. To control both tests, we used a previously implemented two-threshold approach [99] to define the significant pathways by requiring a stringent 10% FDR threshold for at least one test, but allowing for a more relaxed threshold (unadjusted p-value of 0.05) for the other test. The gene sets of KEGG pathways were retrieved using the KEGGREST package.

## Abbreviations

APB: ATAC-PBS Buffer
ATAC-seq: Assay for Transposase Accessible Chromatin using Sequencing
AUC: Area Under Curve
BH: Benjamini and Hochberg
BSA: Bovine Serum Albumin
CC16: Club Cell Secretory protein
CCANs: Cis-Coaccessibility Networks
COPD: Chronic Obstructive Pulmonary Disease
ES: Embryonic Stem
ETS-domain: E26 Transformation Specific-domain
FDR: False Discovery Rate
FRiDHS: Fraction of Reads in DNase I Hypersensitive Sites
FRiP: Fraction of Reads in Peaks
GEM: Gel Bead-In EMulsions
iTSM: pre-indexed Tn5 transposase per well
LHX2: LIM Homeobox 2
LHX6: LIM Homeobox 6
LSI: latent Semantic Indexing
NFB: Nuclei-Freezing Buffer
NFI: nuclear factor I
PBSB: PBS containing 0.04% BSA
PMI: Post-Mortem Interim
RSB: ATAC Resuspension Buffer
scATAC-seq: single-cell ATAC-seq
sci: single-cell combinatorial indexing
scRNA-seq: single-cell RNA-seq
SNP: Single Nucleotide Polymorphism
snRNA-seq: single-nucleus RNA-seq
SNV: Single Nucleotide Variants
SVD: Singular Value Decomposition
TF: Transcription Factors
TF-IDF: Term-Frequency Inverse-Document-Frequency
TMG: Tagmentation Wash Buffer
TSS: Transcription Start Site
txci-ATAC-seq: TenX(10X)-Compatible Combinatorial Indexing ATAC-seq
UMAP: Uniform Manifold Approximation and Projection
WT: Wild Type

## Declarations

### Ethics approval and consent to participate

All animal use was approved by the Institutional Animal Care and Use Committee of the University of Arizona or OHSU and adhered to the NIH Guide for the Care and Use of Laboratory Animals. The details of the animal study reported here are in compliance with ARRIVE guidelines.

Human brain tissue specimens were obtained from the Oregon Brain Bank under OHSU-approved IRB protocols. Human lung tissue specimens were obtained from the Arizona Donor Network with IRB-approved protocols.

### Consent for publication

Not applicable

### Availability of data and materials

The datasets generated during the current study will be made available from the Gene Expression Omnibus repository (https://www.ncbi.nlm.nih.gov/geo/).

### Authors’ contributions

H.Z., R.M.M., A.C.A., and D.A.C. conceived the study. H.Z. performed the experiments and data analysis for the txci-ATAC-seq of the cell line barnyard, human lung, naive mouse lung and liver, CC16 knockout lungs, as well as Fast-txci-ATAC-seq under the supervision of D.A.C. R.M.M. performed the experiments and data analysis for the txci-ATAC-seq of human and mouse brain under the supervision of A.C.A. N.I. and J.G.L. provided the WT and CC16 knockout lungs. D.O.F. and J.J.G. provided the naive mouse lung and liver samples. F.P. provided the human lung sample. N.I., J.G.L., D.O.F., J.J.G., and F.P. aided in the development of nuclei isolation protocol for human lung, mouse lung, and mouse liver samples. The paper was written by H.Z., R.M.M., A.C.A., and D.A.C. All authors read, edited, and approved the final manuscript.

## Acknowledgments

We thank J. Galina-Mehlman, R. Sprissler, and B. Fransway from the University of Arizona Genetics Core for their invaluable assistance. We thank C. Romanoski from the University of Arizona for helpful advice and insightful comments on the manuscript. We also would like to thank C. Thornton from Oregon Health & Science University for help in cell type identification of mouse and human brain tissues.

## Supplementary Figures

**Supplementary Figure 1.**
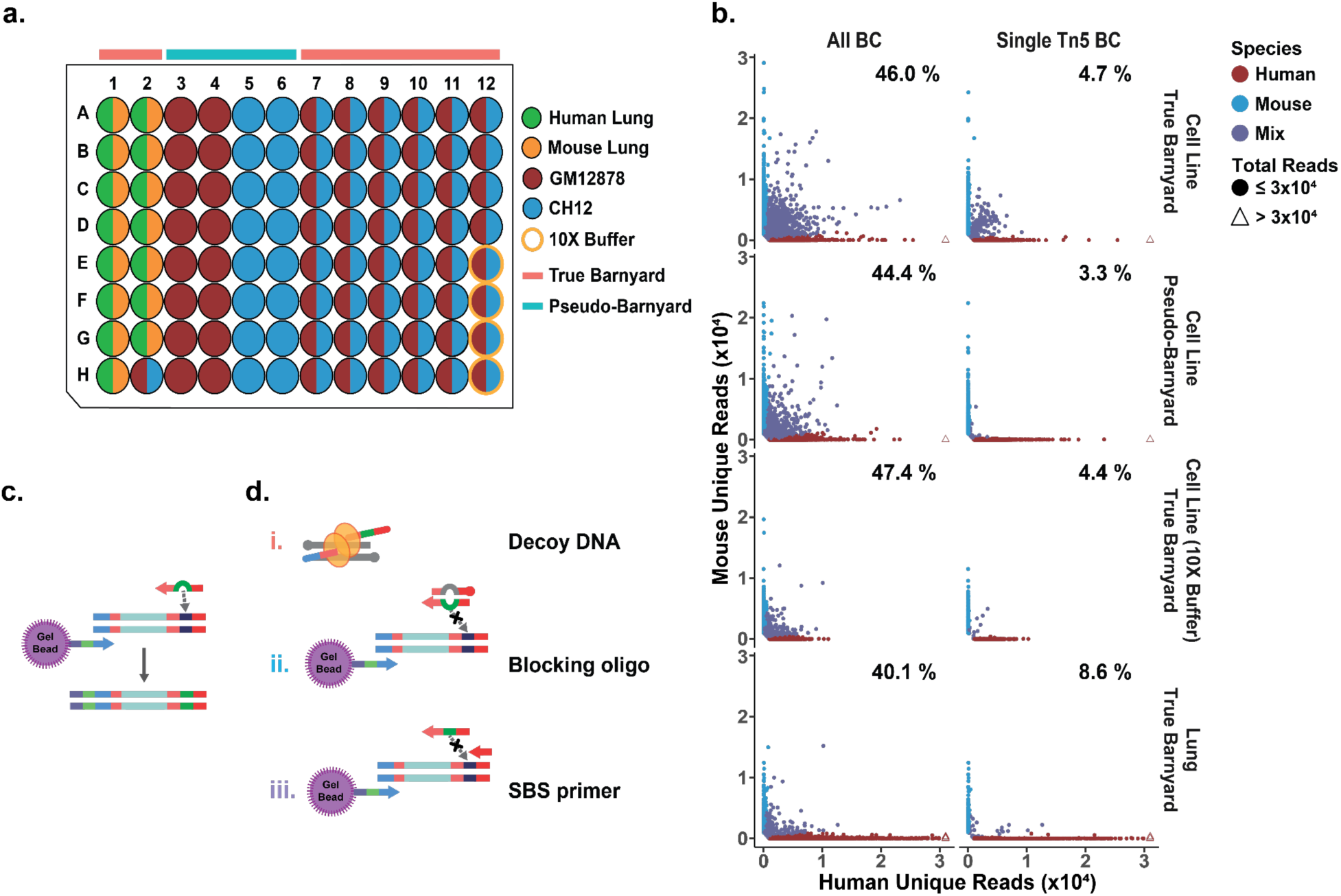
Collision rates of standard 10X protocol coupled with combinatorial indexing. a) Well assignment for each cell source and barnyard design. The wells with a mixture of species are shown as half-circles of two different colors corresponding to each species. The wells where tagmentation was done with the 10X ATAC buffer are highlighted by orange outer circles. b) The scatter plots showing the number of reads mapped to either the human or mouse genome for each barnyard design. The plots on the left-hand side include all cell barcodes, and the plots on the right-hand side only visualize the 10X bead barcodes associated with a single Tn5 barcode. The percentage represents the estimated collision rate for each plot. The x- and y-axes are capped at 3×10^4^ reads, and the cells with more than 3×10^4^ reads are denoted by triangles. c) the theoretical model of the process of Tn5 barcode swapping. d), the proposed strategies to block barcode swapping.

**Supplementary Figure 2.**
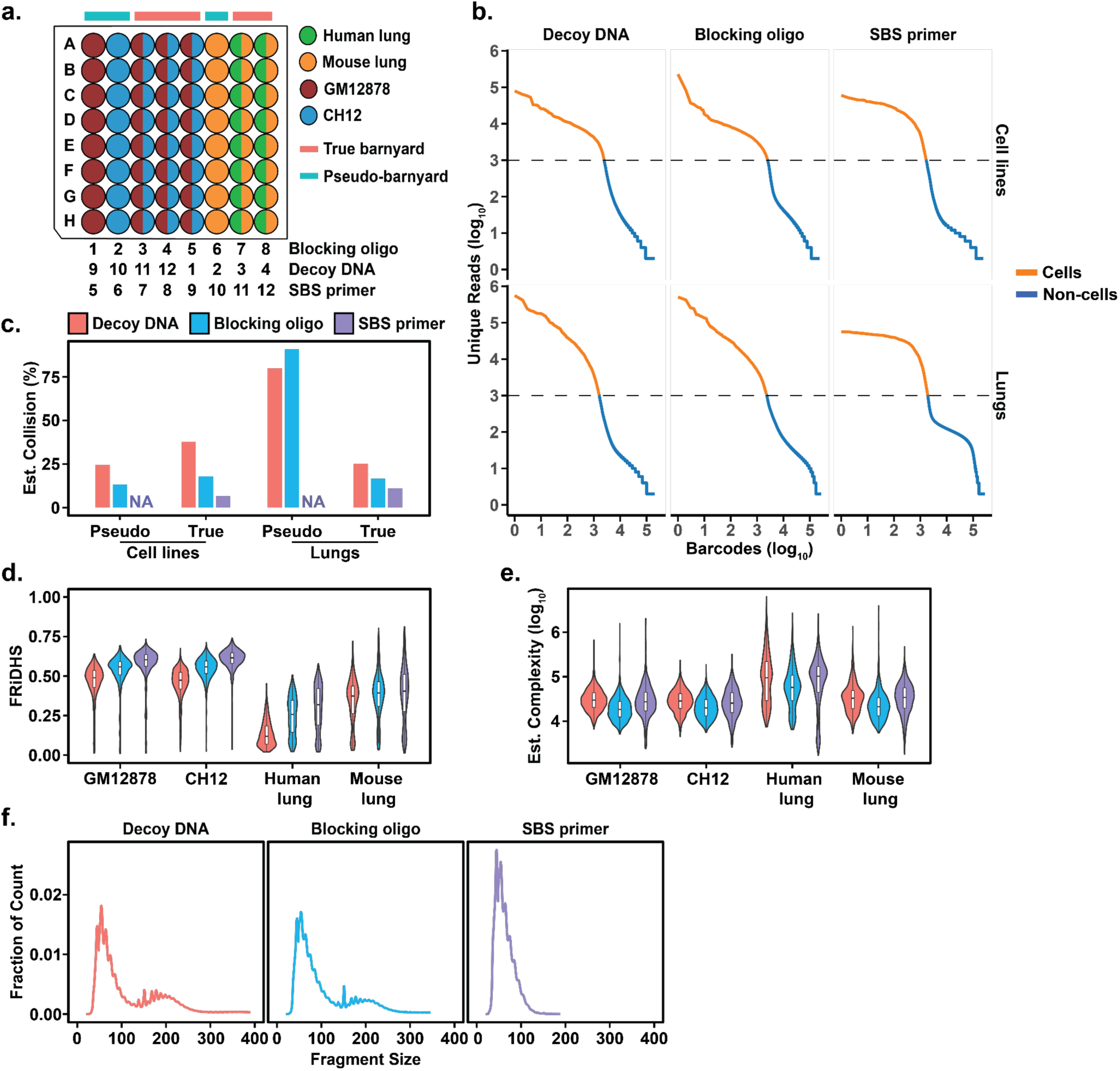
Exponential amplification during GEM PCR enables deconvolution of cells in the same droplet. a) Well assignments for the experiment testing the performance of each blocking strategy. Each blocking condition was allocated to ⅔ of the plate and a total of two 96-well plates were used to test all three conditions. The column ID assigned to each blocking condition is shown under the 96-well plate. b) “Knee” plots showing the separation between cell barcodes (orange line) and background barcodes (blue line) in either cell lines or lung tissues using different blocking methods. The dashed line indicates the threshold (1000 reads) used to identify cell barcodes. c) Comparison of estimated collision rate using different blocking strategies for each barnyard condition. The collision rate is not calculated for the pseudo-barnyard supplemented with SBS primer (positions labeled with NA) due to either no collision cells (cell line) or no human cells (pure mouse wells) observed, although one cell technically met the collision threshold (<90% of reads mapping to the major species) in the pure mouse wells (pseudo-barnyard). d-e) QC metrics of each blocking approach across different cell sources. The (d) FRiDHS and (e) estimated complexity (on a log_10_ scale) in each cell barcode are plotted for each strategy. The color legend is consistent with panel (c). f) The fragment size distribution of each blocking condition.

**Supplementary Figure 3.**
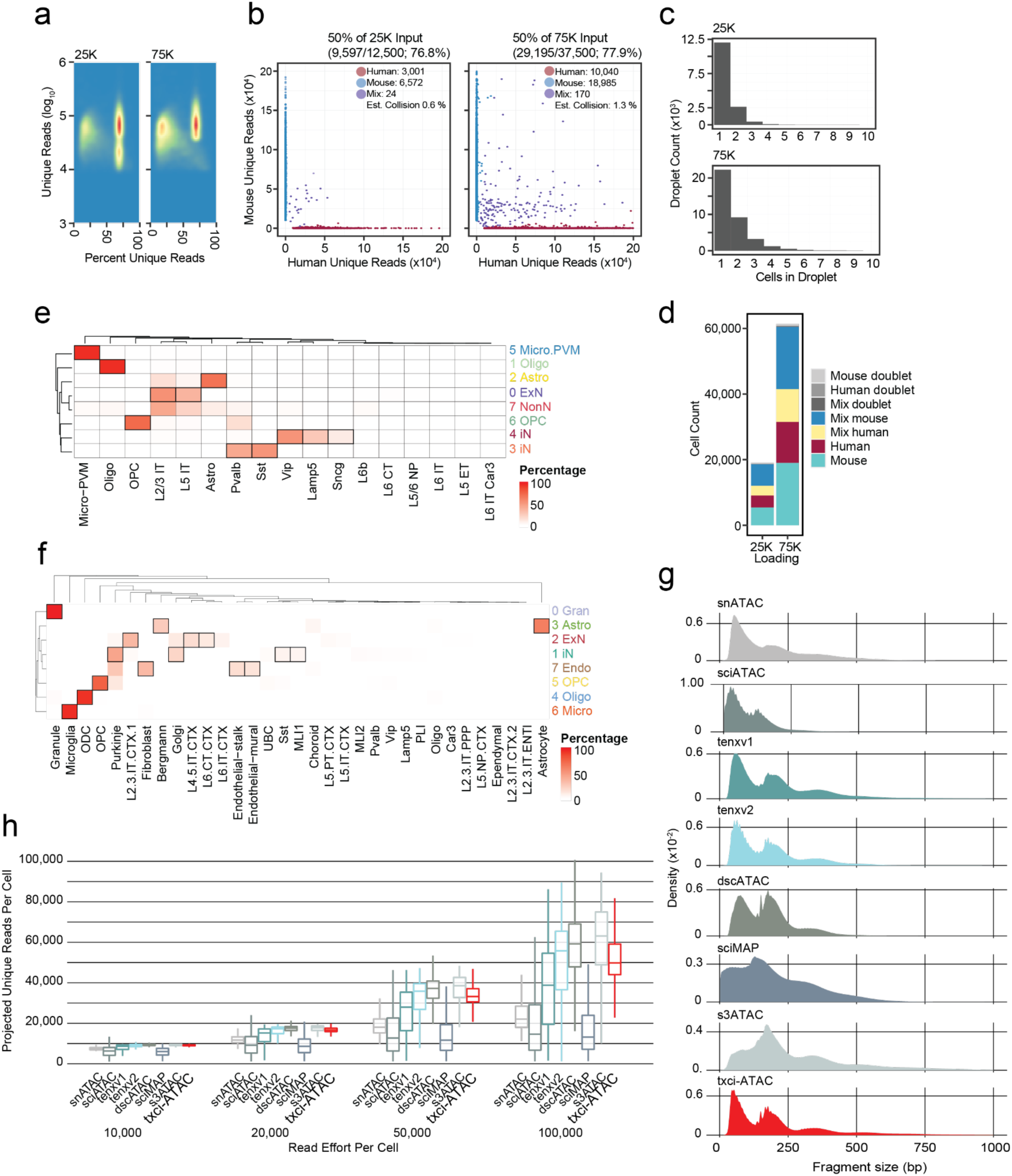
Evaluating the performance of txci-ATAC-seq on brain samples. a) Two-dimensional density map of cells passing initial read filters for percent unique reads (library saturation) and unique read counts. b) Mixed-species tagmentation wells were subject to alignment in both human and mouse reference genomes. c) Number of cells per droplet quantified on a histogram, showing a majority of droplets still contain only a single cell. d) Quantification of cells in 25,000 and 75,000 library pools. Conditions include doublets uncovered either through cross-species alignment (“mixed doublet”) or through a reduced dimension detection strategy (see Methods). Other cells passing these filters are colored by identified species and tagmentation conditions. e,f) Hierarchically clustered heatmap of cell identification in the human cortex (e) and mouse whole brain (f) samples using gene activity scores for label transfer. The Brain Map M1 Cortex RNA dataset was used to annotate human cells and the Brain Map mouse cortex and hippocampus RNA dataset and mouse cerebellum (GSE165371) were used to annotate mouse cells. Values reflect the percentage of cells per cluster with each label as its maximum predicted value. g) Density plots of fragment length for fragments ranging from 1-1000 bp per technology. h) Grouped boxplots of projected unique read count per sequencing effort for each cell by technology.

**Supplementary Figure 4.**
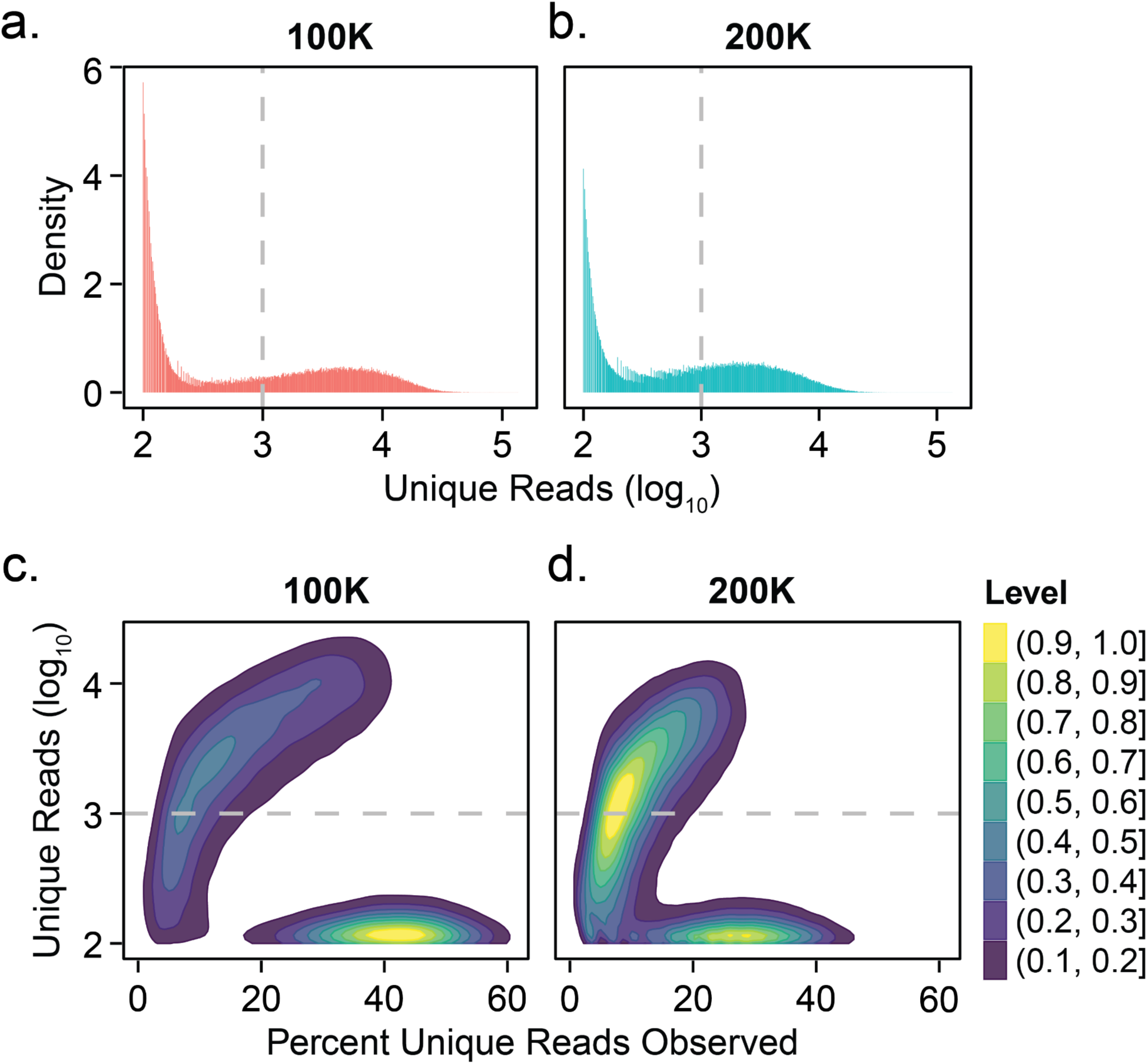
The sequencing depth of txci-ATAC-seq libraries loading 100,000 or 200,000 nuclei. a,b) Histograms showing the distribution of unique read counts (on a log_10_ scale) assigned to each possible barcode combination at the 100,000 (a) and 200,000 (b) nuclei loading inputs. The gray dashed line indicates the threshold (1000 reads) to identify a barcode as a cell. Barcode combinations with fewer than 100 total reads are not plotted. c,d) Contour plots showing the number of deduplicated reads against the estimated percent unique reads observed for each barcode combination at the 100,000 (c) and 200,000 (d) nuclei inputs. The estimated percent of unique reads observed was calculated by dividing the number of observed unique reads by the estimated complexity for each barcode combination. The color legend shows the normalized barcode density (as calculated in ggplot2) scaled from high (yellow) to low (blue). The gray dashed line indicates the threshold to call a cell barcode. The barcode combinations with fewer than 100 total reads are not shown on the plot.

**Supplementary Figure 5.**
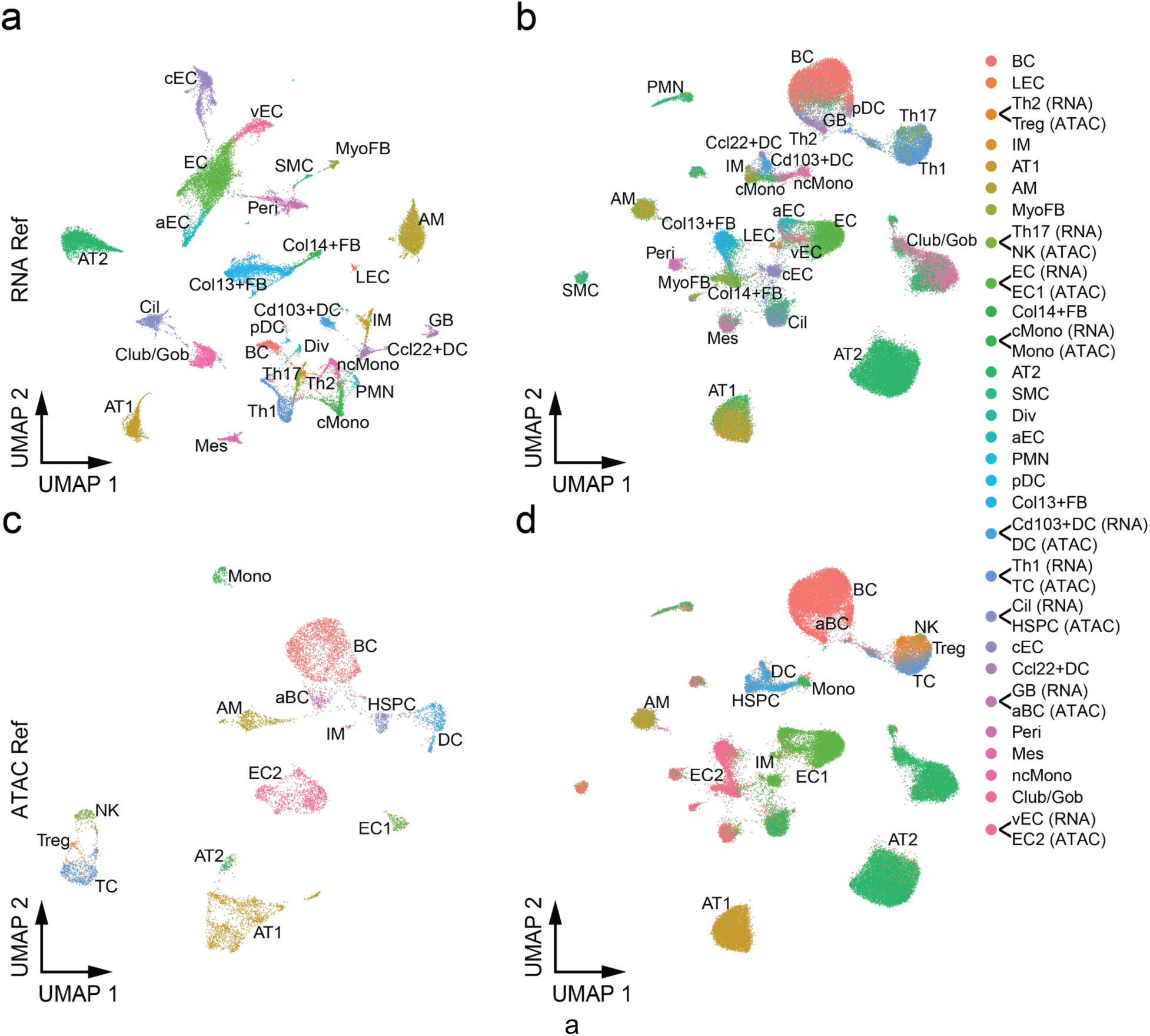
Cell type annotation of mouse lung samples with label transfer. a) UMAP of sc/snRNA-seq reference in which one scRNA-seq sample was integrated with two replicate samples of snRNA-seq, all from the same study. b) UMAP of txci-ATAC-seq data annotated with the labels predicted by the integrated scRNA-seq reference. c) UMAP of the sci-ATAC-seq reference. d) UMAP of txci-ATAC-seq data annotated with the labels predicted by the sci-ATAC-seq reference. The color legend for all panels is shown on the right. The legend labels with the assay enclosed in parentheses (and connected to a color with a line) denote that these cell-type labels are only observed in one reference (“RNA” for the data shown in (a), and “ATAC” for the data shown in (b)) and share a color with a cell type that is only observed in the other reference. Abbreviations: aBC, activated B cells; aEC, arterial endothelial cells; AM, alveolar macrophages; AT1, alveolar type 1 epithelial cells; AT2, alveolar type 2 epithelial cells; BC, B cells; cEC, capillary endothelial cells; Cil, ciliated cells; cMono, classical monocytes; Col13+FB, collagen type XIII α 1 chain positive fibroblasts; Col14+FB, collagen type XIV α 1 chain positive fibroblasts; DC, dendritic cells; Div, dividing cells; EC, endothelial cells; GB, germinal B cells; Gob, goblet cells; HSPC, Hematopoietic progenitors; IM, interstitial macrophages; LEC, lymphatic endothelial cells; Mes, mesothelial cells; MyoFB, myofibroblasts; ncMono, nonclassical monocytes; NK, natural killer cells; pDC, plasmacytoid dendritic cells; Peri, pericytes; PMN, neutrophils; SMC, smooth muscle cells; TC, T cells; Treg, regulatory T cells; vEC, venous endothelial cells.

**Supplementary Figure 6.**
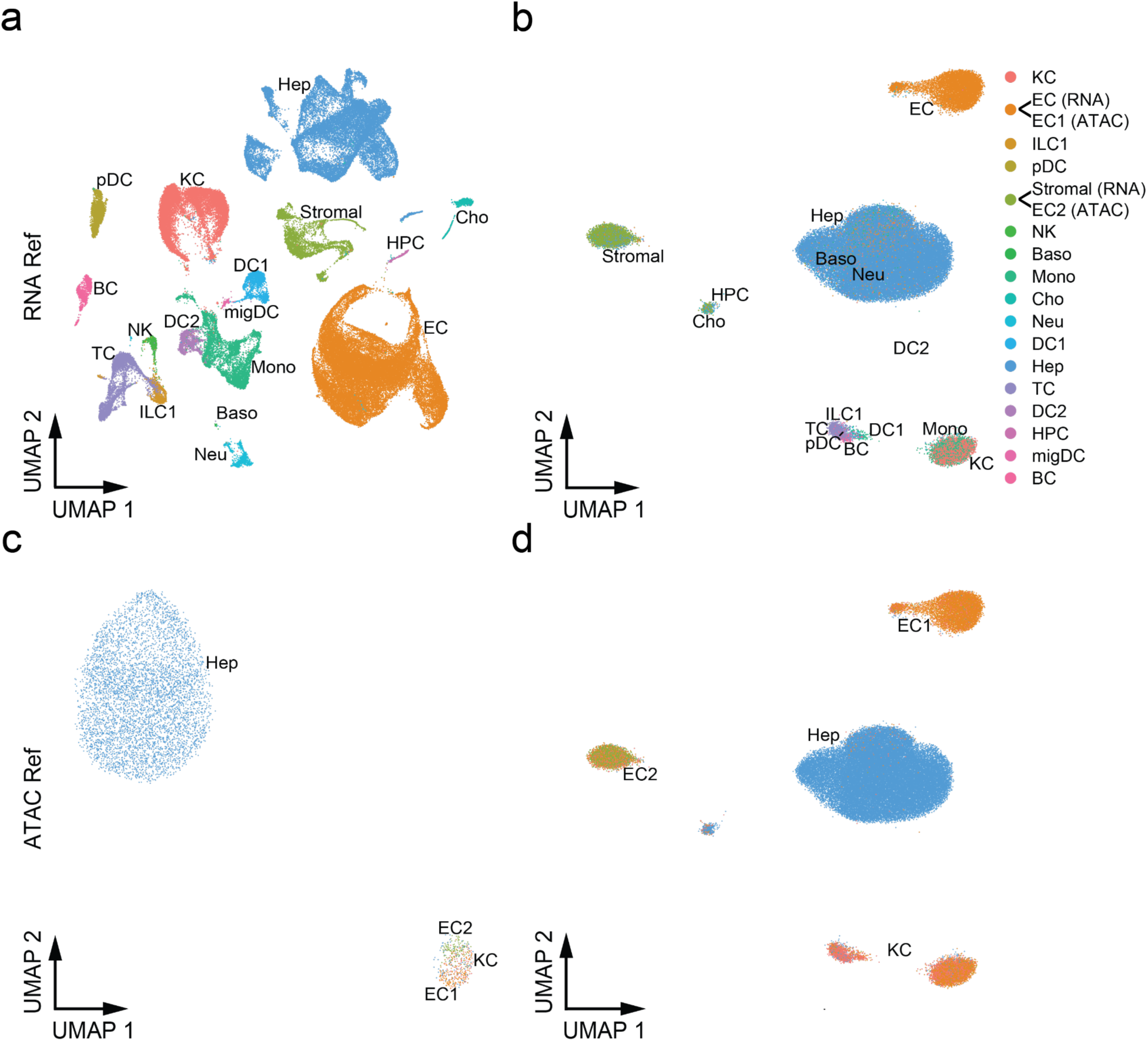
Cell type annotation of mouse liver samples with label transfer. a) UMAP of sc/snRNA-seq reference integrating the snRNA-seq with scRNA-seq using different digestion protocols. b) UMAP of txci-ATAC-seq data annotated with the labels predicted by scRNA-seq reference. c) UMAP of sci-ATAC-seq reference. d) UMAP of txci-ATAC-seq data annotated with the labels predicted by sci-ATAC-seq reference. The color legend for all panels is shown on the right. The legend labels with the assay enclosed in parentheses (and connected to a color with a line) denote that these cell-type labels are only observed in one reference (“RNA” for the data shown in (a), and “ATAC” for the data shown in (b)) and share a color with a cell type that is only observed in the other reference. Abbreviations: Baso, basophils; BC, B cells; Cho, cholangiocytes; DC, conventional dendritic cells; EC, endothelial cells; Hep, hepatocytes; HPC, hepatic progenitor cells; ILC1, type 1 innate lymphoid cells; KC, Kupffer cells; migDC, migratory DCs; Mono, monocytes and monocyte-derived cells; Neu, neutrophils; NK, NK cells; pDC, plasmacytoid dendritic cells; TC, T cells.

**Supplementary Figure 7.**
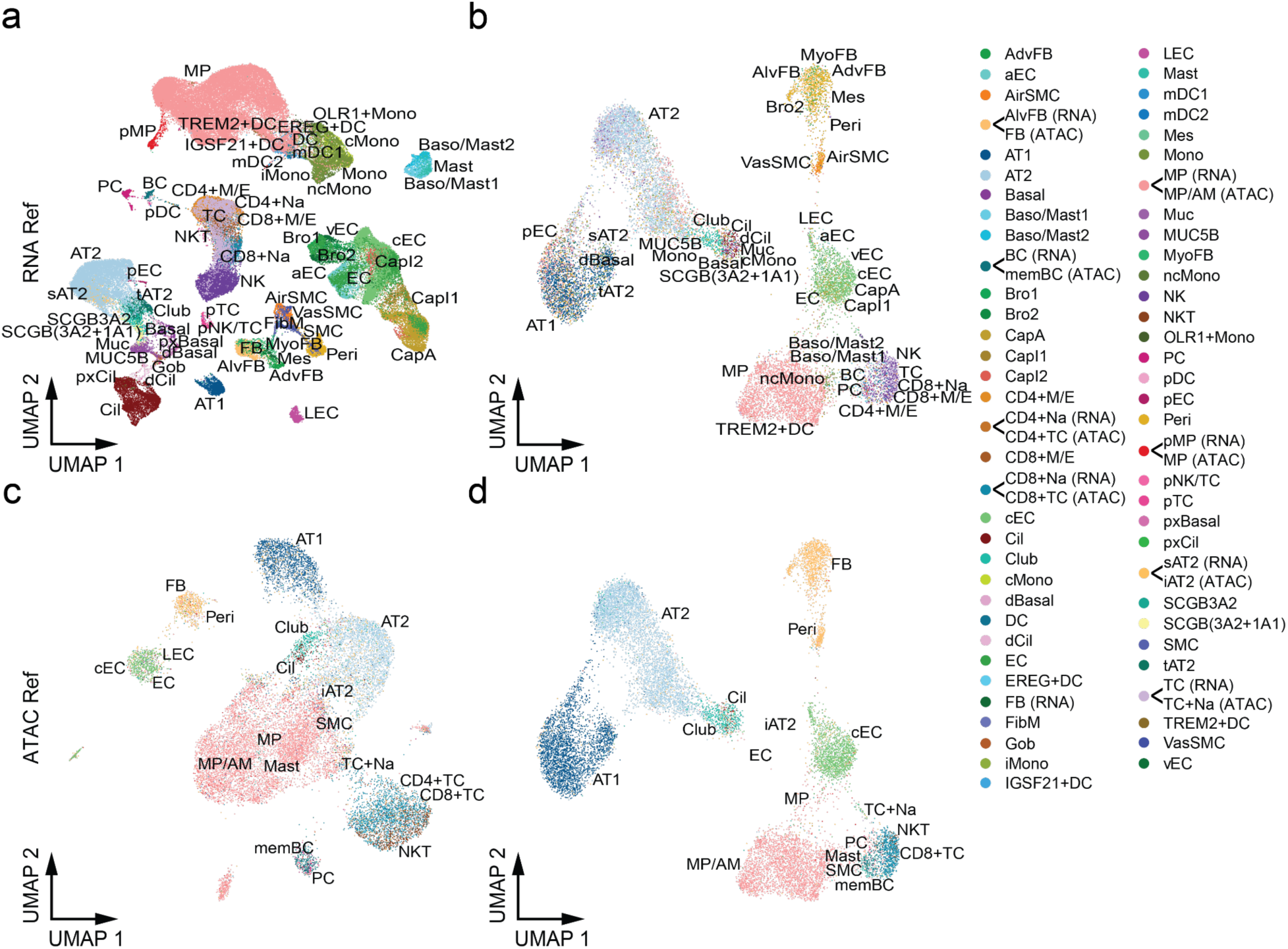
Cell type annotation of human lung sample with label transfer. a) UMAP of scRNA-seq reference integrating the two scRNA-seq datasets. b) UMAP of txci-ATAC-seq data annotated with the labels predicted by scRNA-seq reference. c) UMAP of sci-ATAC-seq reference. d) UMAP of txci-ATAC-seq data annotated with the labels predicted by sci-ATAC-seq reference. The color legend for all panels is shown on the right. The legend labels with the assay enclosed in parentheses (and connected to a color with a line) denote that these cell-type labels share a color with a cell type that is observed in the alternative reference (“RNA” for the data shown in (a), and “ATAC” for the data shown in (b)). Note: the fibroblasts (FB) in ATAC reference share the color with alveolar fibroblasts (AlvFB) rather than FB in RNA reference; The macrophages (MP) in RNA reference share the color with general/alveolar macrophages (MP/AM) rather than MP in ATAC reference. Abbreviations: AdvFB, adventitial fibroblasts; aEC, arterial endothelial cells; AirSMC, airway smooth muscle cells; AlvFB, alveolar fibroblasts; AT1, alveolar type 1 epithelial cells; AT2, alveolar type 2 epithelial cells; Baso/Mast1, basophil/mast cell 1 cells; Baso/Mast2, basophil/mast cell 2 cells; BC, B cells; Bro1, bronchial vessel 1 cells; Bro2, bronchial vessel 2 cells; CapA, capillary aerocytes; CapI1, capillary intermediate 1 cells; CapI2, capillary intermediate 2 cells; CD4+M/E, CD4+ memory/effector T cells; CD4+Na, CD4+ naive T cells; CD4+TC, CD4+ T cells; CD8+M/E, CD8+ memory/effector T cells; CD8+Na, CD8+ naive T cells; CD8+TC, CD8+ T cells; cEC, capillary endothelial cells; Cil, ciliated cells; cMono, classical monocytes; dBasal, differentiating basal cells; DC, conventional dendritic cells; dCil, differentiating ciliated cells; EC, endothelial cells; FB, fibroblasts; FibM, fibromyocytes; Gob, goblet cells; iAT2, alveolar type 2/immune; iMono, intermediate monocytes; LEC, lymphatic endothelial cells; mDC1, myeloid dendritic type 1 cells; mDC2, myeloid dendritic type 2 cells; memBC, memory B cells; Mes, mesothelial cells; Mono, monocytes; MP, macrophages; MP/AM, macrophages (general/alveolar); Muc, mucous cells; MUC5B, MUC5B+ secretory cells; MyoFB, myofibroblasts; ncMono, nonclassical monocytes; NK, natural killer cells; NKT, natural killer T cells; PC, plasma cells; pDC, plasmacytoid dendritic cells; pEC, proliferating epithelial cells; Peri, pericytes; pMP, proliferating macrophages; pNK/TC, proliferating NK/T cells; pTC, proliferating T cells; pxBasal, proximal basal cells; pxCil, proximal ciliated cells; sAT2, signaling AT2 cells; SCGB3A2, SCGB3A2+ secretory cells; SCGB(3A2+1A1), SCGB3A2+ and SCGB1A1+ secretory cells; SMC, smooth muscle cells; tAT2, transitional AT2 cells; TC, T cells; TC+Na, naive T cells; VasSMC, vascular smooth muscle cells; vEC, venous endothelial cells.

**Supplementary Figure 8.**
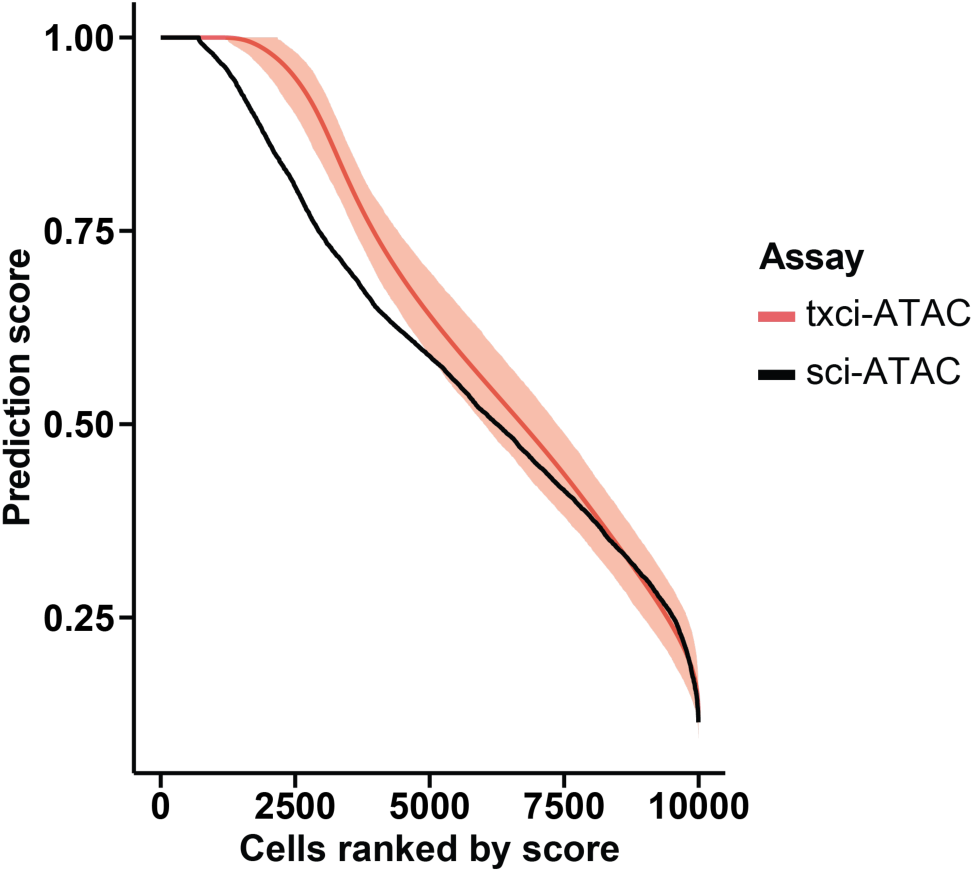
Comparison of prediction accuracy between txci-ATAC-seq and sci-ATAC-seq in mouse lung cells. The txci-ATAC-seq dataset was subsampled to have the same number of cells as that in sci-ATAC-seq data 1000 times. The prediction score (y-axis) calculated by Seurat label transfer using an RNA-seq reference was plotted against the cell ranks (x-axis) based on the prediction score. The red line shows the mean score of 1000 simulations in txci-ATAC-seq. The shaded band is the pointwise 95% confidence interval based on subsampling (from the 2.5% to 97.5% quantile). The black line shows the prediction score in sci-ATAC-seq data.

**Supplementary Figure 9.**
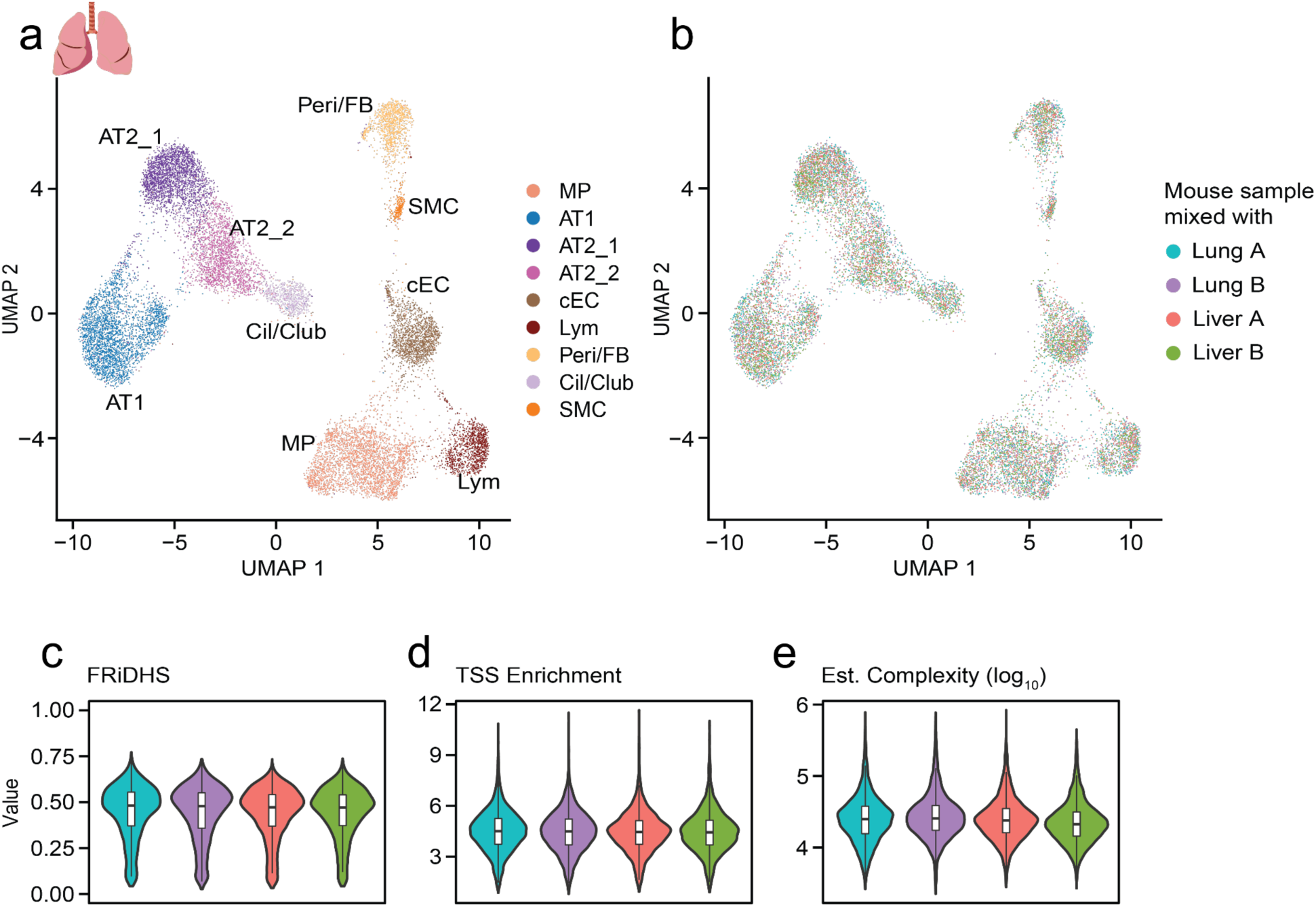
Characterization of cellular heterogeneity in human lung tissue. a) UMAP visualization of human lung nuclei (n = 15,799) identifying 9 distinct cell types. b) UMAP of human lung nuclei visualized by the mouse samples with which they were mixed. c-d) QC metrics of human lung nuclei mixed with different mouse samples. The color legend is consistent with panel (b). The (c) FRiDHS, (d) TSS enrichment score, and (e) estimated complexity (on a log_10_ scale) are plotted for each human lung nuclei sample profiled across different barnyard settings.

**Supplementary Figure 10.**
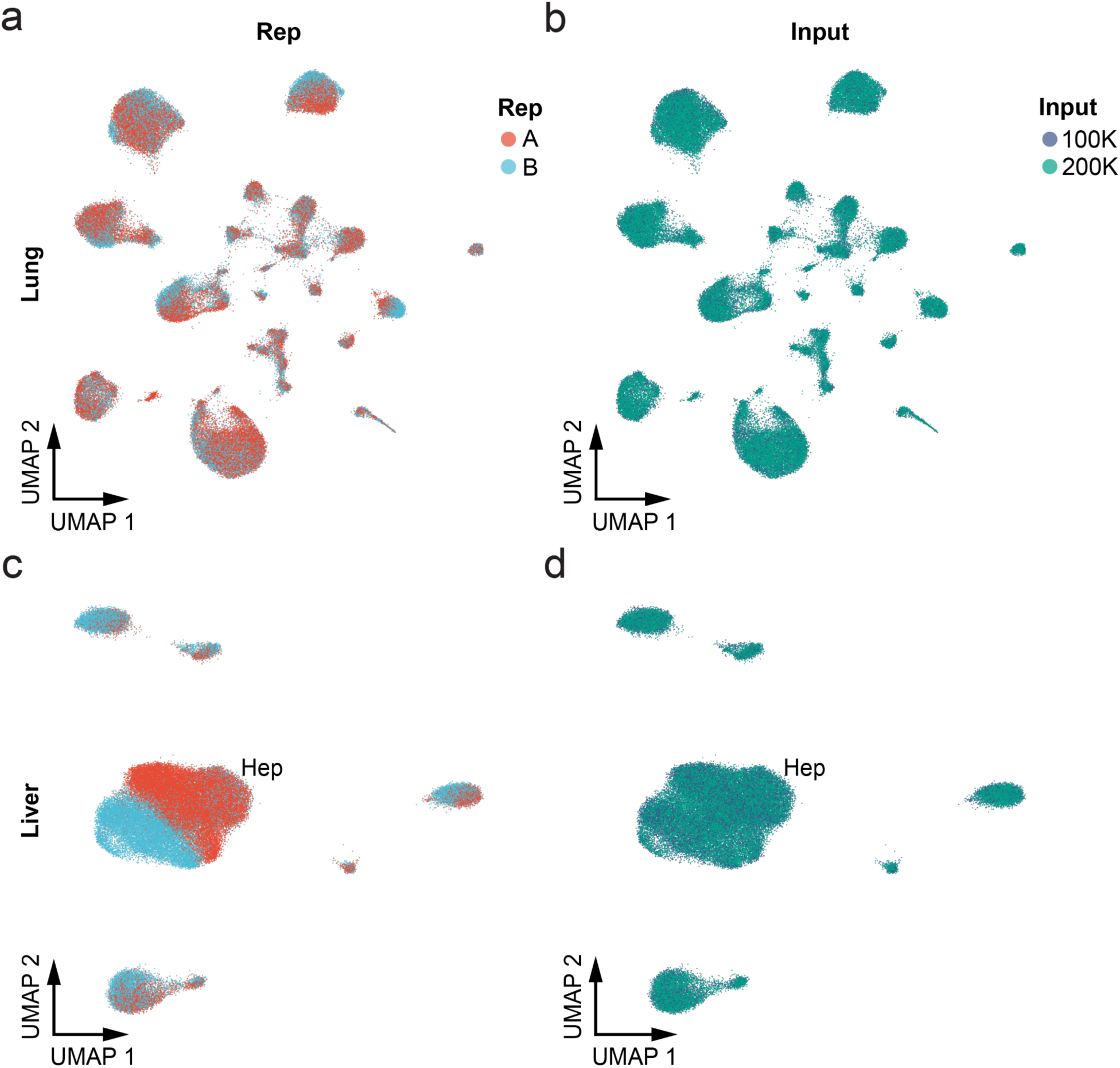
txci-ATAC-seq are robust to batch effects. a,b) UMAP visualization of mouse lung samples showing the batch variance introduced either by (a) mouse replicate or (b) nuclei input on the 10X. c,d) UMAP visualization of mouse liver samples showing the batch variance introduced either by (c) mouse replicate or (d) nuclei input. Colors for replicates and inputs are consistent in both (a,c) and (b,d). Hepatocytes are indicated by “Hep” in (c).

**Supplementary Figure 11.**
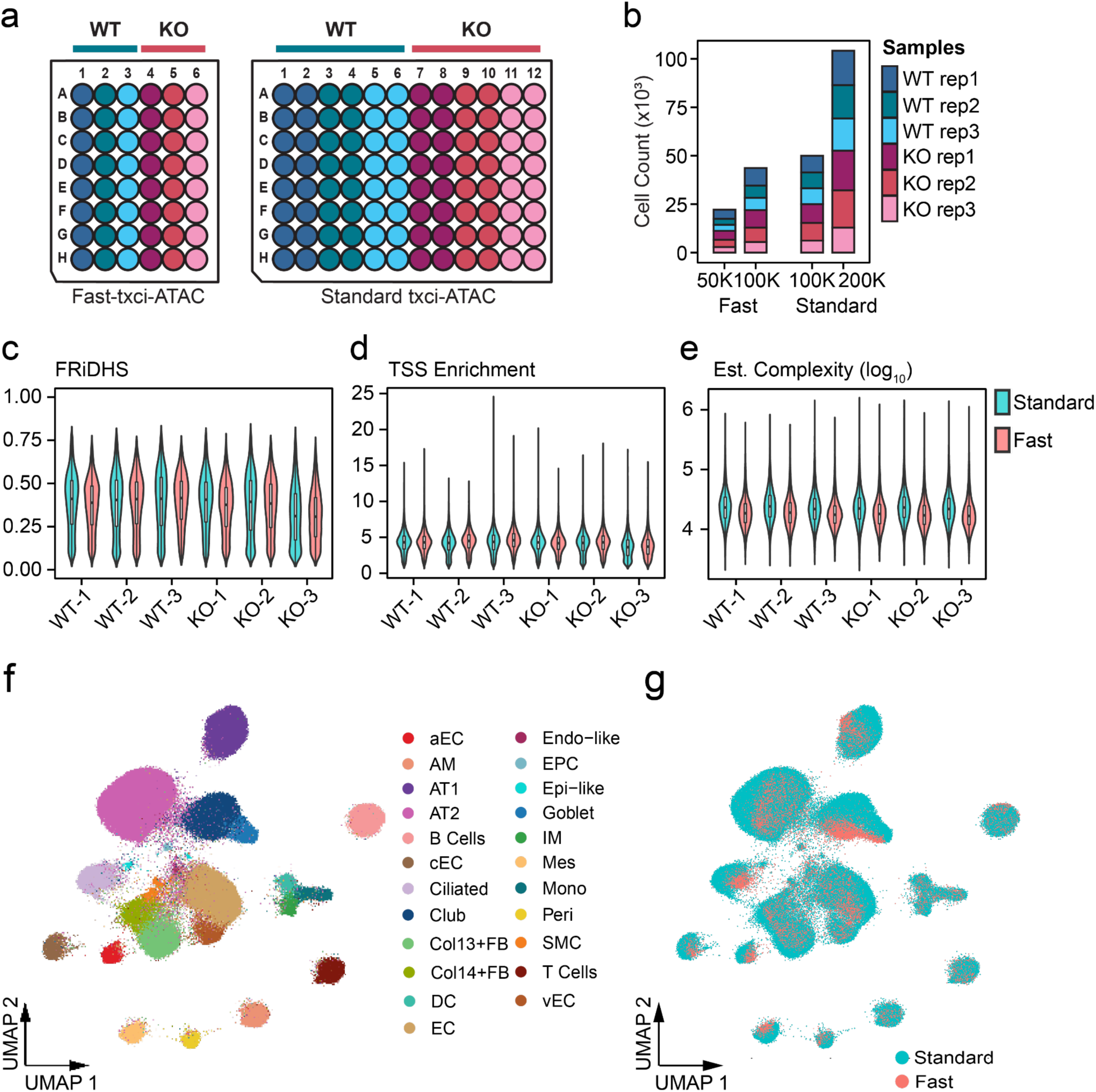
Fast-txci-ATAC-seq improves multiplexing capability without sacrificing data quality. a) Well assignment for the faster and standard versions of txci-ATAC-seq multiplexing WT and CC16 knockout lungs with (3 replicate mice for each genotype). b) The number of nuclei passing quality filters for each protocol at different nuclei loading inputs. c-e) The comparison of quality metrics per cell between the two protocols across 6 mouse lung samples. The Fast-txci-ATAC-seq provided a comparable FRiDHS (c) and TSS enrichment score (d) but slightly lower estimated complexity (e) than the standard protocol. f, g) UMAP visualization of nuclei by co-embedding the standard (n=154,103) and faster (n=65,799) assays. The nuclei are colored either by predicted cell type (f) or ATAC-seq protocol (g).

**Supplementary Figure 12.**
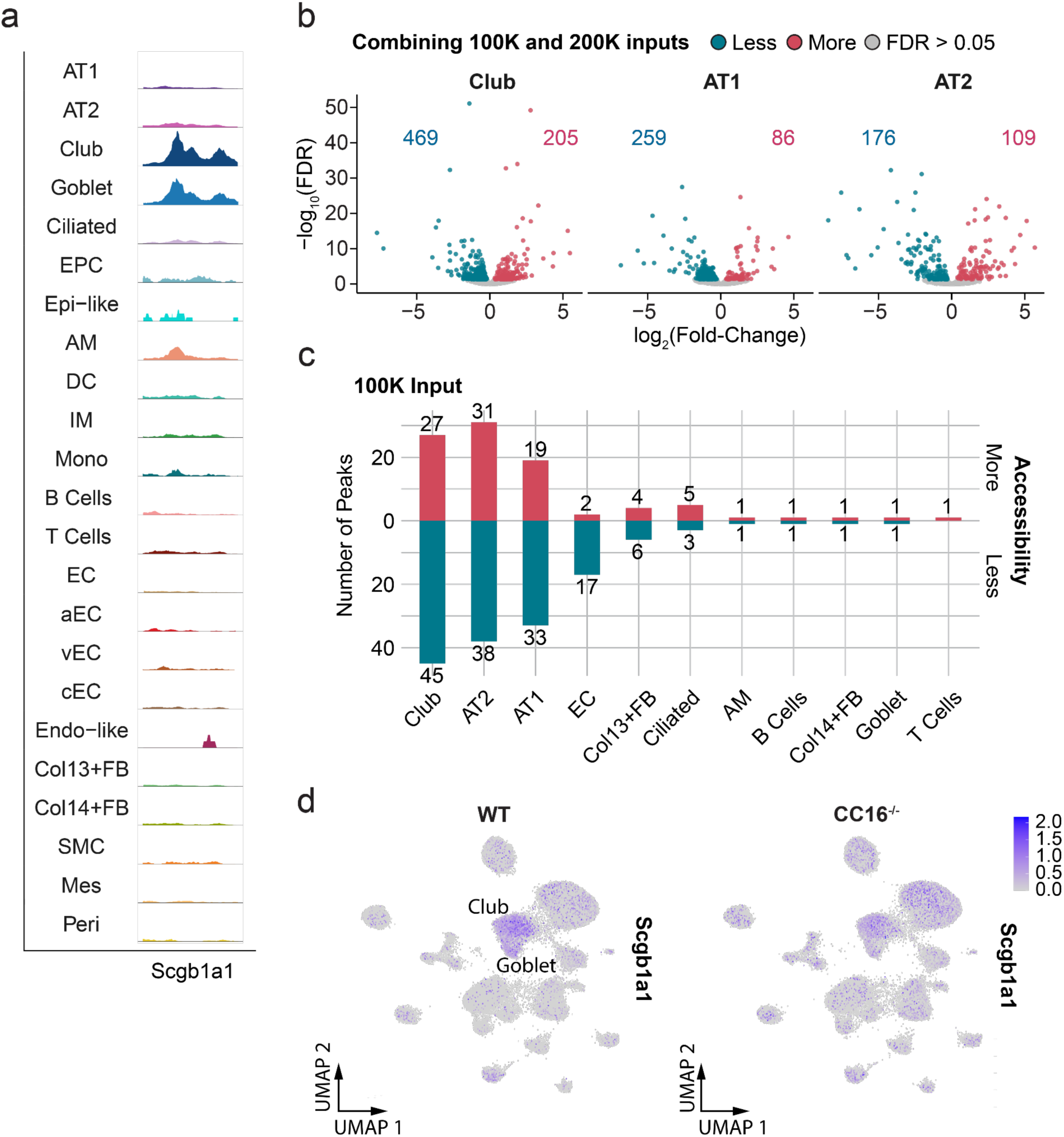
Chromatin accessibility changes induced by CC16^-/-^ deficiency in mouse lung. a) Aggregated chromatin accessibility at the *Scgb1a1* promoter region across cell types. The aggregated accessibility signal for each cluster was normalized by a scaling factor computed as the number of cells in the cluster multiplied by the mean sequencing depth for the cells in that cluster. b) Differentially accessible peaks between CC16^-/-^ and WT samples across club, AT1, and AT2 cells generated by combining the 100,000 and 200,000 nuclei loading inputs. The -log_10_-transformed adjusted p-value for each peak was plotted against the log_2_(fold-change). The color labels the peaks that are less accessible (blue), more accessible (red), and unchanged (gray) in knockout samples. The number of differentially accessible peaks identified in each cell type is displayed within the plot. c) The number of differentially accessible peaks per cell type calculated using the 100,000 input alone. With 100,000 nuclei as input, many fewer peaks were identified as differentially accessible. d) UMAP visualization of nuclei showing the gene activity score of *Scgb1a1* in WT (left) and CC16^-/-^ (right) lungs.

**Supplementary Figure 13.**
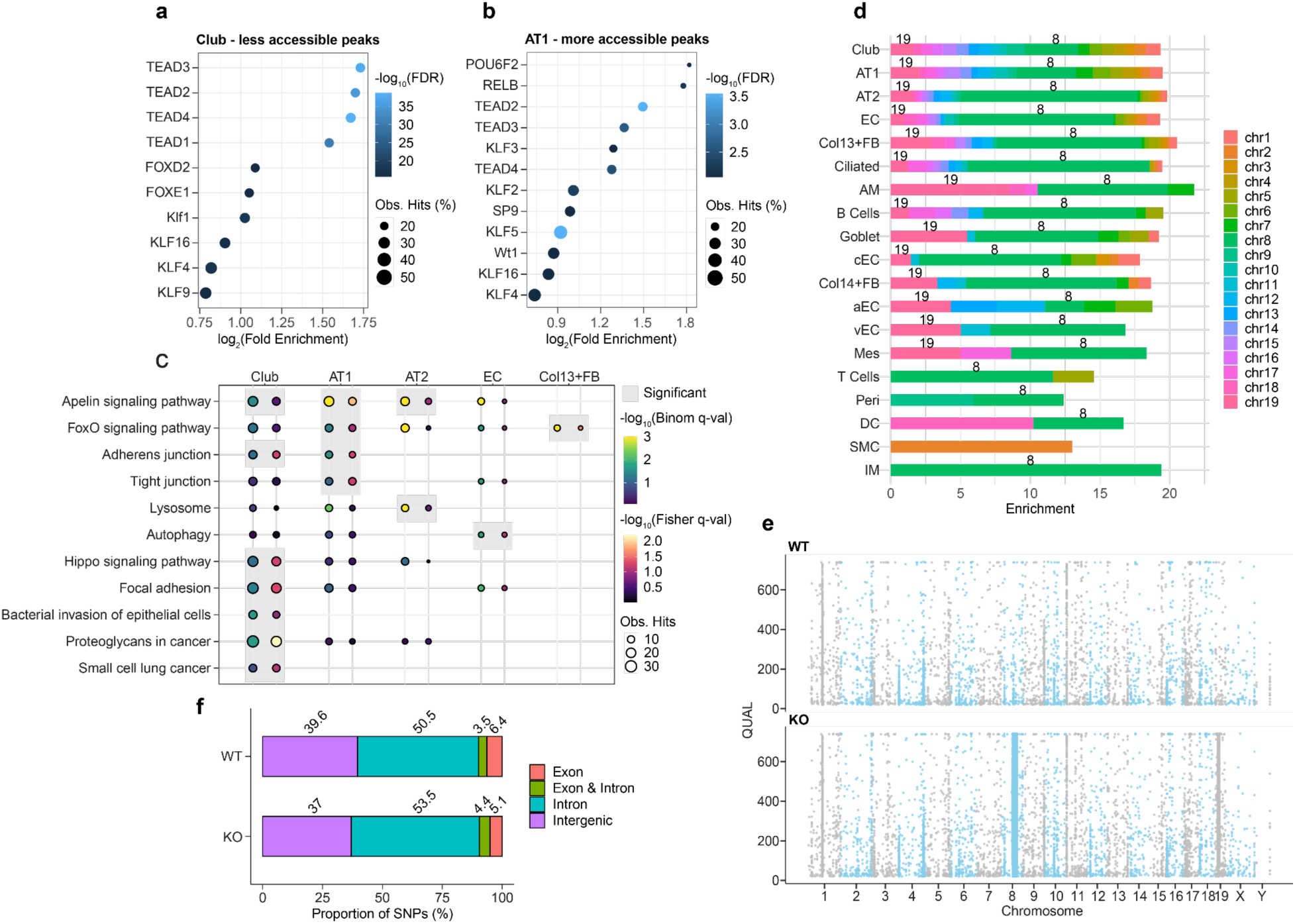
Functional analysis and regulatory variant identification in CC16 deficient mouse. a,b) Top 10 motifs significantly enriched in less accessible peaks identified in CC16^-/-^ club cells (a) and more accessible peaks identified in CC16^-/-^ AT1 cells (b). The dot color encodes the -log_10_-transformed adjusted p-value, while the size of the dot encodes the percentage of observed peaks enriched in each motif. The x-axis displays the fold enrichment on a log_2_ scale. c) KEGG pathways of interest enriched in each cell type. The colors show the -log_10_-transformed adjusted p-value derived from the binomial (blue to green) and hypergeometric (black to red) tests. The dot size denotes the number of observed regions (binomial) or genes (hypergeometric) in each test. The significant pathways passing the two-threshold cutoff are highlighted by gray boxes. d) Enrichment of differential peaks in each chromosome across cell types. The enrichment was calculated by dividing the fraction of differential peaks in each chromosome by the fraction of total peaks identified in each chromosome. e) SNVs identified in WT and knockout samples across chromosomes meeting the quality criteria. The y-axis represents the phred-scaled quality score. f) Proportions of SNPs mapped to each functional genomic category (exon, intron, overlapping regions between exon and intron, and intergenic regions) in WT and knockout samples.

**Supplementary Figure 14.**
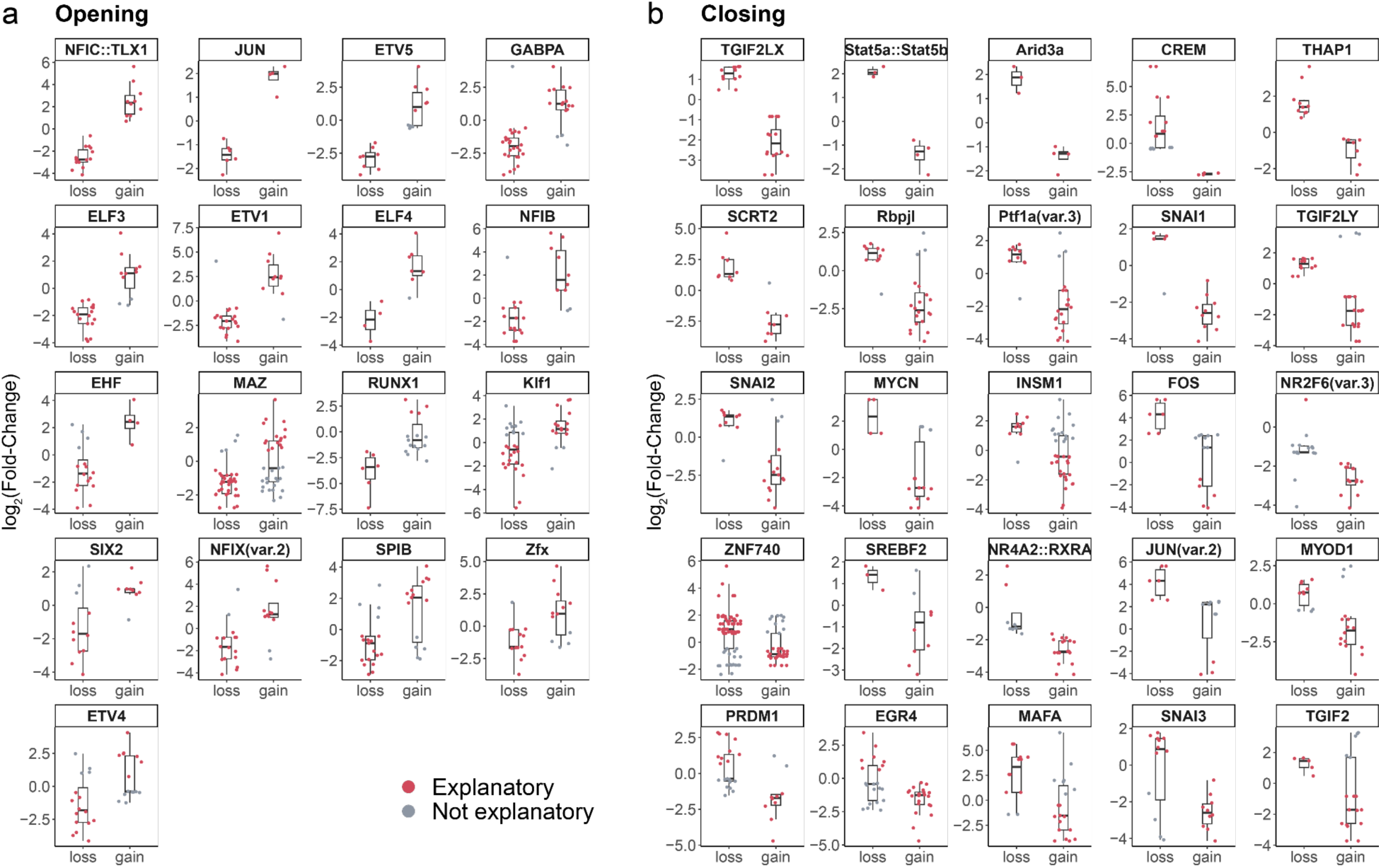
SNP-driven differences in motif usage alter chromatin accessibility. a,b) TFs that are associated with increased (a) or decreased (b) chromatin accessibility when the peaks gain the TF motifs. The y-axis shows the log_2_(fold-change) in chromatin accessibility for the differential peaks identified in the SNV hotspots between CC16^-/-^ and WT samples. A positive log_2_(fold-change) means the peaks are more accessible in the knockout samples. The x-axis indicates the motif hits that are gained or lost in the peaks carrying the CC16^-/-^ SNVs. The instances (red) that exhibit a coherent change in chromatin accessibility with the overall motif effect were considered to explain the observed differences in chromatin accessibility between two genotypes (i.e., for opening TF motifs shown in panel a, the gained instances with a positive log_2_(fold-change) and the lost instances with a negative log_2_(fold-change) were considered explanatory; For closing TF motifs shown in panel b, the gained instances with a negative log_2_(fold-change) and the lost instances with a positive log_2_(fold-change) were considered explanatory).

**Supplementary Figure 15.**
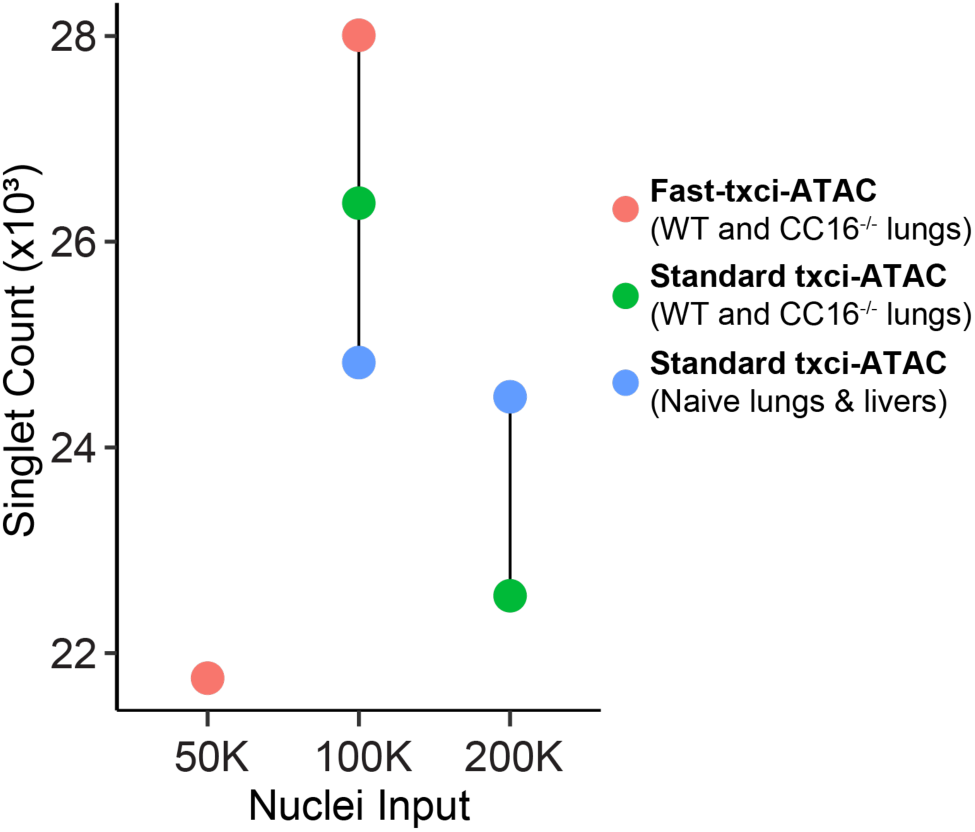
Singlet yields at different numbers of input nuclei. Plot showing the number of nuclei recovered (y-axis) in each experiment stratified by the intended number of input nuclei (x-axis). The color denotes the datasets used to compute the singlets at each nuclei loading input. The maximum yield was obtained with 100,000 nuclei as input.

**Supplementary Figure 16.**
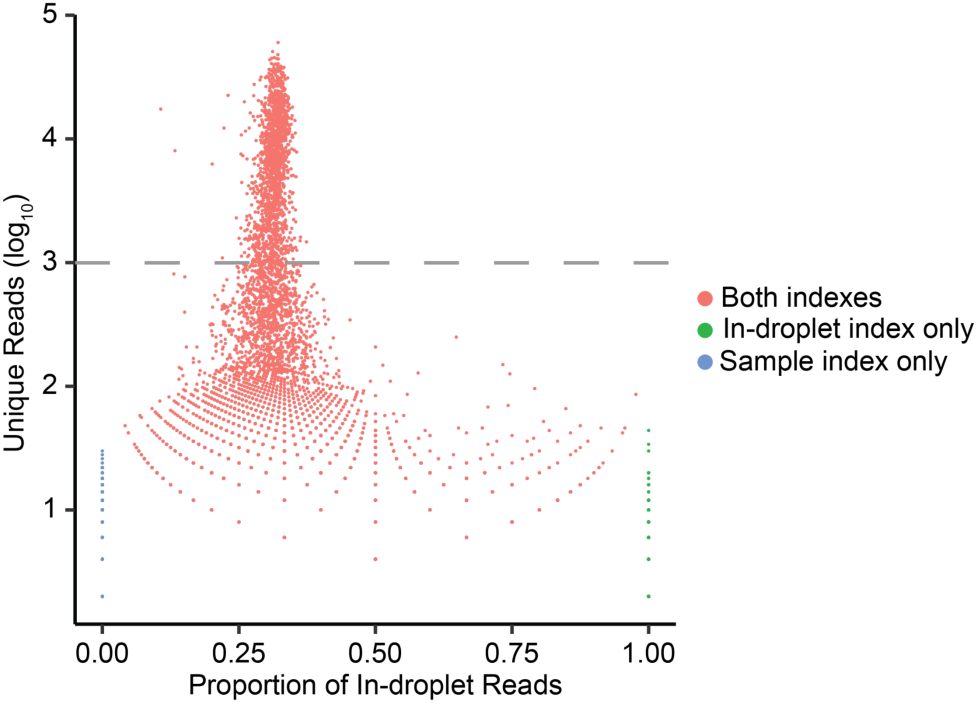
Examination of efficiency for in-droplet and sample index PCR. The total unique reads on a log_10_ scale were plotted against the proportion of in-droplet reads for each barcode. The barcodes with both in-droplet and sample indexes were colored in red. The green and blue dots indicate the barcodes only getting the in-droplet index and the barcodes only getting the sample index, respectively. The gray dashed line indicates the threshold to call a cell barcode.

## Supplementary Tables

**Supplementary Table 1.** Differentially accessible peaks identified between CC16^-/-^ and WT samples for each cell type. This table is provided as a separate file. Column 1: Cell type in which the test was performed. Column 2: Peak region. Columns 3-5: Log_2_(fold-change), raw *p*-value, and FDR-adjusted *p*-value, respectively. Columns 6-11: Log_2_-transformed counts per million (CPM) for each sample, computed using the normalized library sizes. The CPM values for WT samples are shown in columns 6-8, and the CPM values for CC16^-/-^ samples are shown in columns 9-11.

**Supplementary Table 2.** Enriched motifs in more accessible and less accessible peaks in response to CC16 deficiency for each cell type. This table is provided as a separate file. Column 1: Cell type in which the test was performed. Column 2: Changing direction of differentially accessible peaks that were used to perform the test. Column 3: Motif ID. Column 4: Motif name. Column 5: The number of differential peaks that contain the motif identified. Column 6: The number of background peaks that contain the motif identified. Column 7: The percentage of differential peaks that contain the motif identified. Column 8: The percentage of background peaks that contain the motif identified. Column 9: The ratio of the observed frequency of the motif in differential peaks to the expected frequency calculated by the background peaks. Column 10: Raw *p*-value. Column 11: FDR-adjusted *p*-value.

**Supplementary Table 3.** KEGG pathways enriched in differential peaks between CC16^-/-^ and WT samples for each cell type. This table is provided as a separate file. Column 1: Cell type in which the test was performed. Column 2: KEGG pathway ID. Column 3: Description of KEGG pathway. Column 4: Fraction of non-gap base pairs in the genome that lie in the regulatory domain of a gene with the annotation. Column 5: Actual number of differential peaks with the annotation. Column 6: Fold enrichment of number of differential peaks with the annotation. Column 7: Uncorrected *p*-value from the binomial test over genomic regions. Column 8: FDR-adjusted *p*-value for the binomial test. Column 9: Mean absolute distance of input regions to TSS of genes in a gene set. Column 10: Actual number of genes linking to a differential peak with the annotation. Column 11: Number of genes in the genome with the annotation. Column 12: Fold enrichment of number of genes linking to a differential peak with the annotation. Column 13: Uncorrected *p*-value from the hypergeometric test over genes. Column 14: FDR-adjusted *p*-value for the hypergeometric test.

**Supplementary Table 4.**
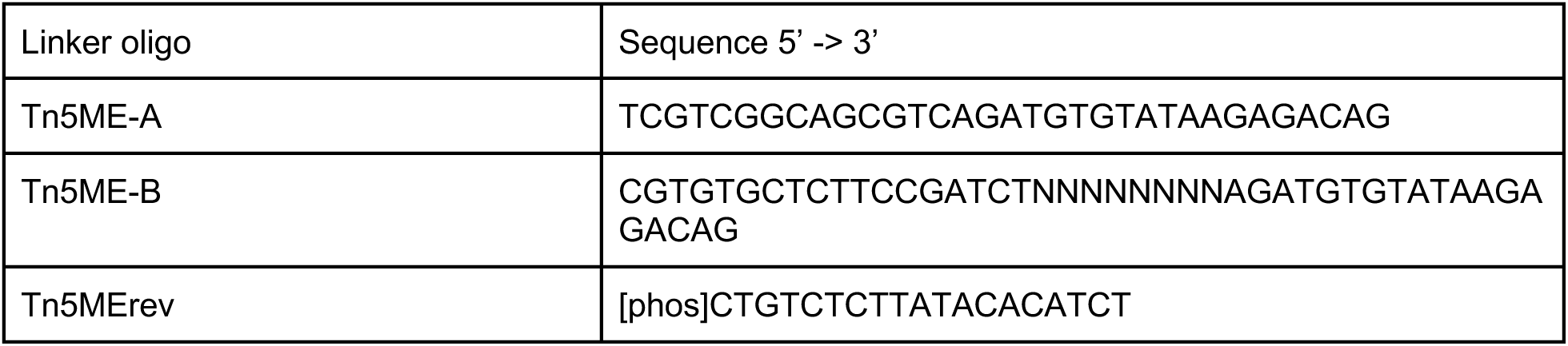
Sequences of Tn5 linker oligos. The ‘N’ bases shown in the Tn5ME-B sequence represent the Tn5 barcodes.

**Supplementary Table 5.** Tn5 barcode sequences. Column 1 shows the well ID for each well on the iTSM plate. Column 2 shows the sequences of Tn5 barcodes assigned to each well. Column 3 is the 12 numerical labels for the plate columns. Column 4 is the 8 alphabetical labels for the plate rows. This table is provided as a separate file.

**Supplementary Table 6.** Barnyard experiment design. Column 1 shows the figure number for each barnyard experiment. Column 2 indicates the barnyard type (True vs. Pseudo). Column 3 shows the cell source of human samples. Column 4 shows the number of human nuclei loaded to each well. Column 5 shows the cell source of mouse samples. Column 6 shows the number of mouse nuclei loaded to each well. Column 7 indicates the nuclei preparation method (Fresh vs Frozen). Column 8 is the well ID on the iTSM plate (see Table S4) assigned to each barnyard experiment. This table is provided as a separate file.

**Supplementary Table 7.**
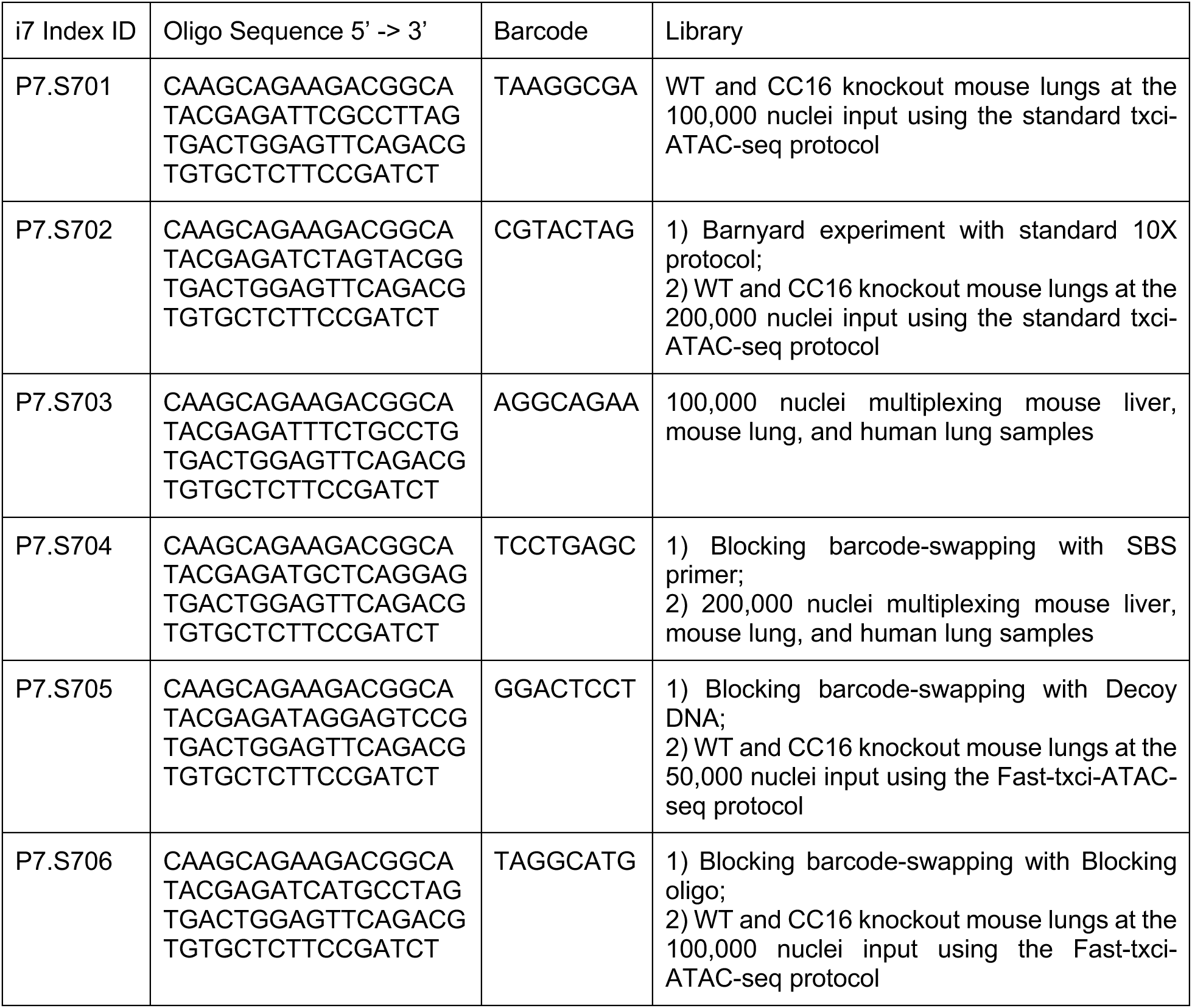
TruSeq i7 index sequences used for each library in Sample Index PCR. Column 1 shows the index ID. Column 2 shows the oligo sequence. Column 3 indicates the barcode sequence assigned to each library shown in Column 4.

**Supplementary Table 8.**
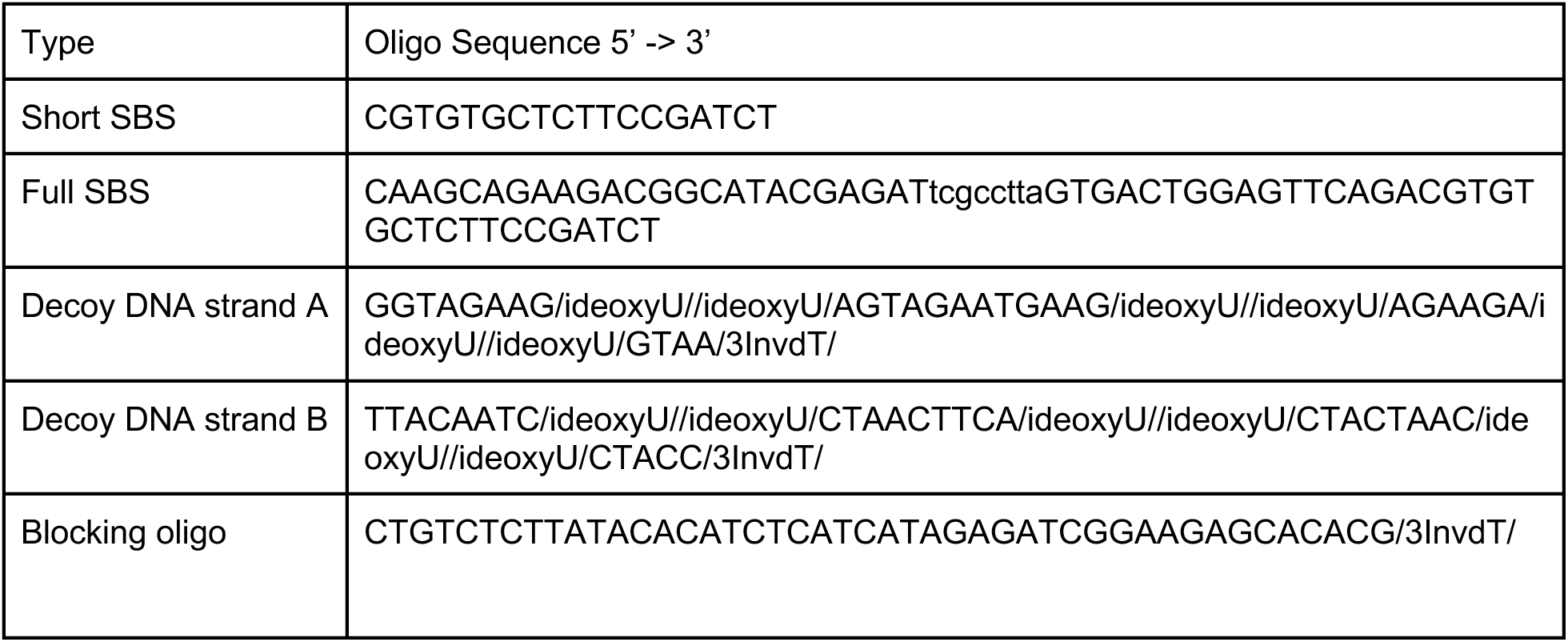
DNA oligonucleotides used to block barcode swapping. Each row provides the sequence of an oligo used in the barcode swapping blocking tests. The lowercase letters shown in the full SBS primer represent the barcode sequence. For Decoy DNA, the strands A and B were annealed to form a duplex DNA.

## References

1. Buenrostro JD, Giresi PG, Zaba LC, Chang HY, Greenleaf WJ. Transposition of native chromatin for fast and sensitive epigenomic profiling of open chromatin, DNA-binding proteins and nucleosome position. Nat Methods. 2013;10: 1213–1218.

2. Domcke S, Hill AJ, Daza RM, Cao J, O’Day DR, Pliner HA, et al. A human cell atlas of fetal chromatin accessibility. Science. 2020;370. doi:10.1126/science.aba7612

3. Cusanovich DA, Hill AJ, Aghamirzaie D, Daza RM, Pliner HA, Berletch JB, et al. A Single-Cell Atlas of In Vivo Mammalian Chromatin Accessibility. Cell. 2018;174: 1309–1324.e18.

4. Satpathy AT, Granja JM, Yost KE, Qi Y, Meschi F, McDermott GP, et al. Massively parallel single-cell chromatin landscapes of human immune cell development and intratumoral T cell exhaustion. Nat Biotechnol. 2019;37: 925–936.

5. Cusanovich DA, Daza R, Adey A, Pliner HA, Christiansen L, Gunderson KL, et al. Multiplex single cell profiling of chromatin accessibility by combinatorial cellular indexing. Science. 2015;348: 910–914.

6. Cao J, Packer JS, Ramani V, Cusanovich DA, Huynh C, Daza R, et al. Comprehensive single-cell transcriptional profiling of a multicellular organism. Science. 2017;357: 661–667.

7. Vitak SA, Torkenczy KA, Rosenkrantz JL, Fields AJ, Christiansen L, Wong MH, et al. Sequencing thousands of single-cell genomes with combinatorial indexing. Nat Methods. 2017;14: 302–308.

8. Wang K, Xiao Z, Yan Y, Ye R, Hu M, Bai S, et al. Simple oligonucleotide-based multiplexing of single-cell chromatin accessibility. Mol Cell. 2021;81: 4319–4332.e10.

9. Lareau CA, Duarte FM, Chew JG, Kartha VK, Burkett ZD, Kohlway AS, et al. Droplet-based combinatorial indexing for massive-scale single-cell chromatin accessibility. Nat Biotechnol. 2019;37: 916–924.

10. Corces MR, Trevino AE, Hamilton EG, Greenside PG, Sinnott-Armstrong NA, Vesuna S, et al. An improved ATAC-seq protocol reduces background and enables interrogation of frozen tissues. Nat Methods. 2017;14: 959–962.

11. Bloom JD. Estimating the frequency of multiplets in single-cell RNA sequencing from cell-mixing experiments. PeerJ. 2018;6: e5578.

12. Bravo González-Blas C, Minnoye L, Papasokrati D, Aibar S, Hulselmans G, Christiaens V, et al. cisTopic: cis-regulatory topic modeling on single-cell ATAC-seq data. Nat Methods. 2019;16: 397–400.

13. Butler A, Hoffman P, Smibert P, Papalexi E, Satija R. Integrating single-cell transcriptomic data across different conditions, technologies, and species. Nat Biotechnol. 2018;36: 411– 420.

14. Wolock SL, Lopez R, Klein AM. Scrublet: Computational Identification of Cell Doublets in Single-Cell Transcriptomic Data. Cell Syst. 2019;8: 281–291.e9.

15. Pliner HA, Packer JS, McFaline-Figueroa JL, Cusanovich DA, Daza RM, Aghamirzaie D, et al. Cicero Predicts cis-Regulatory DNA Interactions from Single-Cell Chromatin Accessibility Data. Mol Cell. 2018;71: 858–871.e8.

16. Stuart T, Srivastava A, Madad S, Lareau CA, Satija R. Single-cell chromatin state analysis with Signac. Nature Methods. 2021. doi:10.1038/s41592-021-01282-5

17. Hodge RD, Bakken TE, Miller JA, Smith KA, Barkan ER, Graybuck LT, et al. Conserved cell types with divergent features in human versus mouse cortex. Nature. 2019;573: 61–68.

18. Lein ES, Hawrylycz MJ, Ao N, Ayres M, Bensinger A, Bernard A, et al. Genome-wide atlas of gene expression in the adult mouse brain. Nature. 2007;445: 168–176.

19. Allen Institute for Brain Science. Allen Brain Cell Types Database. (2019).

20. Allen Institute for Brain Science. Allen Human Brain Atlas. (2020).

21. Korsunsky I, Nathan A, Millard N, Raychaudhuri S. Presto scales Wilcoxon and auROC analyses to millions of observations. doi:10.1101/653253

22. Schep AN, Wu B, Buenrostro JD, Greenleaf WJ. chromVAR: inferring transcription-factor-associated accessibility from single-cell epigenomic data. Nat Methods. 2017;14: 975–978.

23. Preissl S, Fang R, Huang H, Zhao Y, Raviram R, Gorkin DU, et al. Single-nucleus analysis of accessible chromatin in developing mouse forebrain reveals cell-type-specific transcriptional regulation. Nat Neurosci. 2018;21: 432–439.

24. Thornton CA, Mulqueen RM, Torkenczy KA, Nishida A, Lowenstein EG, Fields AJ, et al. Spatially mapped single-cell chromatin accessibility. Nat Commun. 2021;12: 1274.

25. Mulqueen RM, Pokholok D, O’Connell BL, Thornton CA, Zhang F, O’Roak BJ, et al. High-content single-cell combinatorial indexing. Nat Biotechnol. 2021;39: 1574–1580.

26. 10X scATAC-seq v1 dataset for fresh cortex from adult mouse brain. Available: https://www.10xgenomics.com/resources/datasets/fresh-cortex-from-adult-mouse-brain-p-50-1-standard-1-2-0

27. 10X scATAC-seq v2 dataset for 8k adult mouse cortex cells. Available: https://www.10xgenomics.com/resources/datasets/8k-adult-mouse-cortex-cells-atac-v2-chromium-x-2-standard

28. Hao Y, Hao S, Andersen-Nissen E, Mauck WM 3rd, Zheng S, Butler A, et al. Integrated analysis of multimodal single-cell data. Cell. 2021;184: 3573–3587.e29.

29. Koenitzer JR, Wu H, Atkinson JJ, Brody SL, Humphreys BD. Single-Nucleus RNA-Sequencing Profiling of Mouse Lung. Reduced Dissociation Bias and Improved Rare Cell-Type Detection Compared with Single-Cell RNA Sequencing. Am J Respir Cell Mol Biol. 2020;63: 739–747.

30. Guilliams M, Bonnardel J, Haest B, Vanderborght B, Wagner C, Remmerie A, et al. Spatial proteogenomics reveals distinct and evolutionarily conserved hepatic macrophage niches. Cell. 2022;185: 379–396.e38.

31. Travaglini KJ, Nabhan AN, Penland L, Sinha R, Gillich A, Sit RV, et al. A molecular cell atlas of the human lung from single-cell RNA sequencing. Nature. 2020;587: 619–625.

32. Habermann AC, Gutierrez AJ, Bui LT, Yahn SL, Winters NI, Calvi CL, et al. Single-cell RNA sequencing reveals profibrotic roles of distinct epithelial and mesenchymal lineages in pulmonary fibrosis. Sci Adv. 2020;6: eaba1972.

33. Zhang K, Hocker JD, Miller M, Hou X, Chiou J, Poirion OB, et al. A cell atlas of chromatin accessibility across 25 adult human tissues. BioRxiv. 2021. doi:10.1101/2021.02.17.431699

34. Mango GW, Johnston CJ, Reynolds SD, Finkelstein JN, Plopper CG, Stripp BR. Clara cell secretory protein deficiency increases oxidant stress response in conducting airways. Am J Physiol. 1998;275: L348–56.

35. Li X, Guerra S, Ledford JG, Kraft M, Li H, Hastie AT, et al. Low CC16 mRNA Expression Levels in Bronchial Epithelial Cells Are Associated with Asthma Severity. Am J Respir Crit Care Med. 2023;207: 438–451.

36. Laucho-Contreras ME, Polverino F, Gupta K, Taylor KL, Kelly E, Pinto-Plata V, et al. Protective role for club cell secretory protein-16 (CC16) in the development of COPD. Eur Respir J. 2015;45: 1544–1556.

37. Guerra S, Vasquez MM, Spangenberg A, Halonen M, Martinez FD. Serum concentrations of club cell secretory protein (Clara) and cancer mortality in adults: a population-based, prospective cohort study. Lancet Respir Med. 2013;1: 779–785.

38. Keane TM, Goodstadt L, Danecek P, White MA, Wong K, Yalcin B, et al. Mouse genomic variation and its effect on phenotypes and gene regulation. Nature. 2011;477: 289–294.

39. Eisener-Dorman AF, Lawrence DA, Bolivar VJ. Cautionary insights on knockout mouse studies: the gene or not the gene? Brain Behav Immun. 2009;23: 318–324.

40. Webb CF, Bryant J, Popowski M, Allred L, Kim D, Harriss J, et al. The ARID family transcription factor bright is required for both hematopoietic stem cell and B lineage development. Mol Cell Biol. 2011;31: 1041–1053.

41. Stoeckius M, Zheng S, Houck-Loomis B, Hao S, Yeung BZ, Mauck WM 3rd, et al. Cell Hashing with barcoded antibodies enables multiplexing and doublet detection for single cell genomics. Genome Biol. 2018;19: 224.

42. Gaublomme JT, Li B, McCabe C, Knecht A, Yang Y, Drokhlyansky E, et al. Nuclei multiplexing with barcoded antibodies for single-nucleus genomics. Nat Commun. 2019;10: 2907.

43. McGinnis CS, Patterson DM, Winkler J, Conrad DN, Hein MY, Srivastava V, et al. MULTI-seq: sample multiplexing for single-cell RNA sequencing using lipid-tagged indices. Nat Methods. 2019;16: 619–626.

44. Kang HM, Subramaniam M, Targ S, Nguyen M, Maliskova L, McCarthy E, et al. Multiplexed droplet single-cell RNA-sequencing using natural genetic variation. Nat Biotechnol. 2018;36: 89–94.

45. Datlinger P, Rendeiro AF, Boenke T, Senekowitsch M, Krausgruber T, Barreca D, et al. Ultra-high-throughput single-cell RNA sequencing and perturbation screening with combinatorial fluidic indexing. Nat Methods. 2021;18: 635–642.

46. Xu W, Yang W, Zhang Y, Chen Y, Hong N, Zhang Q, et al. ISSAAC-seq enables sensitive and flexible multimodal profiling of chromatin accessibility and gene expression in single cells. Nat Methods. 2022;19: 1243–1249.

47. Mulqueen RM, Pokholok D, Norberg SJ, Torkenczy KA, Fields AJ, Sun D, et al. Highly scalable generation of DNA methylation profiles in single cells. Nat Biotechnol. 2018;36: 428–431.

48. Liscovitch-Brauer N, Montalbano A, Deng J, Méndez-Mancilla A, Wessels H-H, Moss NG, et al. Profiling the genetic determinants of chromatin accessibility with scalable single-cell CRISPR screens. Nat Biotechnol. 2021;39: 1270–1277.

49. Cao J, Cusanovich DA, Ramani V, Aghamirzaie D, Pliner HA, Hill AJ, et al. Joint profiling of chromatin accessibility and gene expression in thousands of single cells. Science. 2018;361: 1380–1385.

50. Zhang H, Rice ME, Alvin JW, Farrera-Gaffney D, Galligan JJ, Johnson MDL, et al. Extensive evaluation of ATAC-seq protocols for native or formaldehyde-fixed nuclei. BMC Genomics. 2022;23: 214.

51. Iannuzo N, Insel M, Marshall C, Pederson WP, Addison KJ, Polverino F, et al. CC16 Deficiency in the Context of Early-Life Infection Results in Augmented Airway Responses in Adult Mice. Infect Immun. 2022;90: e0054821.

52. Rojas-Quintero J, Laucho-Contreras ME, Wang X, Fucci Q-A, Burkett PR, Kim S-J, et al. CC16 augmentation reduces exaggerated COPD-like disease in Cc16-deficient mice. JCI Insight. 2023;8. doi:10.1172/jci.insight.130771

53. Lareau CA, Ma S, Duarte FM, Buenrostro JD. Inference and effects of barcode multiplets in droplet-based single-cell assays. Nat Commun. 2020;11: 866.

54. Stripp BR, Lund J, Mango GW, Doyen KC, Johnston C, Hultenby K, et al. Clara cell secretory protein: a determinant of PCB bioaccumulation in mammals. Am J Physiol. 1996;271: L656–64.

55. Zhai J, Insel M, Addison KJ, Stern DA, Pederson W, Dy A, et al. Club Cell Secretory Protein Deficiency Leads to Altered Lung Function. Am J Respir Crit Care Med. 2019;199: 302–312.

56. Saunders A, Core LJ, Sutcliffe C, Lis JT, Ashe HL. Extensive polymerase pausing during Drosophila axis patterning enables high-level and pliable transcription. Genes Dev. 2013;27: 1146–1158.

57. Cusanovich DA, Reddington JP, Garfield DA, Daza RM, Aghamirzaie D, Marco-Ferreres R, et al. The cis-regulatory dynamics of embryonic development at single-cell resolution. Nature. 2018;555: 538–542.

58. Joshi N, Misharin A. Single-nucleus isolation from frozen human lung tissue for single-nucleus RNA-seq. Available: https://www.protocols.io/view/single-nucleus-isolation-from-frozen-human-lung-ti-zu8f6zw

59. Bolger AM, Lohse M, Usadel B. Trimmomatic: a flexible trimmer for Illumina sequence data. Bioinformatics. 2014;30: 2114–2120.

60. Li H, Handsaker B, Wysoker A, Fennell T, Ruan J, Homer N, et al. The Sequence Alignment/Map format and SAMtools. Bioinformatics. 2009;25: 2078–2079.

61. Li H. Tabix: fast retrieval of sequence features from generic TAB-delimited files. Bioinformatics. 2011;27: 718–719.

62. Li H. Aligning sequence reads, clone sequences and assembly contigs with BWA-MEM. 2013 [cited 23 Jun 2022]. doi:10.48550/arXiv.1303.3997

63. Langmead B, Salzberg SL. Fast gapped-read alignment with Bowtie 2. Nat Methods. 2012;9: 357–359.

64. Wall L, Christiansen T, Orwant J. Programming Perl. “O’Reilly Media, Inc.”; 2000.

65. Zhang Y, Liu T, Meyer CA, Eeckhoute J, Johnson DS, Bernstein BE, et al. Model-based analysis of ChIP-Seq (MACS). Genome Biol. 2008;9: R137.

66. Quinlan AR, Hall IM. BEDTools: a flexible suite of utilities for comparing genomic features. Bioinformatics. 2010;26: 841–842.

67. van Rossum G. Python Reference Manual. 1995.

68. Van Rossum G, Drake FL. Python 3 Reference Manual: (Python Documentation Manual Part 2). CreateSpace; 2009.

69. Dale RK, Pedersen BS, Quinlan AR. Pybedtools: a flexible Python library for manipulating genomic datasets and annotations. Bioinformatics. 2011;27: 3423–3424.

70. R Core Team. R: A Language and Environment for Statistical Computing. 2021. Available: https://www.R-project.org/

71. Melville J. The Uniform Manifold Approximation and Projection (UMAP) Method for Dimensionality Reduction [R package uwot version 0.1.11]. 2021 [cited 23 Jun 2022]. Available: https://CRAN.R-project.org/package=uwot

72. Korsunsky I, Millard N, Fan J, Slowikowski K, Zhang F, Wei K, et al. Fast, sensitive and accurate integration of single-cell data with Harmony. Nat Methods. 2019;16: 1289–1296.

73. Fast Truncated Singular Value Decomposition and Principal Components Analysis for Large Dense and Sparse Matrices [R package irlba version 2.3.5]. 2021 [cited 24 Jun 2022]. Available: https://CRAN.R-project.org/package=irlba

74. Scrucca L, Fop M, Murphy TB, Raftery AE. mclust 5: Clustering, Classification and Density Estimation Using Gaussian Finite Mixture Models. R J. 2016;8: 289–317.

75. Robinson MD, McCarthy DJ, Smyth GK. edgeR: a Bioconductor package for differential expression analysis of digital gene expression data. Bioinformatics. 2010;26: 139–140.

76. Gu Z, Hübschmann D. rGREAT: an R/Bioconductor package for functional enrichment on genomic regions. Bioinformatics. 2022. doi:10.1093/bioinformatics/btac745

77. Tenenbaum D, Maintainer B. KEGGREST: Client-side REST access to the Kyoto Encyclopedia of Genes and Genomes (KEGG). R package version 1.38.0. 2022.

78. Li H. A statistical framework for SNP calling, mutation discovery, association mapping and population genetical parameter estimation from sequencing data. Bioinformatics. 2011. Available: https://academic.oup.com/bioinformatics/article-abstract/27/21/2987/217423

79. McKenna A, Hanna M, Banks E, Sivachenko A, Cibulskis K, Kernytsky A, et al. The Genome Analysis Toolkit: a MapReduce framework for analyzing next-generation DNA sequencing data. Genome Res. 2010;20: 1297–1303.

80. Korhonen J, Martinmäki P, Pizzi C, Rastas P, Ukkonen E. MOODS: fast search for position weight matrix matches in DNA sequences. Bioinformatics. 2009;25: 3181–3182.

81. Wickham H. ggplot2: Elegant Graphics for Data Analysis. Springer Science & Business Media; 2009.

82. Gu Z, Eils R, Schlesner M. Complex heatmaps reveal patterns and correlations in multidimensional genomic data. Bioinformatics. 2016;32: 2847–2849.

83. Sinnamon JR, Torkenczy KA, Linhoff MW, Vitak SA, Mulqueen RM, Pliner HA, et al. The accessible chromatin landscape of the murine hippocampus at single-cell resolution. Genome Res. 2019;29: 857–869.

84. Khan A, Fornes O, Stigliani A, Gheorghe M, Castro-Mondragon JA, van der Lee R, et al. JASPAR 2018: update of the open-access database of transcription factor binding profiles and its web framework. Nucleic Acids Res. 2018;46: D1284.

85. Bakken TE, Jorstad NL, Hu Q, Lake BB, Tian W, Kalmbach BE, et al. Comparative cellular analysis of motor cortex in human, marmoset and mouse. Nature. 2021;598: 111–119.

86. Yao Z, van Velthoven CTJ, Nguyen TN, Goldy J, Sedeno-Cortes AE, Baftizadeh F, et al. A taxonomy of transcriptomic cell types across the isocortex and hippocampal formation. Cell. 2021;184: 3222–3241.e26.

87. Kozareva V, Martin C, Osorno T, Rudolph S, Guo C, Vanderburg C, et al. A transcriptomic atlas of mouse cerebellar cortex comprehensively defines cell types. Nature. 2021;598: 214–219.

88. Amemiya HM, Kundaje A, Boyle AP. The ENCODE Blacklist: Identification of Problematic Regions of the Genome. Sci Rep. 2019;9: 9354.

89. Gencode human reference v39. Available: https://ftp.ebi.ac.uk/pub/databases/gencode/Gencode_human/release_39/gencode.v39.annotation.gtf.gz

90. Gencode mouse reference vM23. Available: http://ftp.ebi.ac.uk/pub/databases/gencode/Gencode_mouse/release_M23/gencode.vM23.annotation.gtf.gz

91. 10X Genomics Build Notes for Reference Packages. Available: https://support.10xgenomics.com/single-cell-gene-expression/software/release-notes/build#hg19_1.2.0

92. Picard toolkit. Broad Institute, GitHub repository. 2019. Available: http://broadinstitute.github.io/picard/

93. Stuart T, Butler A, Hoffman P, Hafemeister C, Papalexi E, Mauck WM, et al. Comprehensive Integration of Single-Cell Data. Cell. 2019. pp. 1888–1902.e21. doi:10.1016/j.cell.2019.05.031

94. Zerbino DR, Johnson N, Juettemann T, Wilder SP, Flicek P. WiggleTools: parallel processing of large collections of genome-wide datasets for visualization and statistical analysis. Bioinformatics. 2014. pp. 1008–1009. doi:10.1093/bioinformatics/btt737

95. Tabula Muris Consortium, Overall coordination, Logistical coordination, Organ collection and processing, Library preparation and sequencing, Computational data analysis, et al. Single-cell transcriptomics of 20 mouse organs creates a Tabula Muris. Nature. 2018;562: 367–372.

96. Granja JM, Corces MR, Pierce SE, Bagdatli ST, Choudhry H, Chang HY, et al. ArchR is a scalable software package for integrative single-cell chromatin accessibility analysis. Nat Genet. 2021;53: 403–411.

97. Benjamini Y, Hochberg Y. Controlling the false discovery rate: A practical and powerful approach to multiple testing. J R Stat Soc. 1995;57: 289–300.

98. Fornes O, Castro-Mondragon JA, Khan A, van der Lee R, Zhang X, Richmond PA, et al. JASPAR 2020: update of the open-access database of transcription factor binding profiles. Nucleic Acids Res. 2020;48: D87–D92.

99. Zhou X, Cain CE, Myrthil M, Lewellen N, Michelini K, Davenport ER, et al. Epigenetic modifications are associated with inter-species gene expression variation in primates. Genome Biol. 2014;15: 547.

